# Bounded by compartment, composition and time: efferocytic transcriptional readouts after ischaemic stroke

**DOI:** 10.64898/2026.11.13.766327

**Authors:** Suping Ding, Yinghao Liu, Yuehuan Zeng, Zhihua Huang, Zhidong Yuan

## Abstract

After ischaemic stroke, efferocytosis is usually inferred from a single bulk-tissue transcript score. Such a score cannot distinguish more phagocytes from more active ones, nor separate clearance from the inflammatory output that follows.

We scored apoptotic demand (DEMAND), clearance machinery (CAPACITY), immunogenic output (OUTCOME) and an inhibitory brake separately across 38 datasets (30 numeric) spanning brain, blood and sorted human parenchyma, rating a program only where 75% of its genes were detected. Each difference was compared with expression-matched random gene sets, checked against leukocyte composition, and reported against the smallest difference its design could resolve. Energy supply and demand were scored on the same scale.

Whole-blood CAPACITY was higher after stroke (+0.613/GSE58294, +0.374/GSE16561), exceeding matched random gene sets. Five lineage proxies left it intact in one cohort (+0.473, 95% CI +0.350 to +0.588) and removed it in a second (+0.014), after age and sex had raised it to +0.488; a third could not be adjusted. Adding the array gradient cut that attenuation to 77.7% (ρ = 0.814), so 97.1% is an upper bound and the cohort difference partly reflects platform. Matched mononuclear cells showed no effect (−0.054; 90% CI −0.218 to +0.110). In rodent brain, supply rose before clearance machinery in three cross-dataset snapshots (oxidative phosphorylation +1.512 in day-3 infiltrating macrophages), with CAPACITY rising and its carrier shifting from microglia at day 3 to macrophages at day 7 — a suggested, not demonstrated order. In the only sorted human parenchyma series, non-microglial myeloid cells made up 37.1% of cells from a CD45-positive sort in stroke donors versus 19.9% in controls (p = 0.057; q = 0.30), and no microglial layer exceeded that series’ resolution limit. Supply and demand ranked samples almost identically, so no transcript-only design can show which sets the limit.

An efferocytic transcript score is bounded by compartment, composition and time: not a flux assay, not a validated stand-alone biomarker in these datasets. Energy supply rises with the work of clearance and can rise in cells that do not engulf, but the two scores move too closely for a transcript-only design to establish which sets the limit.

## Background

Ischaemic stroke leaves a load of dead cells that persists for days. Apoptotic neurons, oligodendrocytes and other parenchymal cells expose phosphatidylserine and release find-me signals, and microglia, border-associated macrophages and infiltrating monocytes are recruited to clear them. Clearance proceeds through recognition, engulfment, phagosome maturation and lysosomal digestion. When this chain fails, undigested DNA and membranes accumulate inside phagocytes, activate cGAS–STING and other innate sensors, and turn a resolving response into a sustained inflammatory one [1–7].

Restoring clearance is therefore an attractive target, and a large literature reports that efferocytosis is suppressed or enhanced after stroke on the basis of transcriptional signatures [8–10]. The inference from a transcript score to a biological function is weaker than it looks. A tissue can score highly for phagocytic genes because it contains more phagocytes, because each phagocyte transcribes the program more strongly, or because a subset of cells with a distinct program has expanded. These alternatives carry different implications for treatment, and a single score over bulk tissue cannot separate them [11,12].

The reason is structural. A bulk score is a function of how many phagocytes a sample contains, how strongly each transcribes the program, and which other cells share the same genes. These three act through the same columns: more cells and more transcript per cell produce the same vector. The separation has to come from outside the score — from samples in which composition was measured, from cells that can be sorted, or from a functional readout taken alongside it. Where none of the three is available, a transcript score has no interpretation.

Three remedies exist and each covers part of the problem. Reference-based deconvolution estimates composition but fails where the platform or tissue has no matching reference, which is the case for most brain series here. Sorted or single-cell profiling removes the composition term and identifies which cell carries a program, but not whether that cell is clearing faster. A functional uptake or degradation assay answers the rate question and says nothing about which transcript sustains it.

Two further problems compound the first. Efferocytosis itself reprograms the phagocyte towards an anti-inflammatory state, so a high clearance score and low inflammation are not independent observations [13]. And stroke is not one tissue at one time: it produces a necrotic core, a penumbra and unaffected tissue, each with a different immune composition [14–17], while blood carries neutrophils that PBMC preparations lack and the peripheral transcriptome varies with sampling design [18]. A signature that rises in one compartment may be flat or reversed in another; treating it as a cross-species biomarker is a category error unless the boundaries have been measured.

We asked four questions: which efferocytic programs change after stroke once machinery is separated from output; in which compartment and at what time; whether a derived balance, θ = DEMAND − CAPACITY, adds information beyond its components; and how much of a tissue-level change reflects composition rather than per-cell capacity. The distinction between how many phagocytes are present and how fast each one clears is old in experimental efferocytosis, where a phagocytic index is counted per cell rather than inferred from a tissue lysate. What has been lacking is a systematic account of where that distinction can be recovered from transcript data and where it cannot. We therefore built a frozen, versioned framework and assembled 38 datasets. Thirty supply numerical results; the other eight were examined and then set aside, each with its reason in Table 1, and two of them nonetheless supply one secondary number each. The result is a map of where the signal can be read, where it cannot, and what would have to be measured next.

**Table 1.**
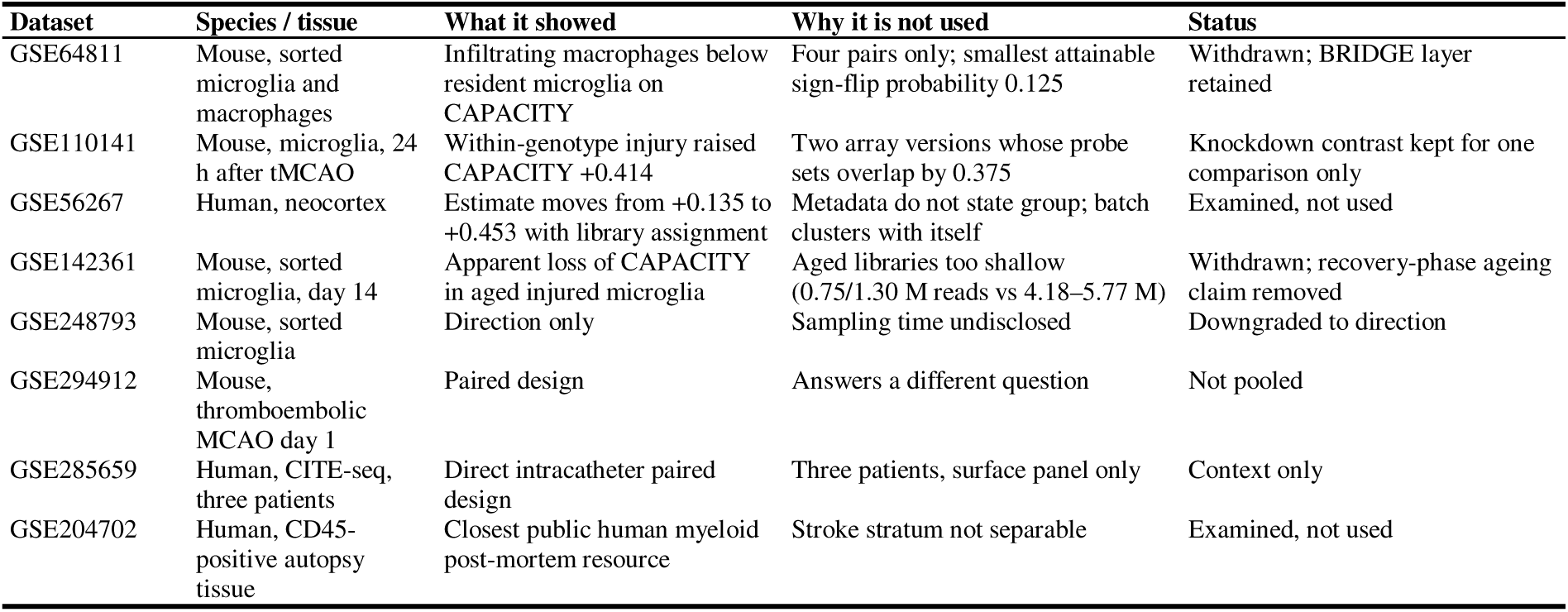
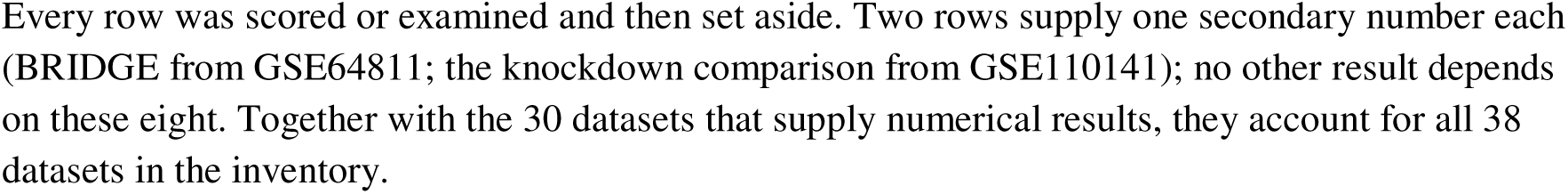
Datasets scored and then withdrawn or downgraded.

## Methods

### Framework construction

Eight programs were assembled from KEGG pathway maps (2026), Gene Ontology annotations and MSigDB v2026.1, together with the published literature [8–10,13] and are set out in Fig. 1. A gene entered a program only when at least two independent sources supported it, and duplicate annotations counted once. DEMAND is the apoptotic-burden program, scored alone and never pooled with the two signal programs beside it, PS_EXPOSURE and FIND_ME. CAPACITY is the union of phosphatidylserine receptors, TAM receptor kinases, engulfment, LC3-associated phagocytosis, phagosome maturation, lysosomal degradation, metabolic processing and six soluble bridging opsonins. OUTCOME was built from KEGG pathways 04620–04623 and GO terms 0002224, 0032481 and 0032731, then stripped of every gene shared with DEMAND or CAPACITY. BRAKE contains six genes: CD300A, CD24, CD47, SIRPA, SIGLEC10 and PECAM1. Five have a documented inhibitory role in engulfment; PECAM1 is expressed on endothelium, platelets and several leukocyte lineages, so its presence in a whole-tissue score may also report vascular or myeloid content. The set was frozen before scoring and we did not remove PECAM1 from it; the consequence for the chronic estimate is stated in the Results. CD300LF was held in a context-dependent category and is not in the six-gene set scored here; adding it back at the day-28 time point gives +1.895 instead of +1.785, so the conclusion does not depend on that choice. INACTIVATION is a two-gene composite (ADAM17 and TIMP3, a shedding enzyme and its inhibitor) analysed as a labelled co-expression quantity, never as net activity. The cGAS–STING cargo set was derived only from genes already in the immunogenic core and scored only in the time-series datasets; its name records the question that motivated it, not a measured cargo flux. No mitochondrial-DNA, TFAM or phagosome-flux assay was performed. A gene had one primary scored program; retained overlaps are recorded in the frozen gene-set record and are not counted twice within a composite. Table 2 records which sets were built and never scored so that names in figures, code and archived gene sets can be reconciled.

**Figure 1.**
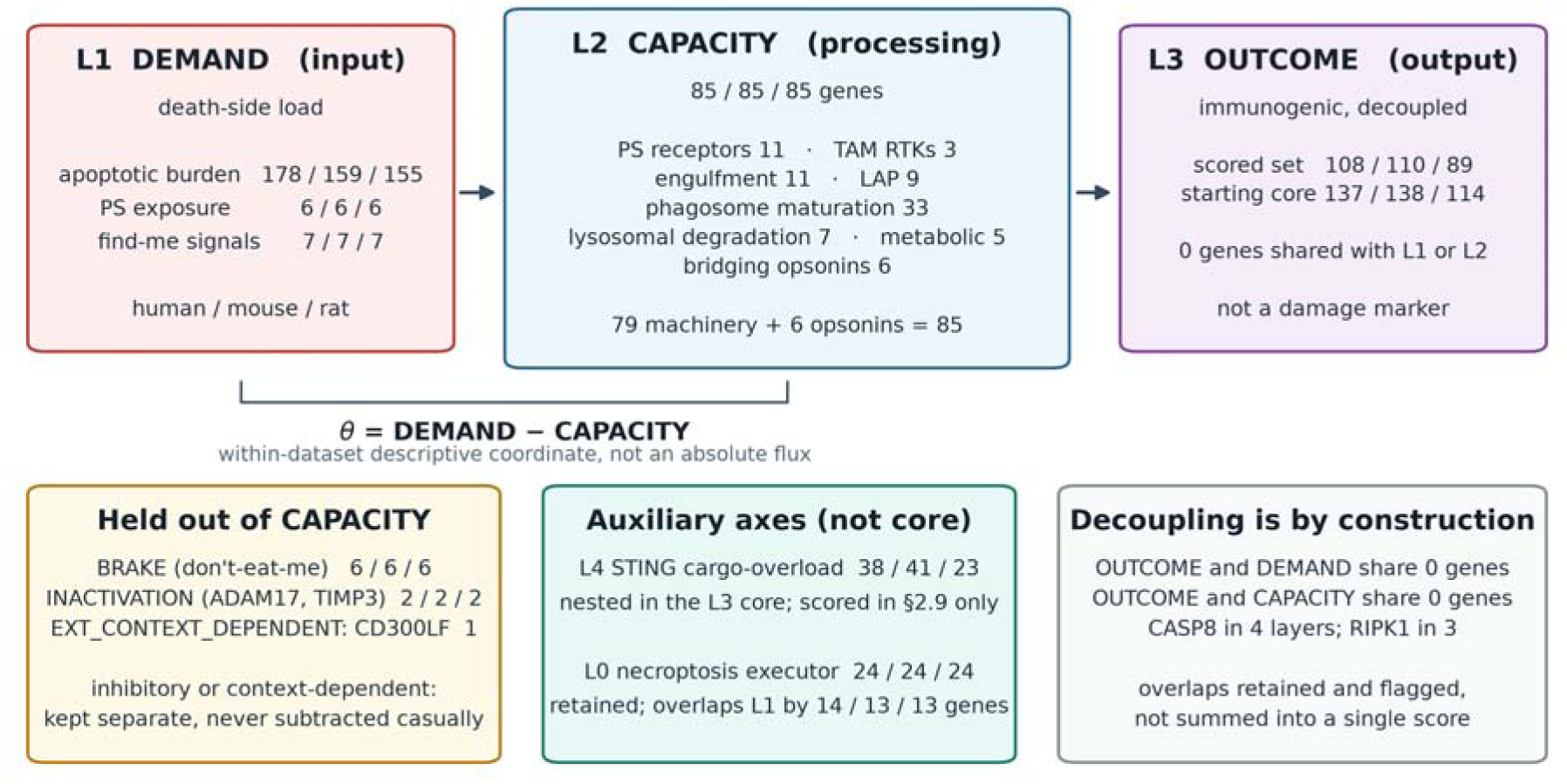
A decoupled three-layer framework. The eight core programs are organised into three layers: a death-side layer (DEMAND, the apoptotic-burden program, with PS_EXPOSURE and FIND_ME scored separately), a processing layer (CAPACITY: receptor, engulfment, phagosome maturation, degradation, metabolic and bridging modules) and an output layer (OUTCOME), with a necroptosis executor and a cGAS–STING extension. The scored OUTCOME set shares no gene with DEMAND or CAPACITY in any species. θ = DEMAND − CAPACITY is a within-dataset difference, not a ratio and not a flux.

**Table 2.** Framework gene-set composition and naming (human / mouse / rat).

| Program (set ID) | Alias / other label in use | Human | Mouse | Rat | Scored | Role |
|---|---|---|---|---|---|---|
| L1_APOPTOTIC_BURDEN | **DEMAND**; θ = DEMAND – CAPACITY | 178 | 159 | 155 | yes | Apoptotic execution and DNA fragmentation |
| L1_PS_EXPOSURE | PS_EXPOSURE | 6 | 6 | 6 | yes | Phosphatidylserine scramblases |
| L1_FIND_ME | FIND_ME | 7 | 7 | 7 | yes | Find-me ligands and receptors |
| L2_CAPACITY | **CAPACITY**; L2_MACHINERY_FULL in older code, identical set | 85 | 85 | 85 | yes | Seven subprograms plus bridging opsonins |
| - L2_PS_RECEPTOR | - | 11 | 11 | 11 | subprogram | Phosphatidylserine receptors, including TREM2/TYROBP |
| - L2_TAM_RTK | - | 3 | 3 | 3 | subprogram | AXL, TYRO3, MERTK |
| - L2_ENGULF | - | 11 | 11 | 11 | subprogram | GULP1/ELMO/DOCK/RAC module and DNM2 |
| - L2_LAP | - | 9 | 9 | 9 | subprogram | LC3-associated phagocytosis |
| - L2_PHAGO_MATUR | - | 33 | 33 | 33 | subprogram | Rab and ARF GTPases, channels, SNAREs |
| - L2_DEGRADATION | - | 7 | 7 | 7 | subprogram | Lysosomal hydrolases and V-ATPase a3 (TCIRG1) |
| - L2_METABOLIC | - | 5 | 5 | 5 | subprogram | Lipid and nuclear-receptor processing |
| L2_DONT_EAT_ME | **BRAKE** | 6 | 6 | 6 | yes | Inhibitory receptors, including CD300A |
| L2_BRIDGE | **BRIDGE**; also counted inside CAPACITY | 6 | 6 | 6 | yes | Opsonins bridging dying cells to phagocytes |
| L2b_INACTIVATION | **INACTIVATION** | 2 | 2 | 2 | yes | ADAM17 and TIMP3; not net activity |
| L3_IMMUNOGENIC_CORE | L3_CORE (majority vote) | 137 | 138 | 114 | no | Starting immunogenic set, before decoupling |
| L3_IMMUNOGENIC_DECOUPLED | **OUTCOME** | 108 | 110 | 89 | yes | Core minus genes shared with DEMAND or CAPACITY |
| L4_STING_CARGO_OVERLOAD | STING (subscore of OUTCOME) | 38 | 41 | 23 | subprogram | cGAS-STING-interferon axis, time series only |
| L0_NECROPTOSIS_EXECUTOR | EXECUTOR | 24 | 24 | 24 | yes | Death-mode comparator, not an efferocytosis program; RIPK1-RIPK3-MLKL axis |

The CAPACITY row is the union of its subprograms. CD300LF is held outside all scored programs. Rows built and never scored (L3_IMMUNOGENIC_CORE_STRICT, L3_IMMUNOGENIC_EXT, L3_IMMUNOGENIC_FULL, L4_PYROPTOSIS, L2_EAT_ME, L2_MACHINERY_FULL) appear because the code and archived gene sets carry them. L0_NECROPTOSIS_EXECUTOR is carried as a death-mode comparator rather than as a component of the efferocytosis framework: it scores the execution machinery of necroptosis, shares no gene with CAPACITY in any species, and therefore reports cell death rather than clearance.

Coverage was computed per dataset as detected program genes divided by program size. A gene counts as measurable only if it clears three filters: it is represented on the platform or annotation; it maps to an unambiguous one-to-one orthologue where species differ; and it survives the expression filter applied to that matrix. Coverage therefore counts presence, not strength — a gene at the detection limit counts as present, because a flat gene contributes almost nothing to a mean of standardised values, whereas removing it by name changes what the program is. A program below 75% coverage, or below five genes, was dropped from that dataset rather than scored on a partial set. Split-half reliability was estimated by splitting each program into two halves 200 times and correlating the half-scores across samples; medians and 95% intervals are reported with Spearman–Brown correction. This estimates internal consistency, not test–retest reliability. For the rat cell-type comparison, each delta was refitted with linear and quadratic terms for the number of detected genes per nucleus, and raw and adjusted estimates are reported side by side. Full membership and the source map are in Supplementary Table S1; file-level checksums are in MANIFEST.md5 inside the Zenodo archive.

### Datasets

We assembled 38 datasets: 36 public accessions and two in-house cohorts. Thirty supply a number in the results; eight public accessions were scored or examined and then set aside (Table 1). The inventory is generated from the analysis ledger rather than typed, and all 38 entries carry accession, species, tissue, assay, sample counts, role and disposition (Supplementary Table S2).

Human whole blood: GSE16561 (39 stroke, 24 control; GPL6883), GSE58294 (23 patients sampled at 3, 5 and 24 h, 23 controls; analysed at the subject level with one mean per patient) and GSE202518 (six severe [NIHSS ≥ 8], six moderate [NIHSS < 8] and four elderly controls reanalysed from a separate study, so its contrast is directional). Human PBMC: GSE22255 (20 stroke, 20 control, matched for age and sex). Human brain: GSE162955 (paired infarct core and contralateral cortex from six post-mortem donors). A second neocortex series, GSE56267, could not be used (Table 1). A third series, GSE331114, sorts CD45-positive cells from fresh parenchyma: four stroke and four control donors, 70,773 cells after quality control, median depth 1,452 to 7,993 unique molecules per cell. Its role is composition and single-transcript detection; its layer scores are reported against an explicit resolution limit and none carries a capacity claim.

For GSE162955 the deposited matrix orders columns as six contralateral and then six core samples while the sample titles are interleaved, so tissue assignment follows column order. It was verified against XIST and seven Y-chromosome genes in 12 of 12 samples, against the original study’s group means (r = +0.985 over 6,061 genes), and by re-deriving its differential-expression list (2,562 of 2,716 genes recovered). Assignment was fixed from the raw CEL files by RMA re-processing and maximum correlation (r = 1.0000). Every paired statistic is also reported over the 48 sex-compatible pairings and all 720 within-block permutations.

Mouse brain: GSE23160 (MCAO 2 h, reperfusion at 2, 8 and 24 h; n = 4 per group), GSE233815 (Visium, five sections), GSE171169 (CD45-high cells at 5 and 14 days), GSE227651 (whole brain at sham, 1, 3 and 7 days), GSE154396 (Cd300a^fl/fl × LysM-Cre, 0, 1 and 3 h), GSE332910 (striatal snRNA, sham, 0, 24 h, 7 and 14 days; n = 3), GSE137482 (3 and 18 months, sham versus pMCAO day 3; n = 6 per cell), GSE137595 (24 arrays, 2 h after 90 min MCAO), GSE174574 and GSE319238 (integrated with GSE227651; 110,664 cells), GSE318800 (Htr2b cKO) and GSE337578 (SeekSpace, day 3). Rat: GSE250245 (sMCAO, mMCAO and sham; six libraries, 73,816 nuclei), GSE23651, GSE37587 (68 paired whole-blood samples from 34 patients) and GSE345807 (day-21 scRNA). GSE163654 (paired mouse–rat orthologues) was used for rank-order comparison only, and GSE267023 (PBMC single-cell) supplied design-limited lineage context. Neither enters a primary claim.

In-house: a mouse P1/P2 tMCAO series at day 1 (21 libraries, two excluded after batch and PCA review, leaving 19) and a rat tMCAO 28-day cohort (Sham 7, Model 3, ICT 5, ICTF 5; 26 animals, six PCA outliers removed, leaving 20). Both exclusions were made after inspecting the data rather than under a rule fixed in advance, and the criteria are recorded in the inventory.

Proteomics, which are not among the 38: PXD012151 (rat cortex at 1, 7 and 14 days; 8,175 protein groups) [19], PXD055588 (mouse microglia at 24 h; 9,785 groups) [20] and PXD025077 (mouse cortex, TMT 10-plex at 1, 3, 7, 14 and 28 days; 8,068 proteins) [21]. These form a layer check and none is a scored dataset. PXD022850 and PXD016538 provide provenance for GSE162955 and GSE137595 respectively and are not scored either.

### Scoring

Bulk and microarray matrices were processed from log2(CPM+1) or from deposited normalised values. GSE23160 and GSE137595 were log2-transformed and quantile-normalised across samples, and in-house counts were transformed with DESeq2 VST. For each dataset and region, each gene was z-scored across samples or cells, and a program score was the unweighted mean of the available gene z-scores. Scores were not z-scored a second time. A positive value means above the dataset average, not an absolute biological zero, so cross-dataset statements are restricted to direction and Cliff’s δ. In GSE225948, cells were normalised to counts per 10,000, log1p-transformed and averaged within each parent compartment and animal before scoring.

Where a gene was represented by multiple probes, its probes were averaged to a single gene-level value; the sum, the median and the single brightest probe were recomputed as a sensitivity analysis (Table S14). Cross-species comparisons used unambiguous 1:1 orthologues. The output layer was also scored against four Hallmark sets on the same samples with identical scoring, to report how far its behaviour departs from a standard inflammatory index. Protein series were handled as a separate layer and not scored. Only one of the three meets the 75% coverage requirement for any program (PXD055588, 73/85 for CAPACITY) and it deposits no per-sample values, so no protein layer score exists; the other two reach 42/85 and 55/85 and supply directional readouts only. Band intensities were summed per protein group, log2-transformed after adding half the smallest positive value, and corrected by subtracting each sample’s whole-proteome median. Layer differences were tested against a competitive permutation null over proteins (5,000 draws).

### Composition adjustment and decomposition

Marker-based proxies were built for five leukocyte lineages after removing framework genes, leaving 19, 6, 12, 7 and 6 genes for neutrophil, monocyte, lymphoid, erythroid and platelet proxies. Each proxy is a z-scored marker set, not an estimated fraction; reference-based deconvolution was not applied because no platform-matched brain signature matrix is available for the whole-blood arrays used here. The five entered jointly as covariates in a linear model that already contained stroke status and, where available, age and sex. Collinearity was checked by variance inflation factor (1.64–3.67).

We tested whether the attenuation was specific to the proxies rather than the cost of adding five covariates, by permuting the proxy block 1,000 times under two nulls: each column permuted independently (preserving its marginal distribution but breaking correlation among proxies), and all five columns permuted together (preserving their joint geometry but breaking their association with CAPACITY). In both cohorts the observed attenuation exceeded every permutation (p = 0.001, the resolution limit of 1,000 draws). The 95th percentile of the null was 7.3% in GSE58294 and 13.4% in GSE16561, against observed attenuation of 22.8% and 97.1%.

A second control asked whether the covariate block moves programs unrelated to leukocytes. Two hundred gene sets were drawn from genes least related to any proxy (maximum absolute correlation below 0.15) and matched to CAPACITY in size; 200 further sets were drawn from all detected non-framework genes; and a third block of 200 was matched one-for-one to the CAPACITY genes on intensity percentile. Each was taken through the identical model, and results are reported as the change in the stroke coefficient.

When the covariates reconstruct the grouping, the adjusted coefficient is a residual of a fit that has already absorbed the group label; we report it with its bootstrap interval and call it unidentified rather than a reversal. In GSE16561, GSE58294 and GSE22255 the proxies recover R² = 0.469, 0.319 and 0.112, so the adjusted values are estimates; in three contrasts of GSE202518 they recover 0.875, 0.920 and 0.963, so there the adjusted values carry no group information.

Oaxaca–Blinder decomposition partitioned the CAPACITY gap into a composition term and a within-cell term under three or four reference bases — three for the rat cell-type gap and four for the human whole-blood gap [22,23]. Bootstrap resampling was stratified by library for the rat data and by group for the blood data, with 1,000 iterations. The human decomposition partitions the raw case–control gap and is therefore not the same estimand as the coefficient attenuation in Table 3; the two are reported side by side.

**Table 3.**
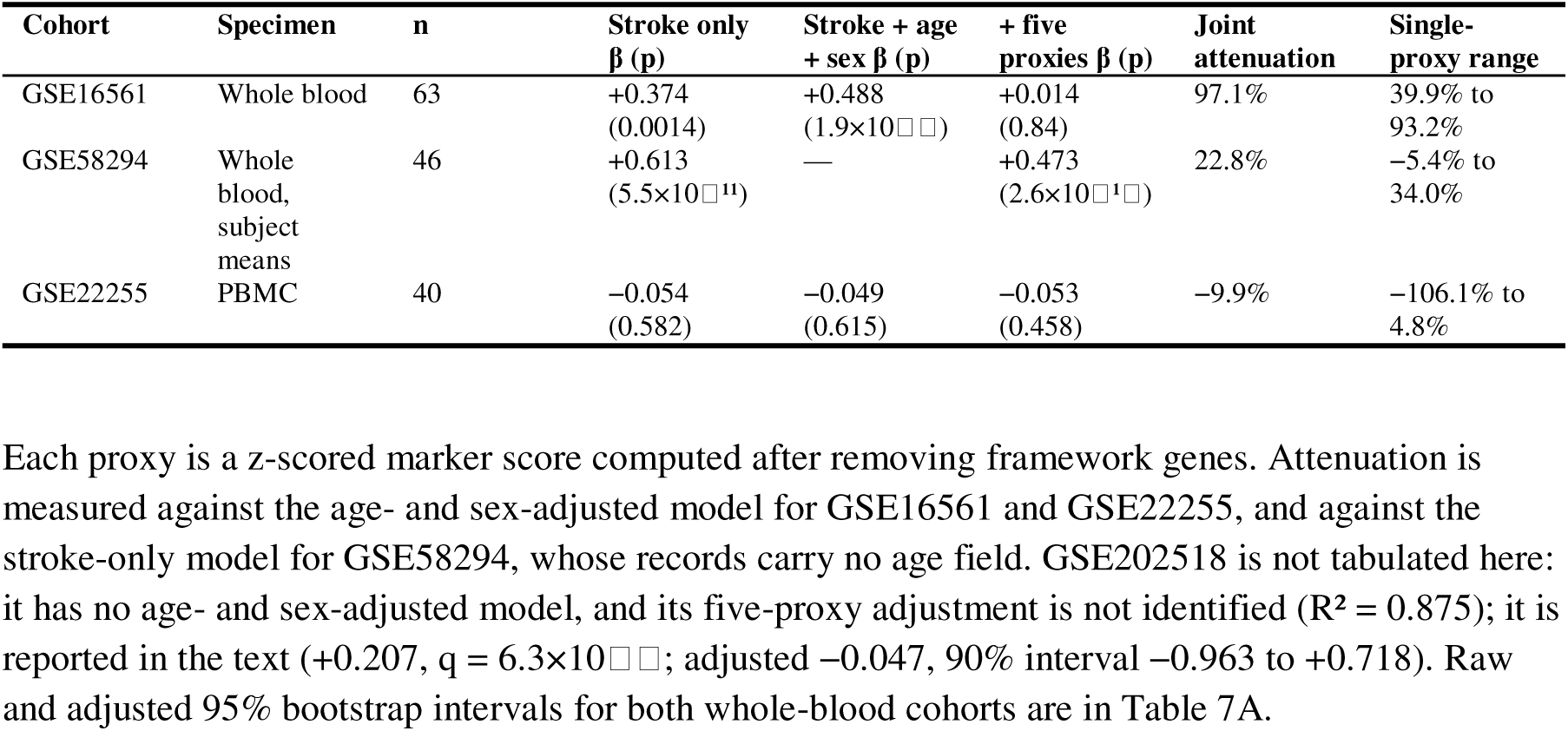
Composition adjustment of the CAPACITY effect in two whole-blood cohorts and one PBMC cohort.

| Cohort | Specimen | n | Stroke only β (p) | Stroke + age + sex β (p) | + five proxies β (p) | Joint attenuation | Single-proxy range |
|---|---|---|---|---|---|---|---|
| GSE16561 | Whole blood | 63 | +0.374 (0.0014) | +0.488 (1.9×10□□) | +0.014 (0.84) | 97.1% | 39.9% to 93.2% |
| GSE58294 | Whole blood, subject means | 46 | +0.613 (5.5×10□¹¹) | — | +0.473 (2.6×10□¹□) | 22.8% | −5.4% to 34.0% |
| GSE22255 | PBMC | 40 | −0.054 (0.582) | −0.049 (0.615) | −0.053 (0.458) | −9.9% | −106.1% to 4.8% |

Each proxy is a z-scored marker score computed after removing framework genes. Attenuation is measured against the age- and sex-adjusted model for GSE16561 and GSE22255, and against the stroke-only model for GSE58294, whose records carry no age field. GSE202518 is not tabulated here: it has no age- and sex-adjusted model, and its five-proxy adjustment is not identified (R² = 0.875); it is reported in the text (+0.207, q = 6.3×10□□; adjusted −0.047, 90% interval −0.963 to +0.718). Raw and adjusted 95% bootstrap intervals for both whole-blood cohorts are in Table 7A.

### Statistical inference

Continuous comparisons used Welch’s t-test; effect sizes are Cliff’s δ. Association p-values came from permutation tests (two-sided for signed group contrasts, one-sided upper-tail for gene-set controls), with 1,500 draws for GSE23160, 999 torus shifts for Visium spatial autocorrelation (Moran’s I [24]), 2,000 draws for expression-matched gene-set controls, 2,000 draws for the detection-matched nulls and 1,000 stratified bootstraps for decomposition. A permutation p cannot be smaller than one over its number of draws plus one, so results are reported as p below that bound, never as p = 0. Two tail conventions appear in this paper: where the count includes only draws more extreme than the observed value the bound is 1/(B + 1), and where a two-sided test of a signed difference also counts the observed value it is 2/(B + 1). At 10,000 draws these are 1×10 and 2×10, and both appear here as the smallest value their test can return. With fewer than five animals per group, p-values appear only where an exact permutation null existed, and effect sizes are primary.

Expression-matched controls were 2,000 non-framework gene sets of the same size and expression-decile profile as the tested program; the detection limits of the sorted-cell series used the same construction with 10,000 draws, the number stated again in the legend of Table 4. Detection-matched nulls additionally matched on detection depth as well as expression under six schemes, of which the two set-level schemes were calibrated against the non-replicating cohort and adopted (Table S16). The 95th percentile of |Δ| across the expression-matched sets is the smallest difference a dataset can distinguish from an arbitrary set of equally expressed genes; a program below it is reported as below the resolution of the design rather than as unchanged. Because the same draws set both the percentile and the p-value, exceeding that threshold and returning p < 0.05 are the same event. This threshold is a nominal 5% screen applied before any correction, and it concerns the expression-matched null only.

**Table 4.**
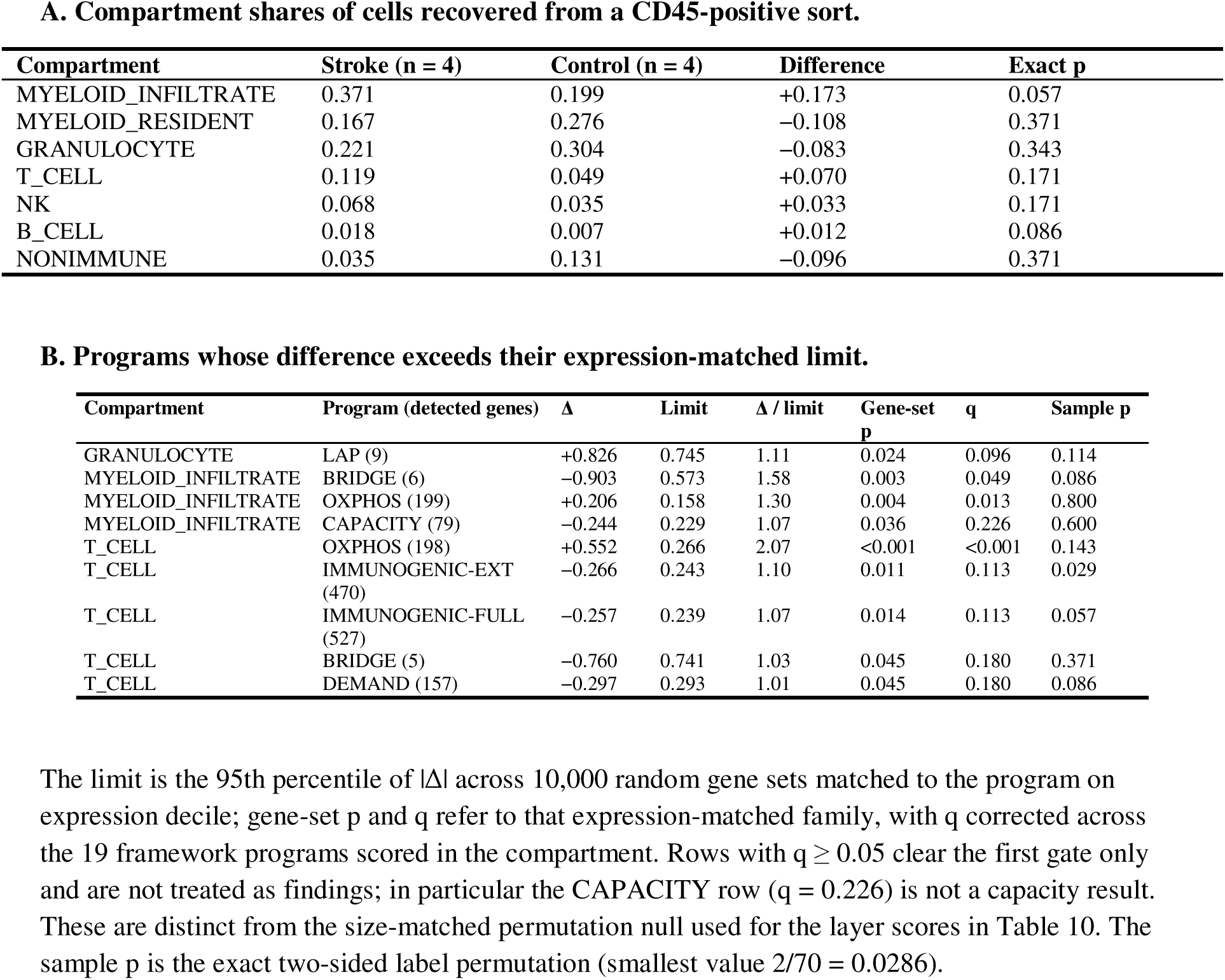
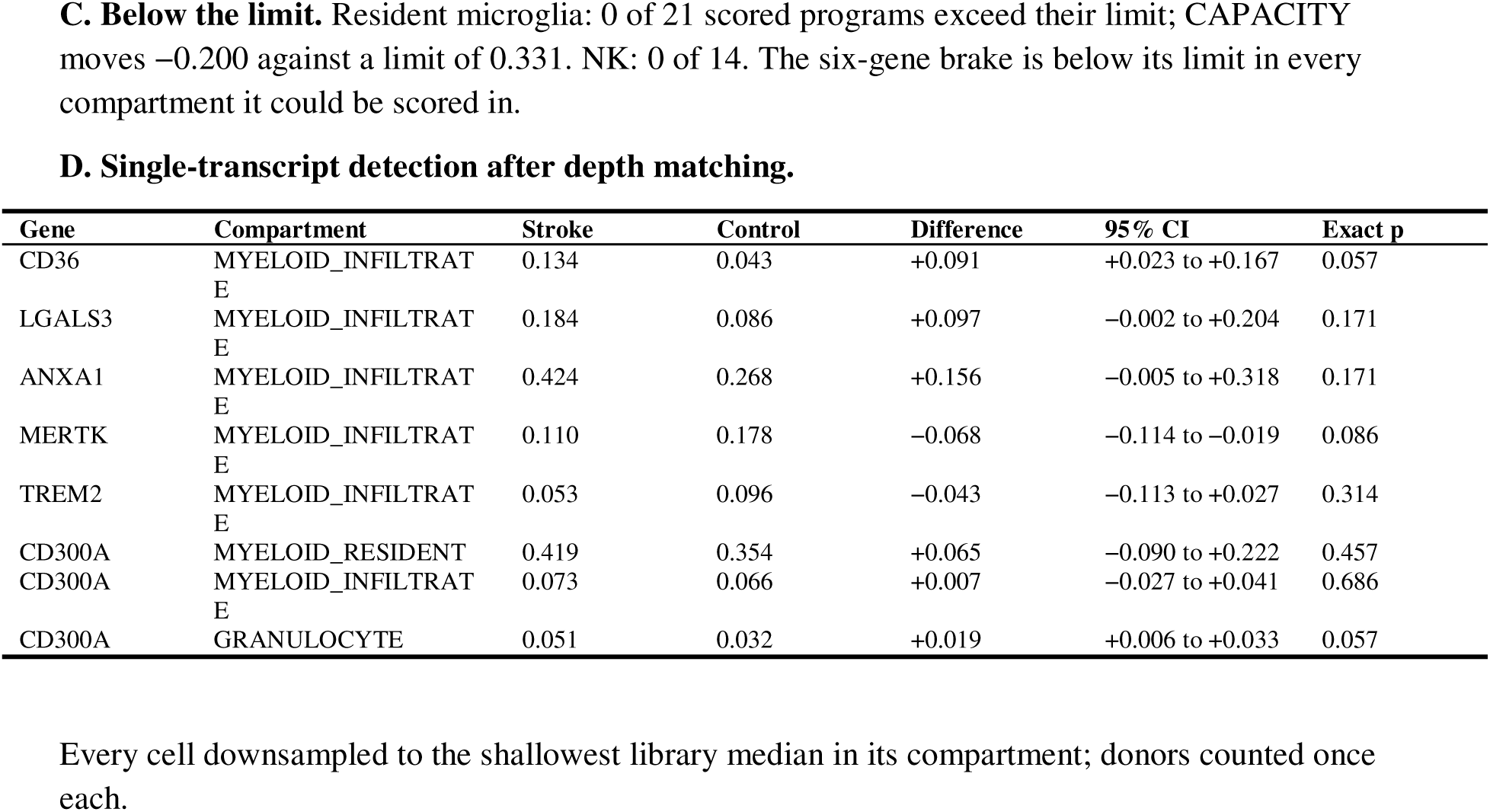
Human brain parenchyma after stroke (GSE331114).

| **A. Compartment shares of cells recovered from a CD45-positive sort.** |  |  |  |  |  |  |  |
|---|---|---|---|---|---|---|---|
| **Compartment** | **Stroke (n = 4)** | **Control (n = 4)** | **Difference** | **Exact p** |  |  |  |
| MYELOID_INFILTRATE | 0.371 | 0.199 | +0.173 | 0.057 |  |  |  |
| MYELOID_RESIDENT | 0.167 | 0.276 | −0.108 | 0.371 |  |  |  |
| GRANULOCYTE | 0.221 | 0.304 | −0.083 | 0.343 |  |  |  |
| T_CELL | 0.119 | 0.049 | +0.070 | 0.171 |  |  |  |
| NK | 0.068 | 0.035 | +0.033 | 0.171 |  |  |  |
| B_CELL | 0.018 | 0.007 | +0.012 | 0.086 |  |  |  |
| NONIMMUNE | 0.035 | 0.131 | −0.096 | 0.371 |  |  |  |
| **B. Programs whose difference exceeds their expression-matched limit.** |  |  |  |  |  |  |  |
| **Compartment** | **Program (detected genes)** | **Δ** | **Limit** | **Δ / limit** | **Gene-set p** | **q** | **Sample p** |
| GRANULOCYTE | LAP (9) | +0.826 | 0.745 | 1.11 | 0.024 | 0.096 | 0.114 |
| MYELOID_INFILTRATE | BRIDGE (6) | −0.903 | 0.573 | 1.58 | 0.003 | 0.049 | 0.086 |
| MYELOID_INFILTRATE | OXPHOS (199) | +0.206 | 0.158 | 1.30 | 0.004 | 0.013 | 0.800 |
| MYELOID_INFILTRATE | CAPACITY (79) | −0.244 | 0.229 | 1.07 | 0.036 | 0.226 | 0.600 |
| T_CELL | OXPHOS (198) | +0.552 | 0.266 | 2.07 | <0.001 | <0.001 | 0.143 |
| T_CELL | IMMUNOGENIC-EXT (470) | −0.266 | 0.243 | 1.10 | 0.011 | 0.113 | 0.029 |
| T_CELL | IMMUNOGENIC-FULL (527) | −0.257 | 0.239 | 1.07 | 0.014 | 0.113 | 0.057 |
| T_CELL | BRIDGE (5) | −0.760 | 0.741 | 1.03 | 0.045 | 0.180 | 0.371 |
| T_CELL | DEMAND (157) | −0.297 | 0.293 | 1.01 | 0.045 | 0.180 | 0.086 |
| The limit is the 95th percentile of \|Δ\| across 10,000 random gene sets matched to the program on expression decile; gene-set p and q refer to that expression-matched family, with q corrected across the 19 framework programs scored in the compartment. Rows with q ≥ 0.05 clear the first gate only and are not treated as findings; in particular the CAPACITY row (q = 0.226) is not a capacity result. These are distinct from the size-matched permutation null used for the layer scores in Table 10. The sample p is the exact two-sided label permutation (smallest value 2/70 = 0.0286). |  |  |  |  |  |  |  |
| **C. Below the limit.** Resident microglia: 0 of 21 scored programs exceed their limit; CAPACITY moves −0.200 against a limit of 0.331. NK: 0 of 14. The six-gene brake is below its limit in every compartment it could be scored in. |  |  |  |  |  |  |  |
| **D. Single-transcript detection after depth matching.** |  |  |  |  |  |  |  |
| **Gene** | **Compartment** | **Stroke** | **Control** | **Difference** | **95% CI** | **Exact p** |  |
| CD36 | MYELOID_INFILTRATE | 0.134 | 0.043 | +0.091 | +0.023 to +0.167 | 0.057 |  |
| LGALS3 | MYELOID_INFILTRATE | 0.184 | 0.086 | +0.097 | −0.002 to +0.204 | 0.171 |  |
| ANXA1 | MYELOID_INFILTRATE | 0.424 | 0.268 | +0.156 | −0.005 to +0.318 | 0.171 |  |
| MERTK | MYELOID_INFILTRATE | 0.110 | 0.178 | −0.068 | −0.114 to −0.019 | 0.086 |  |
| TREM2 | MYELOID_INFILTRATE | 0.053 | 0.096 | −0.043 | −0.113 to +0.027 | 0.314 |  |
| CD300A | MYELOID_RESIDENT | 0.419 | 0.354 | +0.065 | −0.090 to +0.222 | 0.457 |  |
| CD300A | MYELOID_INFILTRATE | 0.073 | 0.066 | +0.007 | −0.027 to +0.041 | 0.686 |  |
| CD300A | GRANULOCYTE | 0.051 | 0.032 | +0.019 | +0.006 to +0.033 | 0.057 |  |

Every cell downsampled to the shallowest library median in its compartment; donors counted once each.

Multiple testing was corrected separately by Benjamini–Hochberg on two-sided p-values from a different, size-matched null of 10,000 draws, whose random stream was derived from a hash of dataset, contrast and program so that a program’s null does not depend on how many others were computed. Requiring both gates is more conservative than either alone, and the two draw sets are independent. A corrected q is defined within the set of tests corrected together, so q values from different sets are not compared; each table legend states the family it used. Cross-dataset statements are counts, not tests. When all 183 arms are placed in one Benjamini–Hochberg family, 37 of the 38 family-internal results survive.

Expression-matched sets do not preserve co-expression: genes are drawn independently within each decile, so the spread of a module mean is narrower than one in which genes rise and fall together. Permuting sample labels keeps that covariance and asks whether the group difference survives relabelling, at a resolution fixed by the design (the smallest two-sided value is 2/C(n_a+n_b, n_a), or 0.10 for three against three and 0.0286 for four against four). Where the control group holds twenty or more samples the two approaches agree. We use the two-step rule described above rather than requiring all three nulls to agree, because a label permutation cannot return a value below 2/C(n_a+n_b, n_a); requiring it would settle the smallest designs by arithmetic instead of by data.

### The energy axis

Because clearance consumes ATP, we scored the energy a program needs on the same standardised scale as the machinery, so a difference in one dataset is commensurable with a difference in another. The axis covered eighteen contrasts across six datasets: the rat time series, the three whole-blood cohorts, the PBMC cohort and the sorted human parenchyma. A contrast was eligible when it compared two groups explicitly and held at least four libraries in each. Thirty datasets were assessed; eighteen contrasts, in six of them, met both conditions. Eligible contrasts that were not run, and the reason in each case, are listed by dataset in Supplementary Table S17.

Supply was measured with three external Hallmark sets — oxidative phosphorylation, glycolysis and hypoxia — each of which shares exactly one gene with CAPACITY (MERTK, TCIRG1 and SRPX for glycolysis, oxidative phosphorylation and hypoxia respectively); these genes were retained because the overlap was far below the prespecified downgrade threshold, so the axes are external but not gene-disjoint. Demand was measured with three Reactome sets chosen for nucleotide-hydrolysing work before the results were inspected: the phagocyte respiratory burst (NADPH oxidase and the V-ATPase proton pump), Fcγ-receptor-driven phagocytosis (the actin machinery of cup closure), and lysosomal vesicle biogenesis. Each group of three formed its own Benjamini–Hochberg set and the two groups were never pooled. A fourth group repeated actin and scission genes taken from CAPACITY itself; it restates genes already in the framework and is not an independent measurement.

A difference was reported only when it exceeded the 95th percentile of the expression- and size-matched null for that dataset and then survived correction within its own group; where only the first held, that is stated. Because the three demand sets do not measure one quantity, each contrast was also classified by how many of them exceeded the threshold and whether they agreed in sign. The supply– demand difference was tested by permuting group labels. Since a z-scored score carries the sample’s global expression axis, the correlation between the two scores was recomputed after subtracting each sample’s mean z over the scoring universe, and both values are reported. Composition adjustment of these new comparisons reused the marker proxies and the cleaning rule of the main analysis.

Human whole-blood analyses used the subject level. For GSE58294 the within-patient ICC of CAPACITY was 0.376, giving a design effect of 1.75 and an effective sample size of about 39 rather than 69. Nested models compared OUTCOME ∼ DEMAND with OUTCOME ∼ DEMAND + CAPACITY; incremental F and ΔR² are reported, with AIC alongside its small-sample caveat. θ = DEMAND − CAPACITY was analysed as a descriptive difference, not a ratio: centred scores have positive and negative values, so a ratio explodes near zero and carries no stable meaning. For PBMC equivalence, two one-sided tests used a 0.3-SD margin.

For sorted populations with three or four libraries per group, each program × contrast carries a detection limit — the minimum detectable standardised difference at α = 0.05 two-sided and 80% power (|d| = 3.071 at 3 versus 3; 2.381 at 4 versus 4). A non-significant contrast below that limit is labelled uninformative, not negative.

Blood-effect uncertainty came from 2,000 subject-or sample-level bootstrap refits. Gene-set dependence was tested by dropping each CAPACITY gene in turn, by 70% random subsets, and by scoring the eight modules separately. Pseudobulk scores averaged nucleus scores within library and cell type, restricted to strata with at least 30 nuclei. Cross-cohort synthesis used Hedges’ g and a DerSimonian–Laird model, reported for heterogeneity only.

### Gene-set provenance

The paired human transcriptomes (GSE162955) are accompanied by a label-free proteome (PXD022850) [25], used only to corroborate tissue assignment. Gene membership is documented in Supplementary Table S1; for the inhibitory brake it follows the primary receptor literature — the CD300 family [26], CD300A binding to phosphatidylserine and phosphatidylethanolamine on dying cells [27], CD47 in microglial pruning [28], and the CD47–SIRPα interaction [29] — and none of those reports supplies an effect size used here.

### Data, code and reporting

Datasets are listed in Supplementary Table S2 with accession, platform, sample counts, role and disposition. Code, gene sets, coverage tables and the dataset manifest are archived in Zenodo; the build is frozen at 2026-10-21 and the analysis at 2026-10-22, with the source map in Supplementary Table S1 and file-level checksums in MANIFEST.md5 inside that archive. The analysis was not pre-registered (individual analyses were pre-specified in locked scripts, which is distinct from registration): the framework, gene sets and decision rules were fixed before the datasets added in later versions were scored, and each of those is marked as such in the inventory, so a reader can tell which results were produced under the rules as first written and which were produced later by the same rules. The two in-house series have no public accession at present and can be made available to reviewers on request; they will be deposited in GEO with raw counts and metadata on acceptance. The analysis archive has DOI 10.5281/zenodo.24415702 (version 1.0.1).

Human analyses use de-identified public data; no new human subjects were recruited. In-house animal samples were generated under institutionally approved protocols. ARRIVE 2.0 reporting guidance does not apply here because no new animal experiments were performed for this manuscript.

### Use of language models in manuscript preparation

An AI coding assistant was used to write and check analysis scripts, to regenerate figures and tables from the tabulated results, and to help reorganise and correct manuscript text. It did not perform any statistical analysis on its own initiative, and it is not an author: every number reported here was checked by the authors against the result files it was generated from. No language model satisfied the journal’s authorship criteria.

## Results

### Whole blood carries a real signal that is mostly composition

In two whole-blood cohorts, CAPACITY was higher after stroke than in controls. In GSE58294, analysed at the subject level because 69 samples came from 23 patients, the difference was +0.613 (95% CI +0.480 to +0.752; Cliff’s δ = 0.970; subject-level Mann–Whitney p = 1.9×10). In GSE16561 it was +0.374 (95% CI +0.157 to +0.584; δ = 0.483). Both exceeded 2,000 expression-matched, non-framework gene sets of the same size and profile (p ≤ 5.0×10, the smallest value 2,000 draws can return; z = 5.48 and 5.60), so the signal is not an artefact of gene number or expression level. Matching the null on detection depth as well as expression, using the set-level scheme calibrated so that the non-replicating PBMC cohort stays non-significant, returns the same bound in both replication cohorts (Table S16), so sequencing depth does not explain it either (Fig. 2A, D).

**Figure 2.**
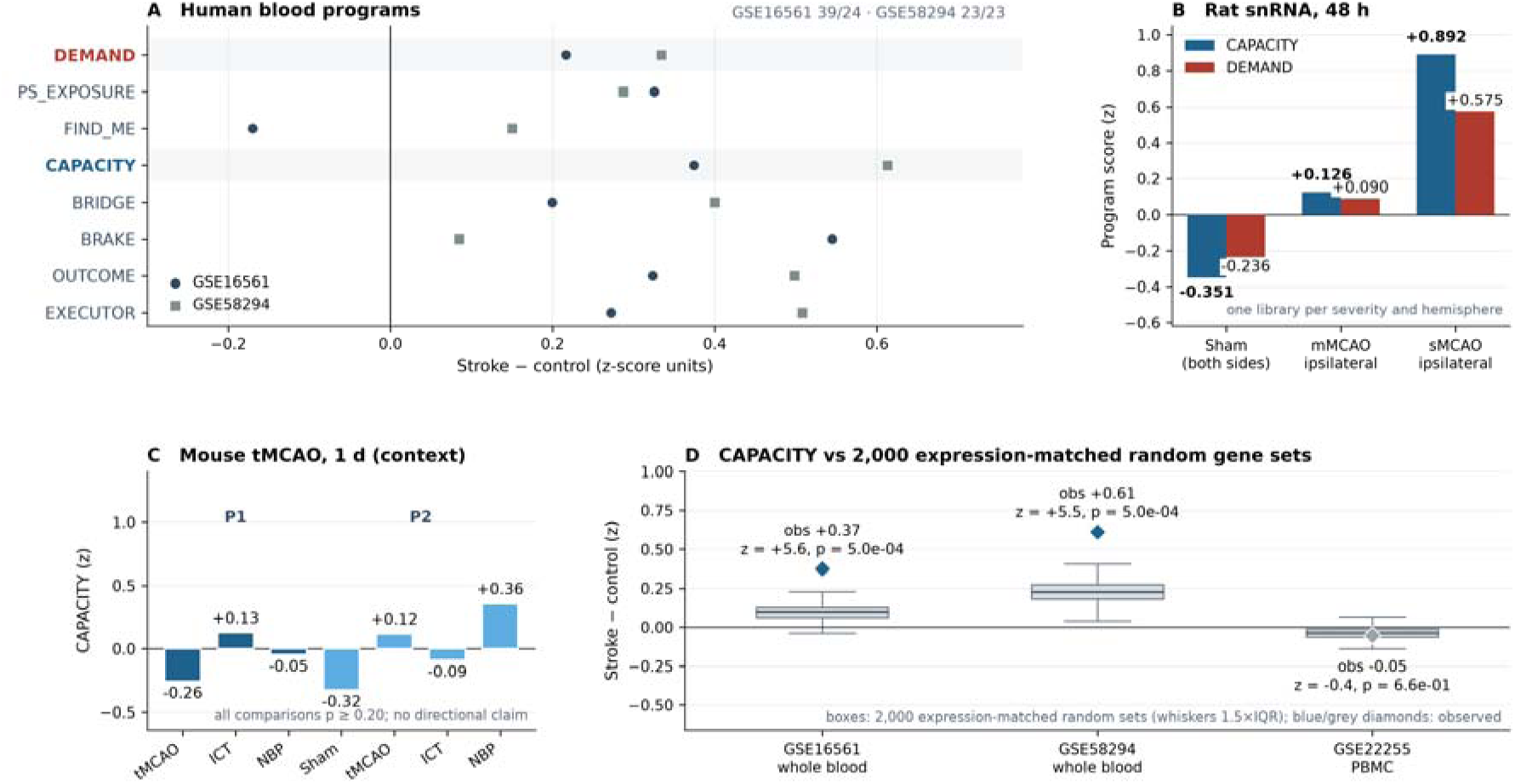
CAPACITY across compartments and against expression-matched controls. (A) Stroke minus control differences in program scores in GSE16561 (39/24) and GSE58294 (23/23), with CAPACITY and DEMAND highlighted. (B) Rat sMCAO single nuclei: CAPACITY rises with severity and reaches +0.892 in severe ipsilateral cortex. (C) Mouse one-day tMCAO context series: the two in-house batches point in opposite directions, all comparisons p ≥ 0.20; this series supplies context only and does not enter a primary claim. (D) Observed CAPACITY against 2,000 expression-matched, non-framework gene sets in both whole-blood cohorts and in PBMCs.

The signal is not specific to CAPACITY. Of 371 framework genes detected in both cohorts, 281 changed in the same direction after stroke (75.7% against a permutation null of 64.0%, z = 6.2). Within CAPACITY, 68 of 83 shared genes moved in the same direction (81.9%; ρ = 0.295, p = 0.0068), higher than the 74.0% for the remaining framework genes but not significantly so (Fisher two-sided p = 0.15). The blood signal is a coordinated myeloid and inflammatory response, not a clearance-specific one.

Composition explains much of the blood signal, but how much depends on the cohort and on which covariates are in the model. In GSE16561 the stroke group was 13.1 years older than controls; adjusting for age and sex raised the CAPACITY coefficient from +0.374 to +0.488, so age was masking part of the effect (Table 5). Adding the five lineage proxies then gave +0.014 (95% CI −0.148 to +0.150), an attenuation of 97.1% (Table 3). In GSE58294 the same proxies removed 22.8% (+0.613 to +0.473; 95% CI +0.350 to +0.588). In GSE202518 the unadjusted direction was reproduced (+0.207, q = 6.3×10□□), but the proxies predict group with R² = 0.875, so the adjusted value (−0.047, interval −0.963 to +0.718) is unresolved. One further comparison changes how these numbers should be read. In GSE16561 CAPACITY correlates with the arrays’ global intensity gradient at ρ = 0.814; adding the gradient to the proxy model moves the coefficient to +0.109 (p = 0.043) and the attenuation to 77.7%. Without age and sex in the baseline the corresponding figures are 82.4% and 65.5%. In GSE58294 that correlation is ρ = −0.008 and the joint model barely moves (26.68% to 26.10%). The two covariate blocks therefore cannot be separated in the cohort carrying the largest attenuation. We report 97.1% as the upper bound of a model-dependent range, not as a share attributable to composition.

**Table 5.**
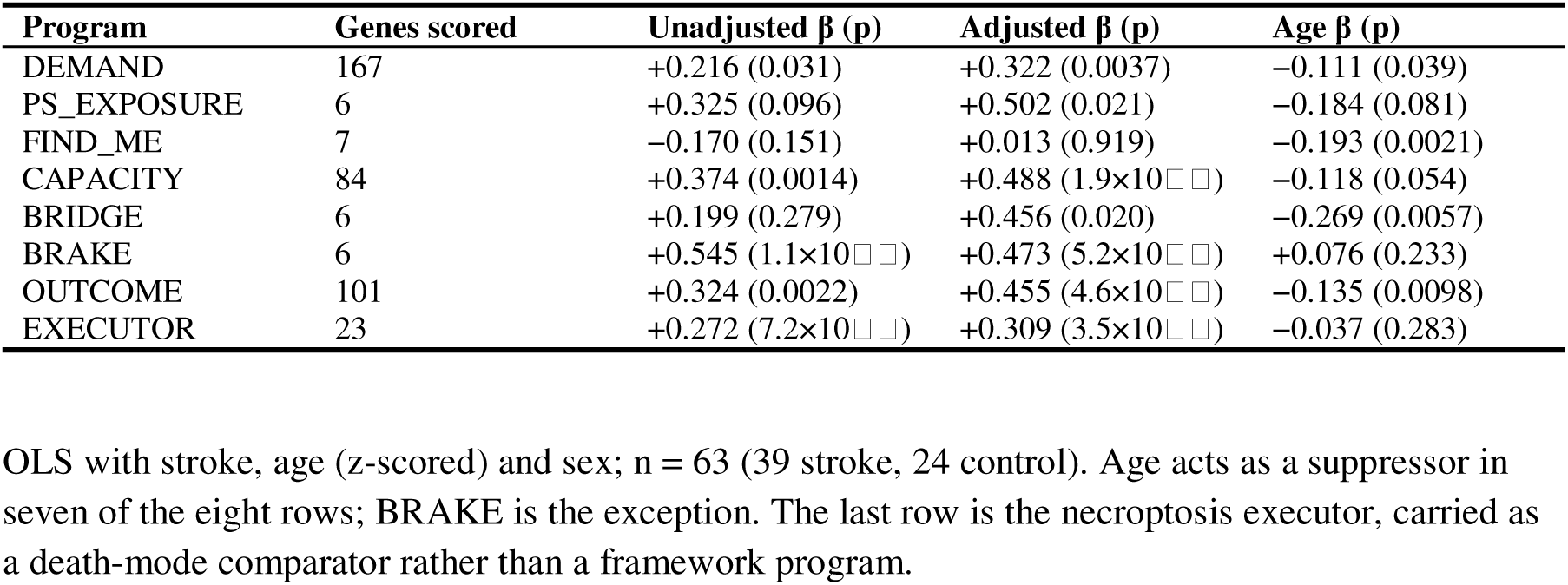
Age and sex adjustment of program scores in GSE16561.

Two controls show that the attenuation is specific to the proxies rather than a generic penalty for adding covariates. Gene sets unrelated to all five proxies moved by a median of 0.036 SD in GSE16561 and 0.025 SD in GSE58294, while CAPACITY moved by 0.309 and 0.164 SD — beyond every one of the 200 matched draws in both cohorts. The gradient alone removes 28.3% of the coefficient in GSE16561 and 0.30% in GSE58294, so it cannot by itself account for the proxy-driven attenuation. But nor can it be treated as independent of it: at ρ = 0.814 in GSE16561 the proxy and gradient blocks share variance, and the joint model leaves a residual coefficient that is still nominally significant (+0.109, p = 0.043). The defensible statement is that the attenuation is specific to the lineage proxies and that its magnitude is not identified separately from the array gradient in that cohort.

Oaxaca–Blinder decomposition of the raw gap points in the same direction without fixing a magnitude (Table 6): composition contributes 22.8 to 23.5% in GSE58294 and 52.1 to 104.5% in GSE16561, a spread that reflects the choice of reference base rather than a precise estimate. No program is carried by one gene or one module: dropping any single CAPACITY gene moved the estimate by less than 0.02, and all eight CAPACITY modules were positive in both cohorts (Table 7E).

**Table 6.** Oaxaca–Blinder decomposition of the CAPACITY gap in whole blood.

| **Cohort** | **Covariates** | **Basis** | **Composition %<br>(95% CI)** | **Within-cell %<br>(95% CI)** | **Raw gap β** |
|---|---|---|---|---|---|
| GSE58294 | 5 lineage proxies | A (reference-priced) | +22.96 (−2.8 to 46.8) | +77.04 (55.5 to 103.0) | 0.613 |
| GSE58294 | 5 lineage proxies | B (target-priced) | +23.53 (−29.5 to 62.5) | +76.47 (42.7 to 128.8) | 0.613 |
| GSE58294 | 5 lineage proxies | Symmetric | +23.24 (−7.7 to 50.5) | +76.76 (56.9 to 106.0) | 0.613 |
| GSE58294 | 5 lineage proxies | Pooled-β | +22.81 (2.2 to 47.5) | +77.19 (58.8 to 95.6) | 0.613 |
| GSE16561 | Age, sex, 5 proxies | A | +52.10 (−47.6 to 183.6) | +47.00 (−92.4 to 118.3) | 0.374 |
| GSE16561 | Age, sex, 5 proxies | B | +104.51 (37.1 to 161.8) | −3.10 (−52.6 to 44.5) | 0.374 |
| GSE16561 | Age, sex, 5 proxies | Symmetric | +78.30 (10.0 to 166.1) | +21.95 (−52.6 to 66.4) | 0.374 |
| GSE16561 | Age, sex, 5 proxies | Pooled-β | +97.88 (31.8 to 156.5) | +3.78 (−38.0 to 41.5) | 0.374 |

The decomposition is of the raw case–control gap, a different estimand from the coefficient attenuation in Table 3; the pooled-basis value of 97.9% in GSE16561 should not be expected to equal the 97.1% in that table. Age and the lineage proxies are not separately identified in GSE16561, so the age and composition terms should be read from the sequential decomposition in the text.

**Table 7.**
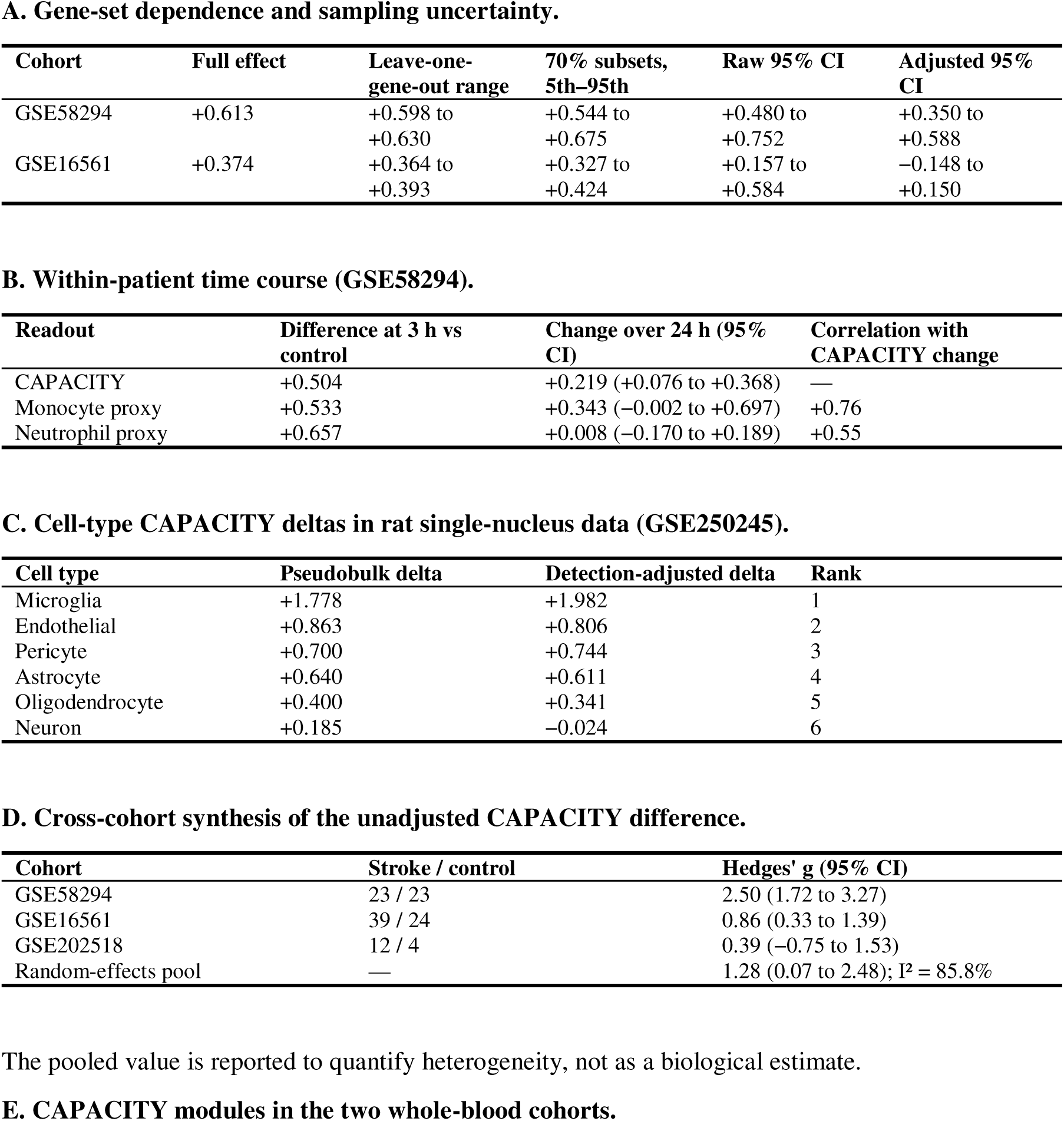
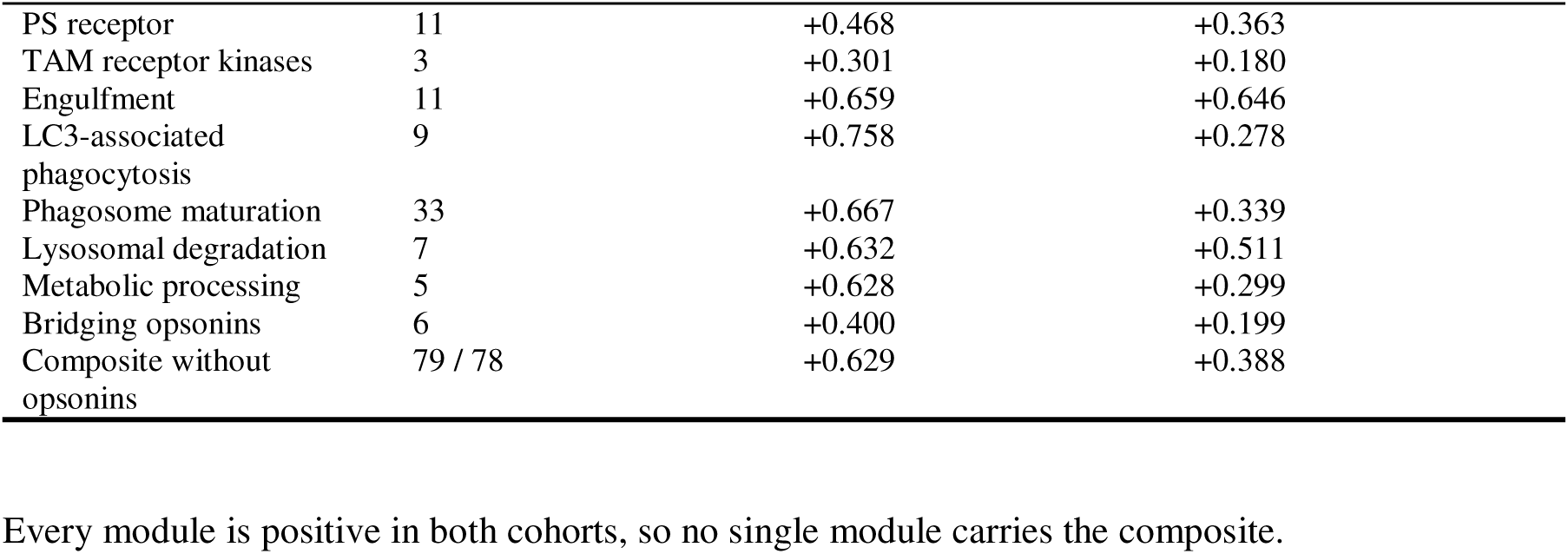
Robustness of the blood signal and the cell-type ranking.

| A. Gene-set dependence and sampling uncertainty. |  |  |  |  |  |
|---|---|---|---|---|---|
| Cohort | Full effect | Leave-one-gene-out range | 70% subsets, 5th–95th | Raw 95% CI | Adjusted 95% CI |
| GSE58294 | +0.613 | +0.598 to +0.630 | +0.544 to +0.675 | +0.480 to +0.752 | +0.350 to +0.588 |
| GSE16561 | +0.374 | +0.364 to +0.393 | +0.327 to +0.424 | +0.157 to +0.584 | −0.148 to +0.150 |
| B. Within-patient time course (GSE58294). |  |  |  |  |  |
| Readout | Difference at 3 h vs control | Change over 24 h (95% CI) | Correlation with CAPACITY change |  |  |
| CAPACITY | +0.504 | +0.219 (+0.076 to +0.368) | — |  |  |
| Monocyte proxy | +0.533 | +0.343 (−0.002 to +0.697) | +0.76 |  |  |
| Neutrophil proxy | +0.657 | +0.008 (−0.170 to +0.189) | +0.55 |  |  |
| C. Cell-type CAPACITY deltas in rat single-nucleus data (GSE250245). |  |  |  |  |  |
| Cell type | Pseudobulk delta | Detection-adjusted delta | Rank |  |  |
| Microglia | +1.778 | +1.982 | 1 |  |  |
| Endothelial | +0.863 | +0.806 | 2 |  |  |
| Pericyte | +0.700 | +0.744 | 3 |  |  |
| Astrocyte | +0.640 | +0.611 | 4 |  |  |
| Oligodendrocyte | +0.400 | +0.341 | 5 |  |  |
| Neuron | +0.185 | −0.024 | 6 |  |  |
| D. Cross-cohort synthesis of the unadjusted CAPACITY difference. |  |  |  |  |  |
| Cohort | Stroke / control | Hedges' g (95% CI) |  |  |  |
| GSE58294 | 23 / 23 | 2.50 (1.72 to 3.27) |  |  |  |
| GSE16561 | 39 / 24 | 0.86 (0.33 to 1.39) |  |  |  |
| GSE202518 | 12 / 4 | 0.39 (−0.75 to 1.53) |  |  |  |
| Random-effects pool | — | 1.28 (0.07 to 2.48); I² = 85.8% |  |  |  |
| The pooled value is reported to quantify heterogeneity, not as a biological estimate. |  |  |  |  |  |
| E. CAPACITY modules in the two whole-blood cohorts. |  |  |  |  |  |
| PS receptor | 11 | +0.468 | +0.363 |  |  |
| TAM receptor kinases | 3 | +0.301 | +0.180 |  |  |
| Engulfment | 11 | +0.659 | +0.646 |  |  |
| LC3-associated phagocytosis | 9 | +0.758 | +0.278 |  |  |
| Phagosome maturation | 33 | +0.667 | +0.339 |  |  |
| Lysosomal degradation | 7 | +0.632 | +0.511 |  |  |
| Metabolic processing | 5 | +0.628 | +0.299 |  |  |
| Bridging opsonins | 6 | +0.400 | +0.199 |  |  |
| Composite without opsonins | 79 / 78 | +0.629 | +0.388 |  |  |

Every module is positive in both cohorts, so no single module carries the composite.

The PBMC control points the same way. In GSE22255, depleted of granulocytes and matched for age and sex, CAPACITY did not differ between stroke and control (−0.054; 90% CI −0.218 to +0.110; p = 0.582, q = 0.914). The interval does not establish equivalence at a 0.3-SD margin, and at n = 20 the power to detect the observed difference is effectively zero, so this is a bound rather than proof of no effect. Serial sampling in GSE58294 does not break the tie: CAPACITY was +0.504 above controls at 3 h and rose by +0.219 over 24 h, but the monocyte proxy rose by +0.343 over the same interval and the two changes correlated at r = 0.76 — a program that moves with the monocytes is not evidence of a program that moves within them. The defensible statement is therefore narrower than “peripheral capacity rises after stroke”: a reproducible change in the circulating myeloid efferocytosis program accompanies stroke, but whether it reflects per-cell regulation or a shift in leukocyte composition differs between cohorts and cannot be settled from bulk blood.

### Human infarct core: disorganisation, not loss of transcripts

If stroke suppressed clearance in the brain, the infarct core should show lower CAPACITY than the contralateral cortex. In six paired post-mortem donors (GSE162955), it did not do so coordinately. The CAPACITY difference was −0.181 (FDR 0.81), and none of 29 scored layers was reproducible across donors. The burden and output layers moved upward instead (output FDR = 0.055), while an oxidative-phosphorylation anchor fell by 0.279. The core is necrotic tissue, so the comparison cannot separate loss of viable phagocytes, RNA degradation and a genuine per-cell failure. It does exclude the simple claim that CAPACITY falls wherever infarcted tissue is sampled: transcripts remain measurable in the core, and 59 of 81 CAPACITY genes are in fact down-regulated (p = 4.8×10□□). The failure is interpretive — a necrotic source cannot be read as per-cell clearance — not a failure of detection. A second human neocortex series, GSE56267, could not be used: its case–control estimate moves from +0.135 (p = 0.52) to +0.453 (p = 0.030) depending on where three unassigned libraries are placed (Table 1).

The human brain data therefore say something more specific than “central failure”: a bulk transcriptome from infarcted tissue is not a reliable instrument for per-cell clearance, because the tissue itself is disorganised. The central side of the dissociation is better supported by the rodent cell-type data, where viable cells can be sorted.

### Rodent brain shows inducible capacity whose carrier changes with time

In rat single-nucleus RNA-seq (GSE250245), microglial CAPACITY rose by +1.853 in the pooled ipsilateral-versus-contralateral comparison (95% bootstrap CI 1.757–1.943), the largest response among six annotated cell types (Table 8; Fig. 4A). Capture depth is the obvious alternative — each nucleus contains on average 7.8 of the 85 CAPACITY genes — but ipsilateral microglia were captured more shallowly than contralateral (1,170 versus 1,420 genes per nucleus), which works against the signal. Refitting each delta with linear and quadratic terms for detected genes preserved the rank order (+1.982), and two further aggregations (pseudobulk and nucleus-summed) agreed on direction. The comparison rests on one animal per severity and hemisphere, so we treat it as a direction rather than an animal-level estimate.

**Figure 3.**
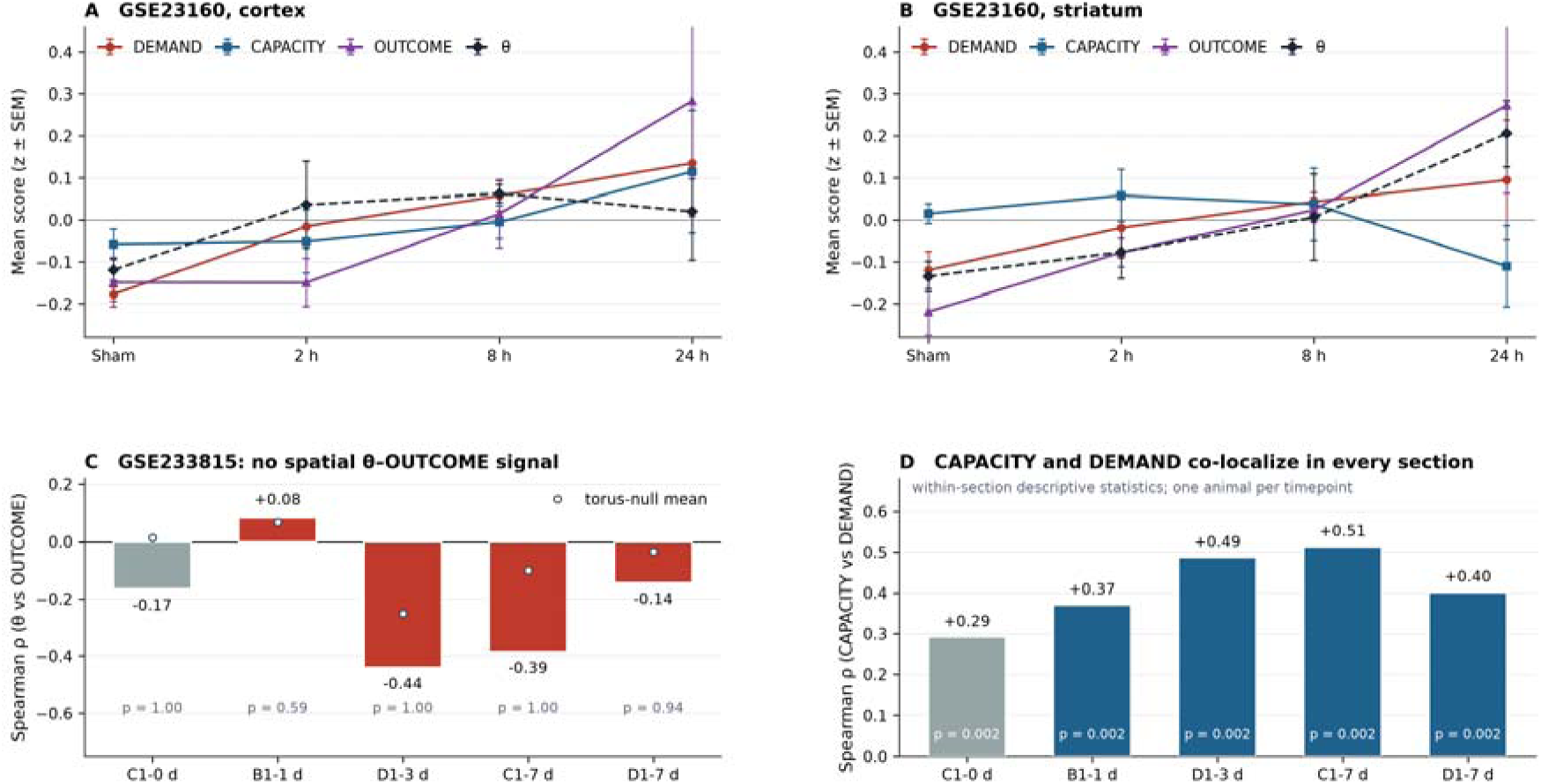
θ in time and space. (A-B) GSE23160: θ, DEMAND, CAPACITY and OUTCOME across 2, 8 and 24 h of reperfusion in cortex and striatum. (C) GSE233815: θ–OUTCOME Spearman ρ across five Visium sections with torus-shift nulls; no section reached nominal significance (p = 0.592 to 1.000). (D) CAPACITY–DEMAND co-localisation (ρ = +0.291 to +0.513), showing why their difference carries little independent spatial structure.

**Figure 4.**
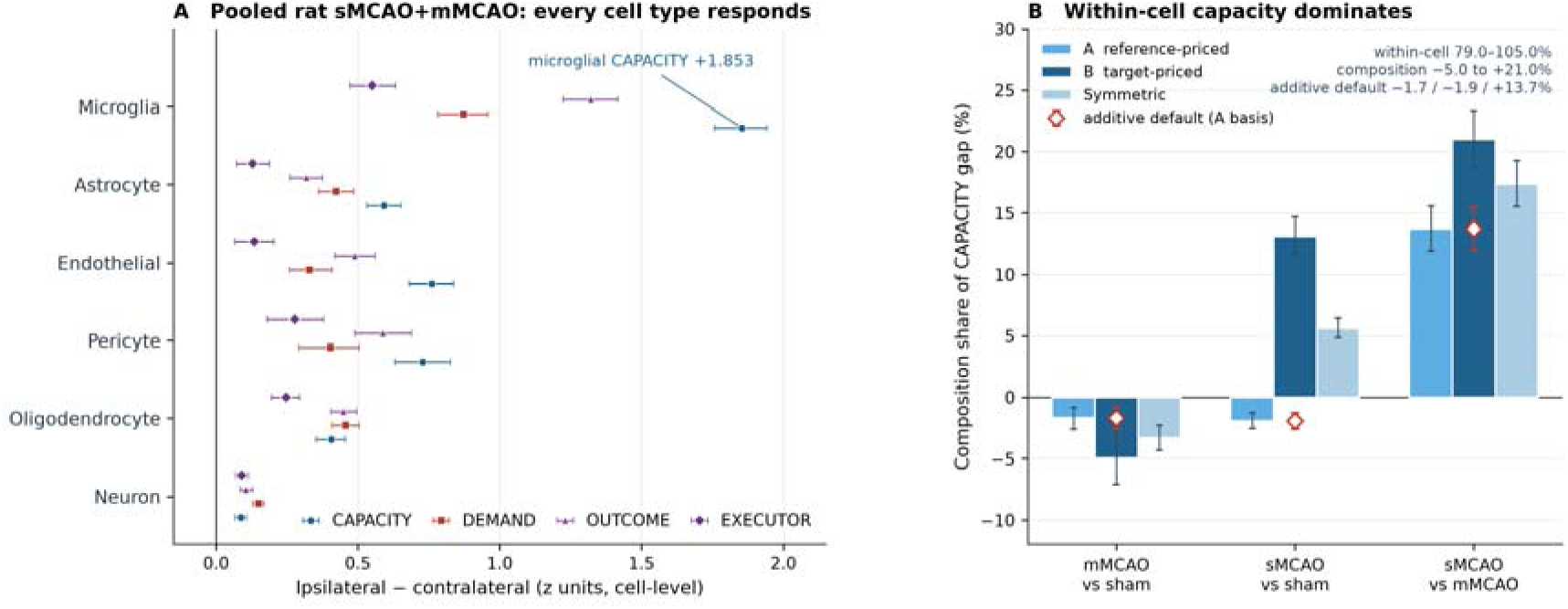
Cell type and composition. (A) Pooled rat sMCAO and mMCAO libraries: CAPACITY, DEMAND, OUTCOME and executor deltas by cell type; microglia show the largest CAPACITY response (+1.853). (B) Oaxaca–Blinder decomposition of the CAPACITY gap under three reference bases: composition −5.0% to +21.0% and within-cell capacity 79.0% to 105.0%.

**Table 8.** Cell-type-specific program deltas in pooled rat sMCAO and mMCAO (GSE250245).

| **Cell type** | **Nuclei (ipsi/contra)** | **CAPACITY Δ (95% CI)** | **Detection-adjusted Δ** | **OUTCOME Δ (95% CI)** | **Split-half sign consistency** |
|---|---|---|---|---|---|
| Microglia | 1608/1687 | +1.853 (1.757 to 1.943) | +1.982 | +1.321 (1.224 to 1.416) | 1.00 |
| Endothelial | 1813/2279 | +0.760 (0.681 to 0.837) | +0.806 | +0.490 (0.418 to 0.562) | 1.00 |
| Pericyte | 1908/576 | +0.729 (0.629 to 0.826) | +0.744 | +0.588 (0.489 to 0.688) | 1.00 |
| Astrocyte | 2427/2539 | +0.592 (0.531 to 0.651) | +0.611 | +0.319 (0.260 to 0.375) | 1.00 |
| Oligodendrocyte | 3846/4091 | +0.406 (0.353 to 0.455) | +0.341 | +0.449 (0.404 to 0.494) | 1.00 |
| Neuron | 10140/14133 | +0.087 (0.067 to 0.107) | −0.024 | +0.106 (0.086 to 0.129) | 1.00 |

Deltas are ipsilateral minus contralateral within the pooled libraries. Each severity-by-hemisphere stratum contains one animal, so these are cell-level descriptors rather than animal-level precision. Intervals are cell-level stratified bootstrap intervals from six libraries. Detection-adjusted Δ refits each cell type's delta with linear and quadratic terms for detected genes per nucleus; the rank order is unchanged.

The decomposition places the rat effect where the blood data did not. Across three reference bases, composition contributed −5.0% to +21.0% of the CAPACITY gap and within-cell capacity 79.0% to 105.0% (Table 9; Fig. 4B). In brain, unlike blood, the signal is carried mainly by a change within cells rather than in which cells are present.

**Table 9.** Oaxaca–Blinder decomposition of the CAPACITY gap in rat cell types (GSE250245).

| Contrast | Basis | Composition % (95% CI) | Within-cell % | ΔS |
|---|---|---|---|---|
| mMCAO − sham | Reference-priced | −1.70 (−2.60 to −0.83) | 101.70 | 0.479 |
| mMCAO − sham | Target-priced | −4.96 (−7.08 to −2.87) | 104.96 | 0.479 |
| mMCAO − sham | Symmetric | −3.33 (−4.33 to −2.29) | 103.33 | 0.479 |

Composition and within-cell terms sum to 100 at each basis by construction. Bootstrap intervals are stratified by library, B = 1,000, and are cell-level rather than animal-level.

Sorted-cell series resolve the time dependence that whole-tissue data blur. At 48 h after tMCAO (GSE105132), infiltrating macrophages and neutrophils scored far above microglia on LAP (+0.858 and +1.140; Cliff’s δ = 1.000 for both). At day 3 (GSE141972), the ordering reversed: resident microglia scored higher than infiltrating macrophages on CAPACITY (macrophage minus microglia −0.274; δ = −0.667). By day 7 (GSE86941), the signal was carried by macrophages: +0.340 against sham (δ = 0.625; q = 0.0015) and +0.193 against resident microglia (q = 0.0276), while microglia did not clear correction (+0.147; q = 0.0583). The carrier is not fixed. These are three different models and sorting schemes, so the carrier is a direction within each series rather than a statement that the same cell moved; what the series establish is that a single time point cannot identify the carrier.

The earliest time point sharpens the ordering. GSE137595 measures mouse cortex 2 h after 90 min MCAO with reperfusion, in a 2 × 2 design (ischaemic versus sham animal, crossed with hemisphere). DEMAND rose by +0.239 and exceeded its expression-matched null, whereas CAPACITY did not move (+0.043; p = 0.66). The demand–capacity gap opened because demand rose, not because machinery was withdrawn.

A bioenergetic program moves earlier than CAPACITY in the subacute window. In GSE86941 at day 3, oxidative phosphorylation rose by +1.512 in macrophages and +0.801 in microglia (q ≤ 0.0006), and glycolysis by +0.413 and +0.213 (q ≤ 0.0009), while CAPACITY did not clear correction in either population (+0.173, q = 0.2944; +0.050, q = 0.2734). Across the six contrasts of that series the bioenergetic term leads the machinery term in five. The energy that engulfment requires [30–33] arrives before the machinery itself (Table 10). This cannot exclude that the bioenergetic transcripts track general activation rather than clearance, but the three canonical biogenesis drivers move the other way in all six contrasts (PPARGC1A −1.88 to −0.11, NRF1 −1.97 to −0.32, TFAM −1.97 to +0.23), so it is not mitochondrial biogenesis.

**Table 10.** A bioenergetic layer leads the clearance machinery in the acute phase (GSE86941).

| Contrast | OXPHOS | GLYCOLYSIS | HYPOXIA | CAPACITY | DEGRADATION | Stage passing FDR |
|---|---|---|---|---|---|---|
| 3 d macrophage vs sham | +1.512 (0.0006)* | +0.413 (0.0009)* | +0.200 (0.0172)* | +0.173 (0.2944) | +0.907 (0.2550) | bioenergetic only |
| 3 d microglia vs sham | +0.801 (0.0003)* | +0.213 (0.0003)* | +0.094 (0.0036)* | +0.050 (0.2734) | +0.836 (0.1368) | bioenergetic only |
| 3 d macrophage vs microglia | +0.710 (0.0006)* | +0.200 (0.1650) | +0.106 (0.5685) | +0.123 (0.8696) | +0.071 (0.9817) | bioenergetic only |
| 7 d macrophage vs sham | +1.320 (0.0006)* | +0.097 (0.0015)* | −0.100 (0.1308) | +0.340 (0.0015)* | +1.180 (0.0314)* | both |
| 7 d microglia vs sham | +1.198 (0.0006)* | −0.031 (0.1359) | −0.196 (0.9887) | +0.147 (0.0583) | +1.207 (0.0184)* | bioenergetic only |
| 7 d macrophage vs microglia | +0.122 (0.0042)* | +0.128 (0.0042)* | +0.096 (0.0078)* | +0.193 (0.0276)* | −0.027 (0.9153) | both |

Values are observed layer deltas in z units with Benjamini–Hochberg q in parentheses, from size-matched permutation nulls (B = 10,000). The family is the six contrasts of this series for the layer concerned; it is not the same family as in Table 14 or Table 4B, and q values from these tables are not compared. Asterisks mark q < 0.05.

### Sorted human parenchyma settles composition, not capacity

A series that sorts CD45-positive cells from fresh human parenchyma (GSE331114) answers the composition question that homogenate cannot. Two properties of the series bound every statement below. Sequencing depth varies 5.5-fold across the eight libraries, and the shallowest library is a stroke donor. The sort is also incomplete: PTPRC is detected in 0.68 to 0.93 of cells per library, and 9.4% of cells fall into non-immune classes — a fraction that reaches 24.6% and 26.3% in two control donors but stays below 10% in all four stroke donors. Contamination and group are therefore partly the same variable, so every inference is restricted to a named compartment.

Three findings follow.

First, composition moved in the direction the rodent data predict. Non-microglial myeloid cells constituted 37.1% of cells recovered from a CD45-positive sort in stroke donors versus 19.9% in controls (+0.173; exact permutation p = 0.057 over all 70 ways of splitting eight donors four and four), while resident microglia fell from 27.6% to 16.7% and granulocytes from 30.4% to 22.1% (Table 4A). With four donors per group the smallest attainable two-sided p is 2/70 = 0.0286, so no composition difference survives correction across the seven compartments; the result is a direction with a plausible mechanism, not a tested effect.

Second, one part of the machinery fell and stayed robust. The bridging-opsonin set in infiltrating myeloid cells moved −0.903, 1.58 times the smallest difference this series can resolve (p = 0.003 against expression-matched random gene sets; q = 0.049 across the nineteen framework programs scored in that compartment), kept its sign in all eight leave-one-donor-out refits (−1.079 to −0.787), and stayed negative under all four covariate adjustments (−0.941 for depth, −0.806 for ambient myelin, −0.510 for non-immune contamination, −0.232 for the infiltrating fraction itself). Composite CAPACITY in the same cells reached 1.07 times that difference (q = 0.23) and reversed sign once ambient RNA was entered, so we report no capacity result from this dataset. Six opsonin genes move; the 85-gene composite does not. A small set clears a given threshold more easily than a large one, so what is established is that the opsonin set is lower, not that clearance is. Two further rows in Table 11 need a note. Oxidative phosphorylation in infiltrating myeloid cells rose +0.197 (1.22 times its limit, q = 0.013), which fits a metabolically active infiltrate. In T cells it rose +0.546 (2.05 times, q < 0.001), the largest standardised effect in the human data; T cells do not engulf, so this is either genuine metabolic activation or a dissociation and ambient-RNA artefact in a series whose depth varies 5.5-fold. Neither row is used to support a capacity claim.

**Table 11.**
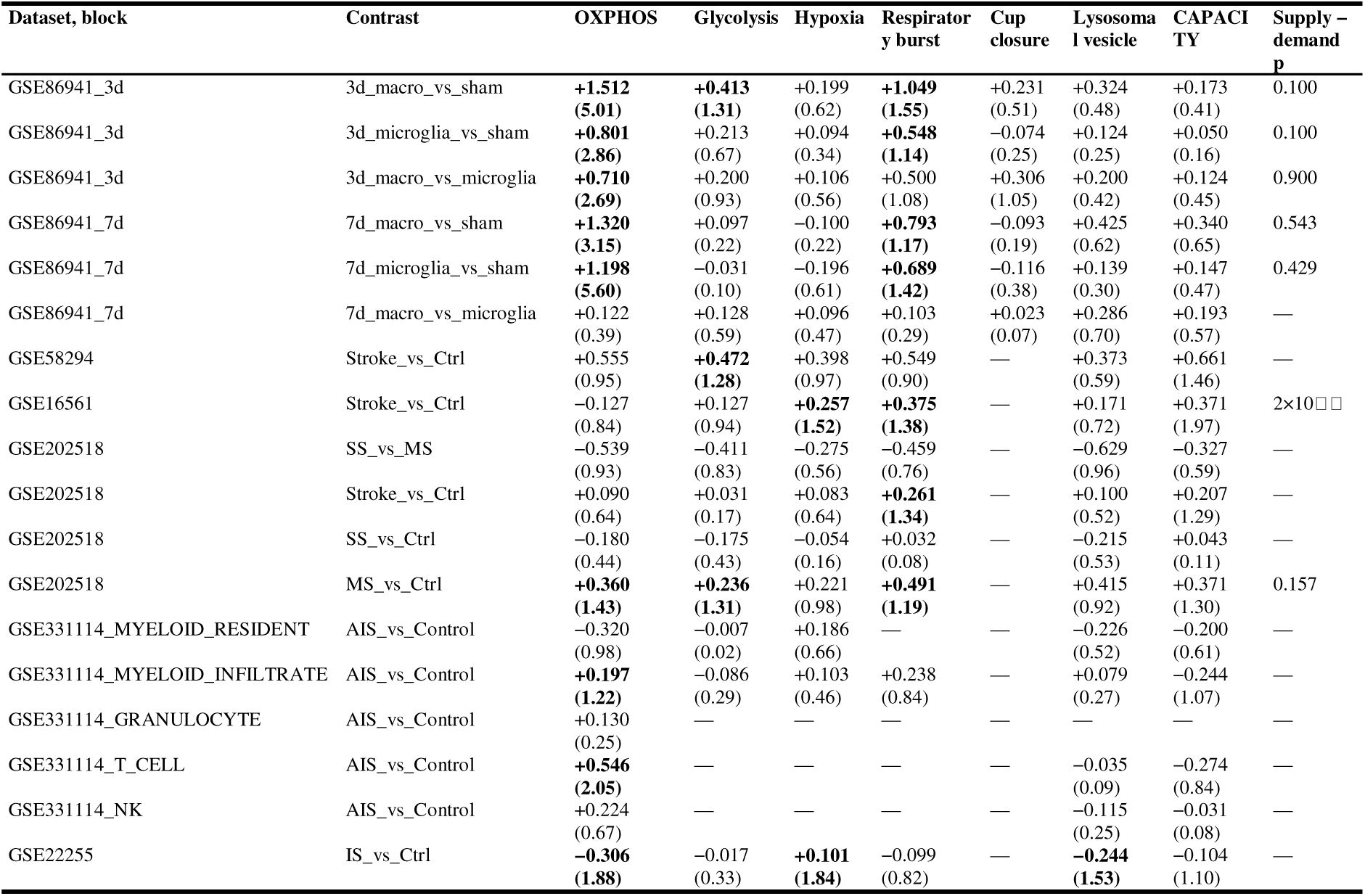
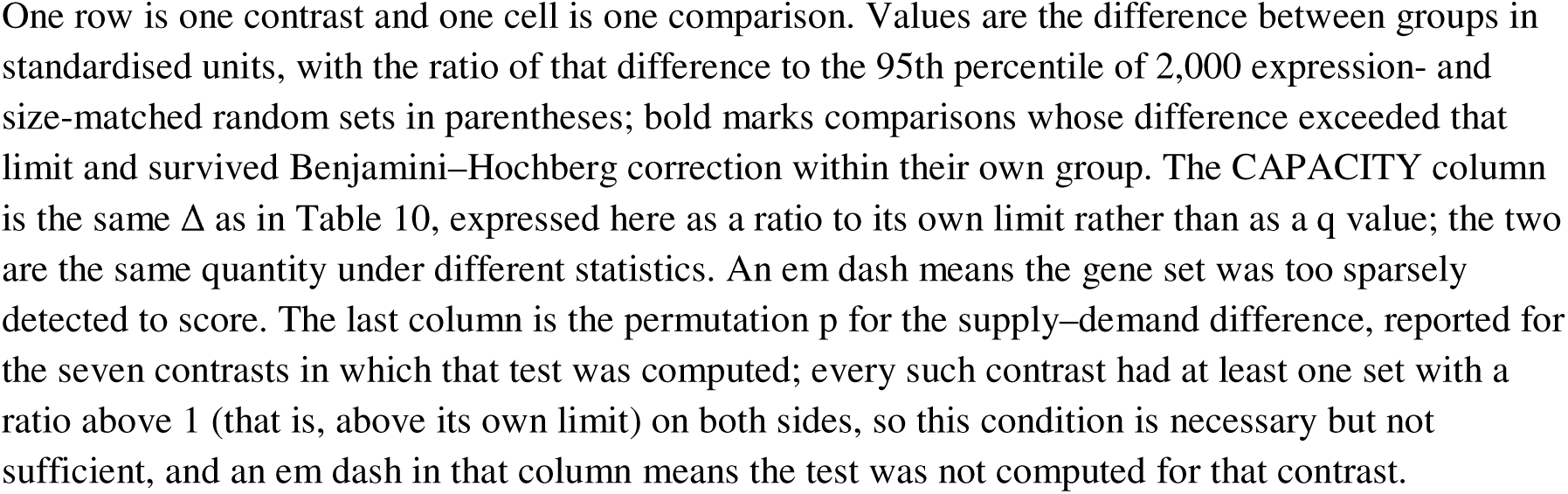
Supply and demand for the energy the program needs (eighteen contrasts across six datasets).

| Dataset, block | Contrast | OXPHOS | Glycolysis | Hypoxia | Respiratory burst | Cup closure | Lysosomal vesicle | CAPACITY | Supply – demand p |
|---|---|---|---|---|---|---|---|---|---|
| GSE86941_3d | 3d_macro_vs_sham | **+1.512 (5.01)** | **+0.413 (1.31)** | +0.199 (0.62) | **+1.049 (1.55)** | +0.231 (0.51) | +0.324 (0.48) | +0.173 (0.41) | 0.100 |
| GSE86941_3d | 3d_microglia_vs_sham | **+0.801 (2.86)** | +0.213 (0.67) | +0.094 (0.34) | **+0.548 (1.14)** | −0.074 (0.25) | +0.124 (0.25) | +0.050 (0.16) | 0.100 |
| GSE86941_3d | 3d_macro_vs_microglia | **+0.710 (2.69)** | +0.200 (0.93) | +0.106 (0.56) | +0.500 (1.08) | +0.306 (1.05) | +0.200 (0.42) | +0.124 (0.45) | 0.900 |
| GSE86941_7d | 7d_macro_vs_sham | **+1.320 (3.15)** | +0.097 (0.22) | −0.100 (0.22) | **+0.793 (1.17)** | −0.093 (0.19) | +0.425 (0.62) | +0.340 (0.65) | 0.543 |
| GSE86941_7d | 7d_microglia_vs_sham | **+1.198 (5.60)** | −0.031 (0.10) | −0.196 (0.61) | **+0.689 (1.42)** | −0.116 (0.38) | +0.139 (0.30) | +0.147 (0.47) | 0.429 |
| GSE86941_7d | 7d_macro_vs_microglia | +0.122 (0.39) | +0.128 (0.59) | +0.096 (0.47) | +0.103 (0.29) | +0.023 (0.07) | +0.286 (0.70) | +0.193 (0.57) | — |
| GSE58294 | Stroke_vs_Ctrl | +0.555 (0.95) | **+0.472 (1.28)** | +0.398 (0.97) | +0.549 (0.90) | — | +0.373 (0.59) | +0.661 (1.46) | — |
| GSE16561 | Stroke_vs_Ctrl | −0.127 (0.84) | +0.127 (0.94) | **+0.257 (1.52)** | **+0.375 (1.38)** | — | +0.171 (0.72) | +0.371 (1.97) | 2×10[illegible] |
| GSE202518 | SS_vs_MS | −0.539 (0.93) | −0.411 (0.83) | −0.275 (0.56) | −0.459 (0.76) | — | −0.629 (0.96) | −0.327 (0.59) | — |
| GSE202518 | Stroke_vs_Ctrl | +0.090 (0.64) | +0.031 (0.17) | +0.083 (0.64) | **+0.261 (1.34)** | — | +0.100 (0.52) | +0.207 (1.29) | — |
| GSE202518 | SS_vs_Ctrl | −0.180 (0.44) | −0.175 (0.43) | −0.054 (0.16) | +0.032 (0.08) | — | −0.215 (0.53) | +0.043 (0.11) | — |
| GSE202518 | MS_vs_Ctrl | **+0.360 (1.43)** | **+0.236 (1.31)** | +0.221 (0.98) | **+0.491 (1.19)** | — | +0.415 (0.92) | +0.371 (1.30) | 0.157 |
| GSE331114_MYELOID_RESIDENT | AIS_vs_Control | −0.320 (0.98) | −0.007 (0.02) | +0.186 (0.66) | — | — | −0.226 (0.52) | −0.200 (0.61) | — |
| GSE331114_MYELOID_INFILTRATE | AIS_vs_Control | **+0.197 (1.22)** | −0.086 (0.29) | +0.103 (0.46) | +0.238 (0.84) | — | +0.079 (0.27) | −0.244 (1.07) | — |
| GSE331114_GRANULOCYTE | AIS_vs_Control | +0.130 (0.25) | — | — | — | — | — | — | — |
| GSE331114_T_CELL | AIS_vs_Control | **+0.546 (2.05)** | — | — | — | — | −0.035 (0.09) | −0.274 (0.84) | — |
| GSE331114_NK | AIS_vs_Control | +0.224 (0.67) | — | — | — | — | −0.115 (0.25) | −0.031 (0.08) | — |
| GSE22255 | IS_vs_Ctrl | **−0.306 (1.88)** | −0.017 (0.33) | **+0.101 (1.84)** | −0.099 (0.82) | — | **−0.244 (1.53)** | −0.104 (1.10) | — |

One row is one contrast and one cell is one comparison. Values are the difference between groups in standardised units, with the ratio of that difference to the 95th percentile of 2,000 expression- and size-matched random sets in parentheses; bold marks comparisons whose difference exceeded that limit and survived Benjamini–Hochberg correction within their own group. The CAPACITY column is the same Δ as in Table 10, expressed here as a ratio to its own limit rather than as a q value; the two are the same quantity under different statistics. An em dash means the gene set was too sparsely detected to score. The last column is the permutation p for the supply–demand difference, reported for the seven contrasts in which that test was computed; every such contrast had at least one set with a ratio above 1 (that is, above its own limit) on both sides, so this condition is necessary but not sufficient, and an em dash in that column means the test was not computed for that contrast.

Third, the capacity question cannot be answered at this depth. The smallest resolvable difference for each program in each compartment runs from 0.15 to 0.90 z units. Against it, none of the 21 layers scored in resident microglia is distinguishable from an arbitrary equally expressed set, and neither is any of the 14 scored in NK cells. CAPACITY in microglia moved by −0.200 against a threshold of 0.331, 0.60 of the smallest difference that series can resolve, so the correct description of that row is not that microglial capacity was unchanged but that this series cannot tell. The same holds for the brake node the framework was built around: the six-gene brake is below its threshold in every compartment where it could be scored, so neither a claim that the brake tightens nor one that it is released is available. Single transcripts agree, once each donor is counted once and every cell is downsampled to a common depth: CD300A is static in both myeloid compartments that carry it (Table 4D), and its apparent raw release was a weighting artefact driven by one deep control donor.

### Energy arrives before the machinery

The work of clearance — polymerising actin into a cup, pinching the membrane, acidifying the phagosome — is part of the claim rather than an addendum to it. Measured on one scale across eighteen contrasts in six datasets, supply behaved as the rat series predicted: fourteen comparisons exceeded both the expression-matched threshold and correction within their own group, spread over eleven of the eighteen contrasts (Table 11). The largest were in the cells that had just entered the lesion: at day 3, oxidative phosphorylation in infiltrating macrophages moved +1.512, five times its own threshold, with glycolysis at +0.413, and at day 7 the same layer stood at +1.320 in macrophages and +1.198 in microglia. In the same day-3 macrophages, CAPACITY moved +0.173 — 0.41 of its own threshold — and never entered correction. A large shift in supply alongside a shift in uptake that the same design cannot place above its threshold is indirect evidence of ordering, not a demonstrated sequence. The two programs also have different thresholds (0.301 for oxidative phosphorylation, 0.42 for CAPACITY), and a wider threshold makes the machinery harder to pass. We did not test a program × time interaction; what the data show is that each layer crosses its own detection limit at a different time.

Demand was detected above both gates in eight comparisons across eight contrasts, and in the rat it appeared in the same populations where supply moved. The respiratory-burst set rose +1.049 in day-3 infiltrating macrophages, +0.793 at day 7 in macrophages and +0.689 in microglia. Cup closure exceeded its threshold in only one contrast and did not survive correction there (+0.306 in day-3 macrophages against microglia, q = 0.124), and lysosomal vesicle biogenesis never rose above its threshold anywhere. The proton-pump and oxidase part of the demand side rises with the supply side, while the actin and vesicle-biogenesis parts cannot be resolved — consistent with a cell already competent to engulf that is increasing its killing and acidification capacity.

In whole blood the three cohorts disagreed, and the disagreement is the result. GSE58294 moved on glycolysis (+0.472, 1.28 times its threshold); GSE16561 on hypoxia (+0.257) and the respiratory burst (+0.375); GSE202518 on the respiratory burst (+0.261). Direction does not carry across cohorts; what carries is that in each of them at least one supply set and at least one demand set exceed their thresholds, and the set that does so is never the same one twice.

The supply–demand difference could be tested in seven of the eighteen contrasts (Table 11), and six gave nothing (p between 0.100 and 0.900). The seventh, GSE16561, gave −0.187 at p = 0.0002 — the demand side rose further than supply — but that does not survive composition adjustment: the constituent sets of both sides move towards zero in that cohort (supply to +0.039, −0.020 and −0.073, and the respiratory burst to +0.116 with an interval spanning zero), so the gap tracks the leukocyte shift rather than a per-cell deficit. Across the contrasts the two scores ranked samples almost identically, ρ between 0.590 and 0.952 (0.429 to 0.933 after removing each sample’s global expression axis). Where composition adjustment is identified it changes the answer: in GSE58294 the energy shift survives it (glycolysis +0.400, hypoxia +0.350, oxidative phosphorylation +0.274, respiratory burst +0.360; adjusted permutation p ≤ 0.007), while in GSE16561 the same adjustment removes it.

None of this measures flux. A transcript score for oxidative phosphorylation reports the abundance of messages for the machinery, not the rate at which it turns over ATP. What the exercise establishes is a boundary: in sorted brain cells the supply and demand sides of efferocytosis rise together in the cells that have just arrived; in whole blood the same measurement separates the program from its cells only in the one cohort where the proxies leave enough of the group label to estimate.

### The demand–capacity difference does not add information

We defined θ = DEMAND − CAPACITY as a difference, not a ratio, because centred scores have no stable zero (Fig. 3A, B). In GSE23160 (mouse MCAO with 2/8/24 h reperfusion; n = 4 per group), θ correlated with OUTCOME in the striatum (ρ = 0.768; permutation p = 2.7×10□³), but DEMAND correlated at least as strongly (ρ = 0.806; p = 1.33×10□³). Nested models made the same point: in cortex, OUTCOME ∼ DEMAND explained R² = 0.661 and adding CAPACITY raised this to 0.724 (ΔR² = 0.063; p = 0.108); in striatum, 0.762 to 0.819 (ΔR² = 0.057; p = 0.064). Information criteria pointed the other way (ΔAIC 1.31 and 2.40), and neither supports treating θ as an improvement on DEMAND (Table 12). Across five Visium sections the θ–OUTCOME correlation ranged from −0.441 to +0.083 with no section reaching significance (Table 13; Fig. 3C), because CAPACITY and DEMAND are tightly co-localised (ρ = +0.291 to +0.513; Fig. 3D), leaving a near-random difference. θ remains a descriptive shorthand, not a predictor: in these data DEMAND carries the temporal association and CAPACITY adds little at n = 16. We scored the necroptosis executor in parallel as a death-mode comparator, not as a component of the framework. It rose by 24 h in both regions (+0.198 in cortex, permutation p = 0.114, BH q = 0.190; +0.308 in striatum, p = 0.029 at the 2/70 permutation floor, q = 0.143) and reached family-wise significance in neither. The expectation pre-specified in the locked analysis script before the results were inspected — pre-specification of one analysis, not registration of the study — was not supported in either region (cortex D_early = +0.469, 95% CI −0.440 to +1.575, P(D ≤ 0) = 0.140; striatum +0.803, 95% CI −0.472 to +2.509, P = 0.145); we report this null in full. The executor shares no gene with CAPACITY in any species, so the two are not algebraically redundant; the nonsignificant comparator does not by itself establish specificity, but it removes a simple shared-gene explanation for the CAPACITY signal.

**Table 12.** Incremental validity of θ and CAPACITY beyond DEMAND (GSE23160, n = 16 per region).

| **Region** | **Model** | **R²** | **ΔR² vs DEMAND** | **F** | **p** | **β CAPACITY** | **AIC** |
|---|---|---|---|---|---|---|---|
| Cortex | DEMAND | 0.661 | — | — | — | — | −7.55 |
| Cortex | DEMAND + CAPACITY | 0.724 | 0.063 | 2.98 | 0.108 | +0.489 | −8.86 |
| Cortex | θ only | 0.031 | — | — | — | — | +9.25 |
| Striatum | DEMAND | 0.762 | — | — | — | — | −13.01 |
| Striatum | DEMAND + CAPACITY | 0.819 | 0.057 | 4.11 | 0.064 | −0.476 | −15.41 |
| Striatum | θ only | 0.591 | — | — | — | — | −4.36 |

Spearman correlations with OUTCOME: cortex, θ ρ = +0.138 (p = 0.582), DEMAND ρ = +0.674 (p = 0.0067), CAPACITY ρ = +0.459 (p = 0.083); striatum, θ ρ = +0.768 (p = 0.0027), DEMAND ρ = +0.806 (p = 0.0013), CAPACITY ρ = −0.097 (p = 0.720).

**Table 13.** Spatial association statistics (GSE233815).

| **Section** | **Day** | **Spots** | **θ–OUTCOME ρ** | **permutation p** | **CAPACITY–DEMAND ρ** | **permutation p** | **Reading** |
|---|---|---|---|---|---|---|---|
| C1-control | 0 | 2416 | −0.166 | 1.000 | +0.291 | 0.002 | null |
| B1-D1 | 1 | 2498 | +0.083 | 0.592 | +0.370 | 0.002 | null |
| D1-D3 | 3 | 2548 | −0.441 | 1.000 | +0.486 | 0.002 | null, inverse sign |
| C1-D7 | 7 | 2591 | −0.385 | 1.000 | +0.513 | 0.002 | null, inverse sign |
| D1-D7 | 7 | 1916 | −0.144 | 0.936 | +0.401 | 0.002 | null |

Spot-level correlations are within-section descriptive statistics.

### Chronic and aged brain: output shifts downstream, the brake does not tighten

In the rat 28-day cohort, CAPACITY rose by +0.992, the brake by +1.785, and a two-gene inactivation composite by +2.058 (Fig. 6A, B). The brake rose more than the machinery, but the two programs do not share a dispersion scale, so that is not an effect size; the data show co-induction, not that the brake alone limits clearance. This series is whole tissue and PECAM1 was not measured in it, so the six-gene set was scored on the five remaining genes; the same set scored without PECAM1 returns the same +1.785, because a gene that is absent from the matrix cannot enter the mean. The composition question is still open for the five genes that were measured: BRAKE tracks a myeloid marker score closely (Spearman rho = 0.826) and an endothelial score less so (rho = 0.684), and adjusting for both does not change the sign of the stroke coefficient, which at three model animals is a direction and not a partition. No brake-specific claim is made here. CD300A, the most tractable brake node, was induced (+2.46 VST), consistent with but not proof of a brake-dominant chronic state. Three model animals is too few for anything but effect sizes.

**Figure 5.**
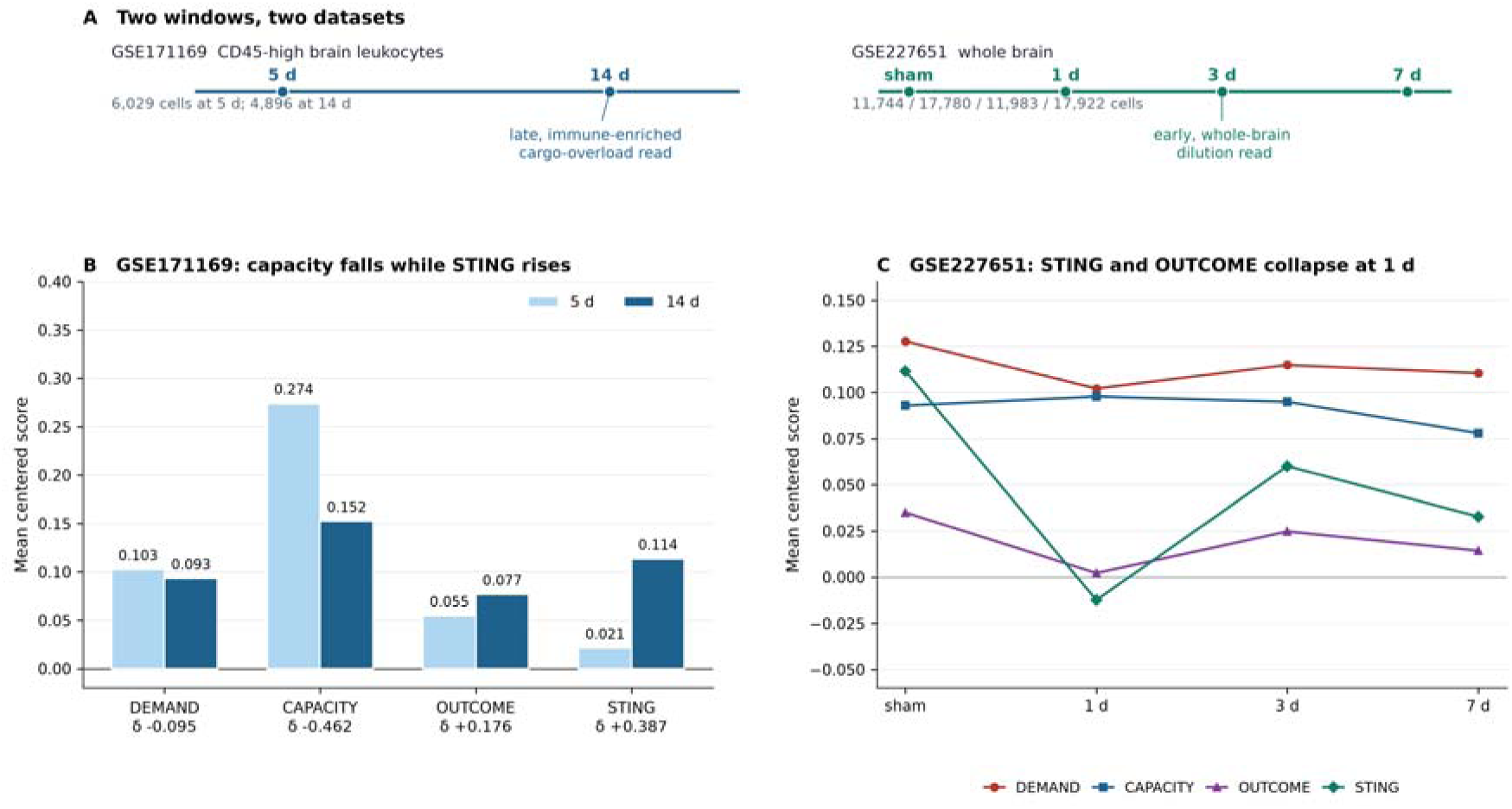
A late immunogenic-output state with STING co-movement in infiltrating leukocytes. (A) Design: GSE171169 (CD45-high cells at 5 and 14 days) and GSE227651 (whole brain at sham, 1, 3 and 7 days). (B) GSE171169: CAPACITY falls (δ = −0.462) while the STING subscore (δ = +0.387) and immunogenic output (δ = +0.176) rise. (C) GSE227651: STING is suppressed at day 1 and recovers only partially; whole-brain dilution prevents a myeloid-specific reading.

**Figure 6.**
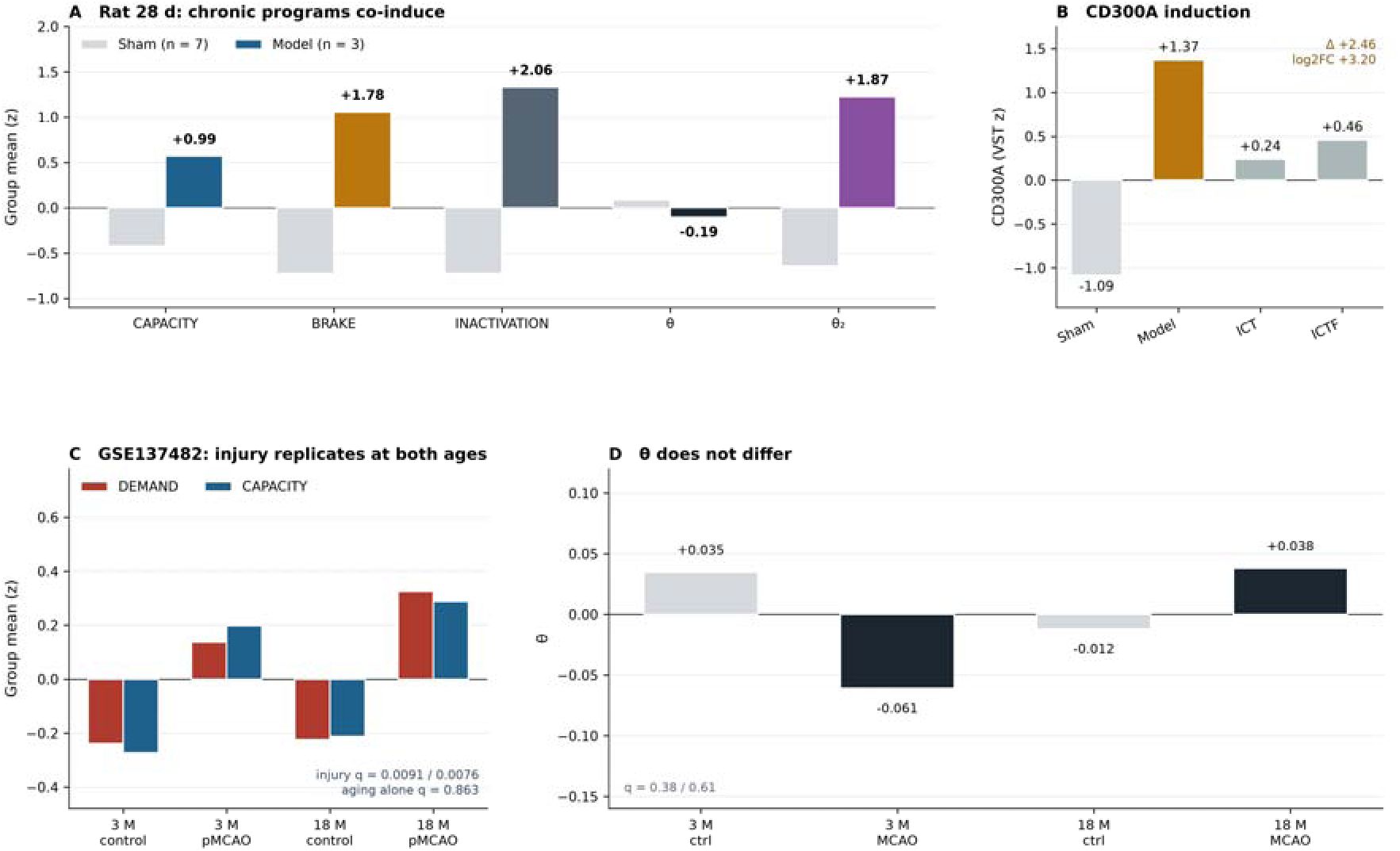
Chronic co-induction and the boundaries of the brake claim. (A) Rat 28-day cohort: CAPACITY (+0.992), BRAKE (+1.785) and INACTIVATION (+2.058) all rise, while θ remains closed (−0.189). (B) CD300A induction in the same cohort (+2.46 VST). (C) GSE137482 (3 versus 18 months): injury replicates at both ages, and ageing alone is null. (D) θ does not differ after injury at either age.

In the ageing series, the brake did not tighten with age in the two series whose libraries are deep enough to score. In GSE225948 (pseudobulked within compartment and animal), the brake moved +0.255 in aged microglia (q = 0.7041), −0.075 in aged monocyte-derived cells and −0.263 in aged endothelium; CAPACITY was likewise unchanged. In bulk cortex (GSE137482), the brake was +0.377 (q = 0.1210) and CAPACITY +0.090 (q = 0.0613). An age-related rise in inhibitory tone, the one age effect that would blunt a brake-directed intervention, is therefore unsupported in these two series; the day-14 recovery phase is not covered (Table 14).

**Table 14.**
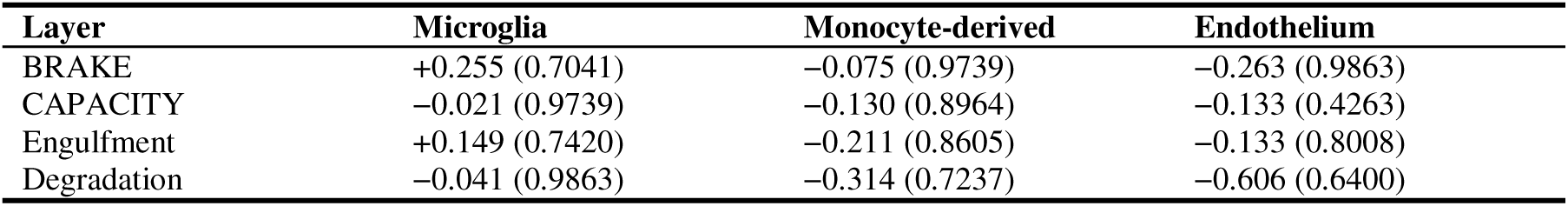
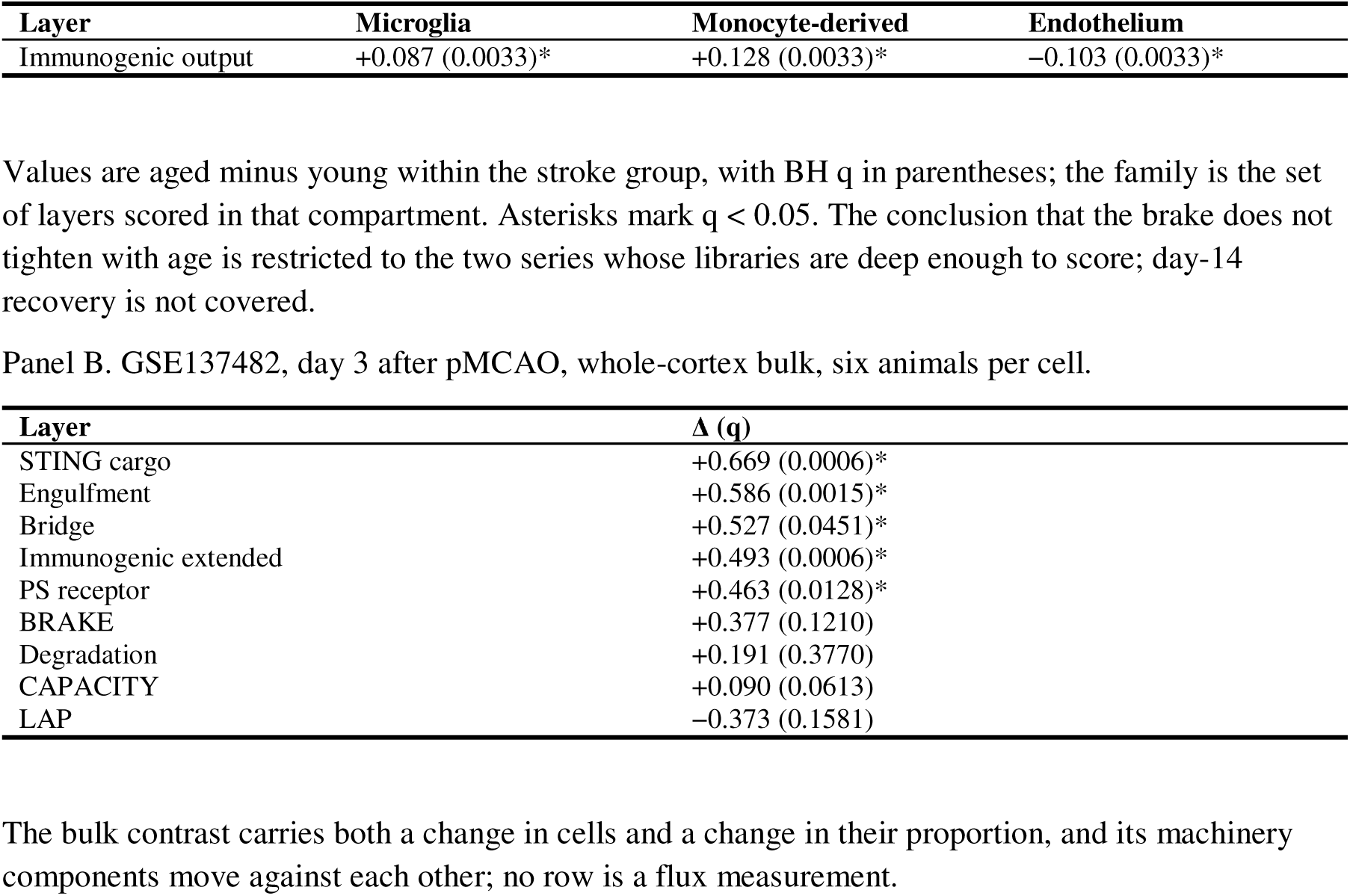
Ageing leaves the brake and the machinery unchanged in sorted cells.

| **Layer** | **Microglia** | **Monocyte-derived** | **Endothelium** |
|---|---|---|---|
| BRAKE | +0.255 (0.7041) | −0.075 (0.9739) | −0.263 (0.9863) |
| CAPACITY | −0.021 (0.9739) | −0.130 (0.8964) | −0.133 (0.4263) |
| Engulfment | +0.149 (0.7420) | −0.211 (0.8605) | −0.133 (0.8008) |
| Degradation | −0.041 (0.9863) | −0.314 (0.7237) | −0.606 (0.6400) |
| **Layer** | **Microglia** | **Monocyte-derived** | **Endothelium** |
| Immunogenic output | +0.087 (0.0033)* | +0.128 (0.0033)* | −0.103 (0.0033)* |
| Panel B. GSE137482, day 3 after pMCAO, whole-cortex bulk, six animals per cell. |  |  |  |
| **Layer** | **Δ (q)** |  |  |
| STING cargo | +0.669 (0.0006)* |  |  |
| Engulfment | +0.586 (0.0015)* |  |  |
| Bridge | +0.527 (0.0451)* |  |  |
| Immunogenic extended | +0.493 (0.0006)* |  |  |
| PS receptor | +0.463 (0.0128)* |  |  |
| BRAKE | +0.377 (0.1210) |  |  |
| Degradation | +0.191 (0.3770) |  |  |
| CAPACITY | +0.090 (0.0613) |  |  |
| LAP | −0.373 (0.1581) |  |  |

Values are aged minus young within the stroke group, with BH q in parentheses; the family is the set of layers scored in that compartment. Asterisks mark q < 0.05. The conclusion that the brake does not tighten with age is restricted to the two series whose libraries are deep enough to score; day-14 recovery is not covered.
The bulk contrast carries both a change in cells and a change in their proportion, and its machinery components move against each other; no row is a flux measurement.

What does move with age is the output end. Within compartment, the immunogenic set was higher in aged microglia (+0.087, q = 0.0033) and aged monocyte-derived cells (+0.128, q = 0.0033), and lower in aged endothelium (−0.103, q = 0.0033). Bulk cortex agreed in direction (+0.493, q = 0.0006). The largest age-associated difference was not on uptake at all: the cGAS–STING cargo set moved by +0.669 (q = 0.0006). Set beside the chronic data, this describes a sequence in which uptake is preserved, cargo-set transcripts rise, and output becomes more immunogenic. It is a reading, not a demonstration — no dataset measures cargo per cell — but it places the age-vulnerable part of the framework downstream of uptake.

### Where the claim stops: genetic, protein and compartment boundaries

Three boundaries were tested rather than asserted. The first is genetic. In GSE154396, Cd300a deletion did not raise CAPACITY, and degradation was lower in knockout macrophages (δ = −0.27 to −0.41 across 0, 1 and 3 h); the comparison has one sample per genotype and time, so it is descriptive, but it rules out a simple reading in which deleting the brake releases capacity at the transcript level. A brake can be released post-translationally without leaving a transcriptional trace, and this dataset can only answer the transcriptional question. A second genetic comparison in the day-3 sorted series (GSE141972) used TAK1-mKO rather than a brake deletion; layer scores were computed under the same locked null, and FDR-passing changes in the PS-receptor and immunogenic-core layers are recorded in the reproducibility inventory, but with three libraries per cell type and a signalling node distinct from CD300A these contrasts are context and are not pooled with the frozen framework contrasts or reported as framework effects.

The second is protein. Three proteomic series were examined as layer checks (Fig. 7; Table 15). Only one reaches 75% coverage for CAPACITY — PXD055588, mouse microglia sorted 24 h after tMCAO, 73 of 85 genes — and it deposits no per-sample values, so even the one proteome that meets the coverage requirement cannot be scored. The other two fall below that requirement (42/85 and 55/85) and give directional readouts only. Whole-tissue proteomics sees the degradative end of the machinery and is largely blind to the receptor and metabolic components. Formal protein–transcript concordance is therefore not estimable from these series; the only paired human material (GSE162955 with PXD022850, ρ = +0.32 across 1,834 proteins) supports tissue assignment but is too modest and composition-confounded to validate a program.

**Figure 7.**
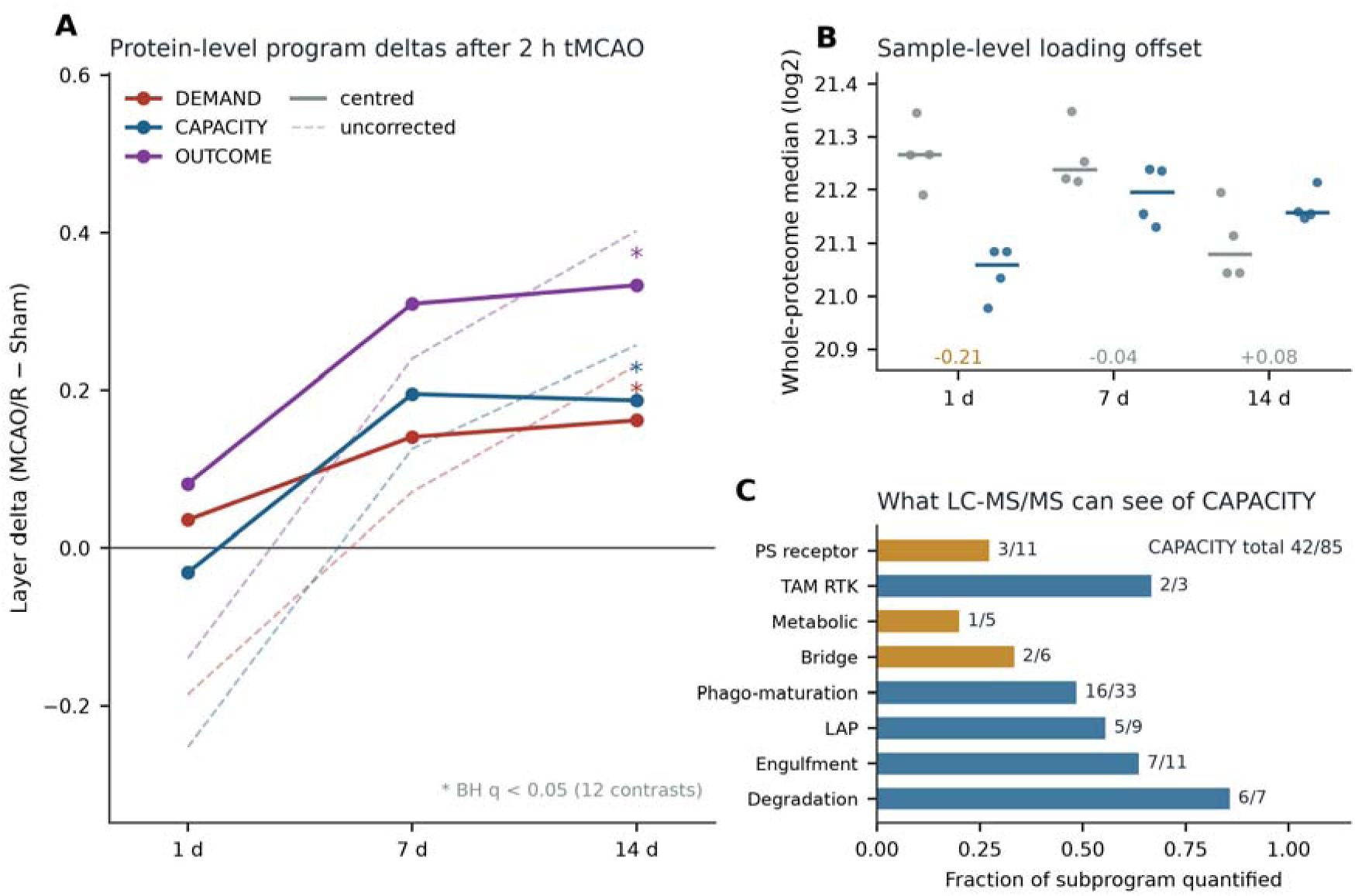
Protein-level readouts in rat cortex (PXD012151). (A) Layer differences for DEMAND, CAPACITY and OUTCOME at 1, 7 and 14 days, after subtracting each sample’s whole-proteome median. Asterisks mark contrasts surviving Benjamini–Hochberg across the twelve testable layer × time combinations (only the 14-day points). (B) Per-sample whole-proteome median by group. (C) Fraction of each CAPACITY subprogram quantified; the receptor and metabolic components are largely invisible to whole-tissue LC-MS/MS.

**Table 15.**
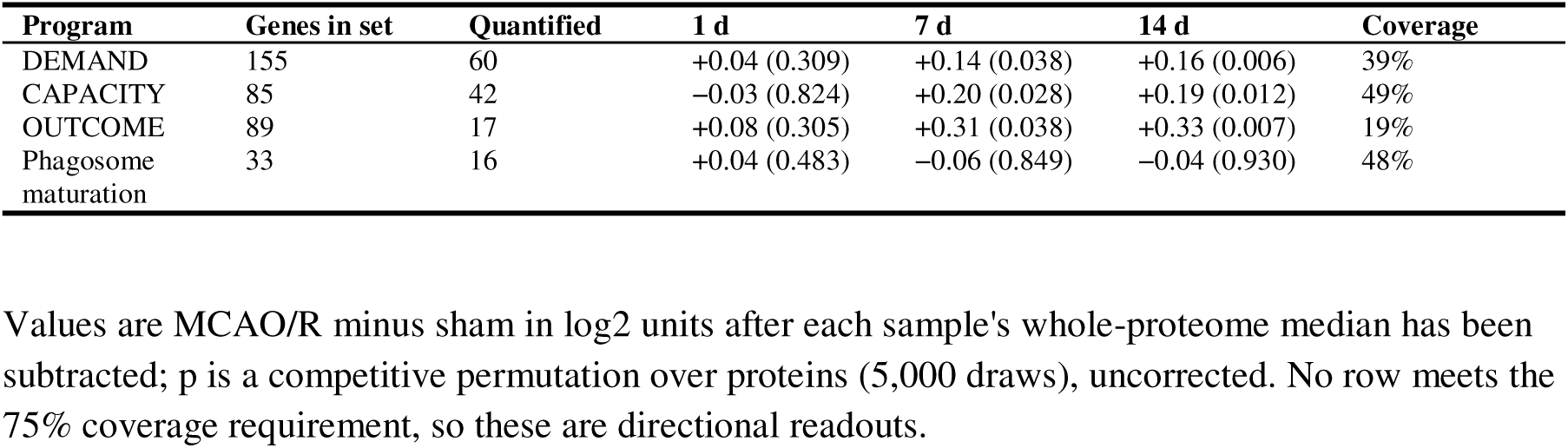
Protein-level readouts in rat cortex across three reperfusion times (PXD012151).

| **Program** | **Genes in set** | **Quantified** | **1 d** | **7 d** | **14 d** | **Coverage** |
|---|---|---|---|---|---|---|
| DEMAND | 155 | 60 | +0.04 (0.309) | +0.14 (0.038) | +0.16 (0.006) | 39% |
| CAPACITY | 85 | 42 | −0.03 (0.824) | +0.20 (0.028) | +0.19 (0.012) | 49% |
| OUTCOME | 89 | 17 | +0.08 (0.305) | +0.31 (0.038) | +0.33 (0.007) | 19% |
| Phagosome maturation | 33 | 16 | +0.04 (0.483) | −0.06 (0.849) | −0.04 (0.930) | 48% |

Values are MCAO/R minus sham in log2 units after each sample's whole-proteome median has been subtracted; p is a competitive permutation over proteins (5,000 draws), uncorrected. No row meets the 75% coverage requirement, so these are directional readouts.

The third is the late immune compartment. In CD45-high cells from GSE171169, CAPACITY fell from day 5 to day 14 (δ = −0.462) while the STING subscore rose (δ = +0.387) and OUTCOME rose (δ = +0.176), with DEMAND flat. The whole-brain series GSE227651 did not show the same switch — STING was suppressed at day 1 and recovered only partially. Both are compatible with a switch that requires persistent cargo, and both rest on two animals’ worth of cells per time point, so they are directions (Fig. 5).

## Discussion

### What the study shows

Four findings organise the study. First, the strongest positive signal — higher CAPACITY in whole blood after stroke — replicates in two cohorts and exceeds both expression-matched and detection-matched nulls, yet survives composition adjustment in only one cohort, cannot be identified in a third, and is absent in matched PBMCs. Second, human infarct core shows no coordinated loss of clearance transcripts; the tissue is disorganised, so a bulk transcriptome from it cannot be read as per-cell clearance. Third, in rodent brain CAPACITY is inducible, but the cell that carries it changes across snapshots, and energy supply moves before the machinery. Fourth, the one human series that sorts parenchymal cells resolves the composition half of the question and cannot test the capacity half: infiltrating myeloid cells expand and the opsonin set falls within them, while the depth spread leaves every microglial layer below its resolution limit. The program is real; what it means depends on where, in which cells, and when it is measured.

The argument does not rest on the direction of any adjusted coefficient. It rests on a difference in how the two blood cohorts behave under the same adjustment: the five proxies remove a quarter of the CAPACITY effect in one and nearly all of it in another, while in a third they predict the grouping almost perfectly (R² = 0.875). That this is not a uniform loss of precision is confirmed by the negative-control sets. It is also not a clean separation of composition from per-cell regulation. In GSE16561, CAPACITY tracks the array gradient at ρ = 0.814 and the joint model returns 77.7% attenuation with a residual coefficient of +0.109 (p = 0.043); in GSE58294, ρ = −0.008 and the joint model returns 26.10% against 26.68%. The cohorts differ in platform, probe set and sampling design, and we have not shown that a marker z-score means the same thing on both. Whether the difference in attenuation is biological or instrumental is therefore not identified here.

The negative results carry the same weight as the positive ones. θ adds no predictive value beyond DEMAND and is spatially null. The brake and the machinery co-induce in the chronic phase, so a brake-limited state cannot be inferred from their relative sizes. Age does not tighten the brake or expand the machinery; it changes what clearance produces. An efferocytic transcript score reports which program is transcribed in a sample of known composition at a known time. Until it is paired with a functional measurement, it says nothing about flux.

### Why a bounded measure is still useful

A measurement need not be a biomarker to be worth reporting. A high CAPACITY score in whole blood can mean more neutrophils, more monocytes, or more transcript per cell, and the composition adjustment says which is plausible in a given cohort. A high score in brain can mean more microglia, more macrophages, or a stronger program per cell, and the cell-type data say the answer changes between day 3 and day 7. A high score in the chronic phase can mean that clearance has failed or that it was induced together with its own brake.

Transcriptomic signatures are increasingly used to stratify patients, nominate drug targets and interpret animal models. Each use requires an explicit statement of what the score represents. We propose the five items in Table 16 as a reporting checklist, and Table 17 as a statement of what each design can establish. A score reported without these items cannot be compared across studies, whatever its p-value. The framework also explains why cross-species comparisons have disagreed: rat, mouse and human material differ in the cells present, the time window and tissue viability, so a conserved signature would be remarkable and its absence is expected.

**Table 16.** What a reportable efferocytic transcript score must specify.

| Element | What must be stated | Why it changes the reading | Example from this study |
|---|---|---|---|
| Compartment | Tissue, CSF, whole blood or PBMC, with species | Whole blood contains neutrophils that PBMC lack | CAPACITY +0.613 in whole blood; −0.054 in PBMC |
| Time since injury | Hours or days after onset or reperfusion | The cell carrying the program changes with phase | Macrophage-led at 48 h; microglial at day 3; macrophage at day 7 (three series, not one tracked cohort) |
| Cell composition | Measured fractions or validated proxies | A tissue score can rise because phagocyte number rose | Attenuation of 22.8% and 97.1% (upper bound; 77.7% with array gradient) in two blood cohorts |
| Unit of inference | Tissue, animal or cell | Cell-level intervals are not animal-level precision | Rat snRNA: one animal per severity and hemisphere |
| Program definition | Gene-set version, coverage and scale | Sets that mix clearance with output cannot separate them | OUTCOME shares no gene with DEMAND or CAPACITY |

None of these converts a transcript score into a measure of engulfment.

**Table 17.** What each design can establish about efferocytosis after stroke.

| **Design** | **What it measures** | **What it cannot separate** | **Use in this study** |
|---|---|---|---|
| Bulk tissue signature | Mean transcript level per sample | More phagocytes from more transcript per cell | Blood and brain cohorts |
| Lineage-proxy adjustment | Covariation with marker sets | Proxy scores are not cell fractions | GSE58294, GSE16561, GSE202518 |
| Sorted cells or single nuclei | Program within an annotated cell type | Capture depth, dissociation stress, replication | GSE250245, GSE86941, GSE171169 |
| Spatial transcriptomics | Co-localisation of programs | Cell identity at each spot; causal direction | GSE233815 |
| Functional uptake assay | Engulfment rate and cargo load | Which transcript sustains the rate | Not performed; proposed next step |
| Genetic perturbation | Necessity of a node | Transcript-level from protein-level release | GSE154396, GSE141972 |

The rows are complementary; a disagreement between two rows is often a difference in question rather than a contradiction.

### Relation to prior work

That efferocytosis is transcriptionally induced after stroke is reported repeatedly [8–10,13]. Yuan et al. nominated efferocytosis-related hub genes and a ceRNA network from bulk and single-cell data [9], and Xie et al. reviewed the mechanisms of efferocytosis in ischaemic resolution [8]. We reproduce induction in reperfused rodent brain and in whole blood. The difference lies in what the signal is taken to mean. Experimental efferocytosis has long counted a phagocytic index per cell rather than inferring uptake from the number of phagocytes in a lysate; our contribution is to show where that distinction can be recovered from transcript data across compartments, and where it cannot. In whole blood the proxy-driven attenuation spans 22.8% to 97.1% depending on cohort and model, and is not separately identified from the array gradient in the cohort with the largest value. A hub gene expressed by neutrophils will track neutrophil abundance in the same way.

Table S13 sets five prior reports against what this study can and cannot say about each, using a fixed vocabulary (reproduced, bounded, not tested, out of scope). No row is a refutation; the purpose is to make the difference between measurement frames explicit. The largest single gap it records is the protein layer: soluble TREM2 and MFG-E8 are prominent in the mechanistic literature [34–36] and are not measurable from the data assembled here.

The receptor literature provides a mechanistic frame that our data support only in part. CD300A blockade enhances efferocytosis by infiltrating myeloid cells and improves outcome after stroke [37], and CD300A has been implicated in ischaemic injury outside the brain [38]; Tufan et al. showed that pause-release can unleash macrophage capacity rapidly [39], a timescale snapshots cannot resolve. Deng et al. placed the constraint on sustained clearance in phagocyte metabolism, with adenosine A2A signalling engaging HIF1α-dependent glycolysis [40]. Our day-3 observation that oxidative phosphorylation and glycolysis rise before CAPACITY is consistent with that placement and extends it to a transcriptional ordering: the machinery is not the first mover, the energy that supports it is. Two further constraints sit outside our measurement frame: phagocytic capacity is saturable [41], which a transcript score cannot address by construction, and modulators such as the sigma-1 receptor [42] appear in no program scored here, so they are untested rather than rejected.

The negative genetic result is equally informative: Cd300a deletion did not raise CAPACITY, arguing against using CAPACITY as a proxy for the functional effect of a brake perturbation. And on whether microglia or infiltrating macrophages dominate clearance, the answer depends on time — at 48 h, macrophages and neutrophils lead on LAP; at day 3, microglia lead on composite CAPACITY; by day 7, the signal is macrophage-borne. A single-time-point comparison cannot adjudicate this, and studies naming one cell type as the efferocyte should state their time window.

### Implications

The clinical implication is mostly negative. CAPACITY is not a stand-alone biomarker: it did not separate stroke from control in the one PBMC cohort, it did not fall coordinately in infarct core, and in whole blood its adjusted effect is cohort-dependent. Whether the boundaries mapped here transfer to a clinical cohort is untested: the public series do not report stroke subtype, reperfusion status or a common sampling window in a way that would let us stratify them, so the compartment and time limits should be treated as properties of these datasets until a prospective cohort measures them. A trial using the score as a pharmacodynamic readout would need to specify compartment, time window and composition adjustment in advance, because a change in the score could be a change in the patients rather than their phagocytes. Stratification is a stronger demand than a group comparison, and a program whose split-half correlation is 0.445 in the striatum (0.616 after Spearman–Brown correction) cannot classify individuals.

The therapeutic implication is more specific. If the age- and chronic-phase effects lie downstream of uptake, then interventions that raise clearance capacity may not reach the step that matters. In the chronic phase the brake and machinery rise together, and CD300A is induced at the transcript level, but that induction has no human counterpart in the sorted parenchyma data and does not survive our own deletion test. In aged brain the brake does not tighten, but immunogenic output and STING cargo rise. Both lines point away from more capacity as a universal strategy and towards the handling of cargo and the control of output. Three predictions follow, each with a way to fail: a brake-directed intervention should reduce immunogenic output without requiring a rise in CAPACITY; a STING-directed intervention should retain efficacy in the chronic phase; an age-aware intervention should target cargo handling rather than uptake. Testing them requires engulfment and degradation measured directly (pHrodo or DQ-BSA uptake, phagosome maturation and cargo load) alongside the transcript score in the same animals, at more than one time point, with composition measured.

### Limitations

The study is observational and secondary: it re-analyses public datasets and two in-house cohorts and includes no prospective experiment in which clearance is manipulated and measured. The datasets differ in platform, species, tissue and time, which is a strength for mapping boundaries and a weakness for causal inference. No dataset carries more than five of the seven evidence layers, and the three the argument needs together — composition, compartment and time — are never jointly testable in any of them. The carrier sequence (48 h, day 3 and day 7) is read from three series under three injury models and three sorting schemes, so it is a cross-dataset synthesis and a hypothesis, not an order observed within one sample set (Supplementary Table S19).

Composition was addressed with marker-based proxies, not deconvolution. The proxies track cell abundance without being proportions, and the adjustment can absorb both composition and per-cell covariance. A reduction of 22.8% in one cohort and 97.1% in another, with the third unidentified, is consistent with a compositional explanation but does not prove one; the random-proxy null establishes specificity to the proxies, not identity with cell proportions. The two cohorts also differ in array platform, probe set and sampling design, and we have not verified that a marker z-score means the same thing across platforms. The gradient correlation differs sharply between them (ρ = 0.814 versus −0.008), which is consistent with a platform contribution to the difference. Whether the difference between 22.8% and 97.1% is biological or instrumental cannot be settled with these arrays. Reference-based deconvolution with a platform-matched signature matrix would be the next step; the public references available here did not permit it.

Cell-level data carry their designs’ limits. The rat single-nucleus libraries have one animal per severity and hemisphere — so depth gradient and hemisphere effect are fully nested and cannot be separated — and a mean of 7.8 CAPACITY genes per nucleus, which is why the cell-type ranking is a direction, not an estimate. The sorted-cell series are small (three or four libraries per group), which is why every such contrast carries a detection limit and why a non-significant result below that limit is labelled uninformative rather than negative. The sorted human parenchyma series has four donors against four and a 5.5-fold depth range, which is why its layer scores are reported against a resolution limit rather than as tests. The 28-day cohort has three model animals. The energy axis was run on six of the thirty datasets, and the 2 h mouse series — the earliest time point — has three sham animals and was not run, so whether supply has moved at 2 h is untested.

The recovery-phase question is unanswered. GSE142361 was the only series measuring young and aged microglia at day 14, and its aged libraries are too shallow (0.75/1.30 M reads against 4.18–5.77 M) to separate a loss of capacity from a global intensity deficit. Depth and age group are not completely confounded in that series: one aged library (A_S_N3, 5,124,213 reads) falls within the young range, and the correlation between age and log2 depth, though strong (r = −0.732), is not degenerate. With only eight libraries a covariate adjustment would be unstable; we therefore used the depth-matched and quantile-matched sensitivity series already computed, in which all ten aged-minus-young layer contrasts kept their sign (CAPACITY in sham −0.3877 to −0.3516, in MCAO −0.9448 to −0.9263), together with seven further variants of the contrast analysis. The ageing result is restricted to the two series whose libraries are deep enough to score, and does not extend to day-14 recovery. We regard this as the single largest gap in the study. The protein layer is incomplete in a way that cannot be repaired from the deposited files. Finally, the framework measures transcripts, not engulfment rate, lysosomal function or the number of uncleared apoptotic cells, and the output layer correlates at 0.93 to 0.94 with a standard inflammatory-response set, so its independence from inflammation is a statement about which genes it contains, not about what it measures. Neither the analysis plan nor the criteria for setting a dataset aside were pre-registered, which is why Table 1 lists every such dataset.

## Conclusions

Efferocytic transcriptional readouts after stroke are bounded by compartment, composition and time. CAPACITY is induced in reperfused rodent brain and in whole blood, but the blood signal is largely compositional under one covariate block in the cohort where the adjustment is decisive (97.1%, falling to 77.7% when the array gradient is included), attenuated by less than a quarter in another, and unidentifiable in a third; it was not detected in PBMCs and showed no coordinated change in human infarct core. In sorted human brain parenchyma, infiltration rises and the opsonin set falls within the infiltrating cells, but the depth of that series cannot resolve the capacity question. The demand– capacity difference adds no predictive value beyond demand, and in the chronic-phase series the brake and machinery co-induce while immunogenic output and cargo transcripts shift downstream; in the ageing series with adequate depth the brake does not tighten, a result that does not extend to day-14 recovery. These findings argue for reporting efferocytic transcripts as a bounded measurement with explicit covariates, and for testing therapeutic hypotheses against functional clearance rather than against the size of a transcriptional program.

## List of abbreviations

AIC: Akaike information criterion
BH: Benjamini-Hochberg
BRAKE, CAPACITY, DEMAND, INACTIVATION, OUTCOME: framework program scores named in Table 2
cGAS: cyclic GMP-AMP synthase
CI: confidence interval
FACS: fluorescence-activated cell sorting
FDR: false discovery rate
GEO: Gene Expression Omnibus
LFC: log2 fold change
MCAO: middle cerebral artery occlusion
PBMC: peripheral blood mononuclear cell
PRIDE: PRoteomics IDEntifications database
sMCAO/mMCAO: spontaneous/reperfused middle cerebral artery occlusion as labelled in the source datasets
STING: stimulator of interferon genes
VST: variance stabilising transformation.

## Declarations

### Ethics approval and consent to participate

Every human analysis in this manuscript is a reanalysis of previously published, de-identified data that is openly available under the accessions listed in Availability of data and materials; no new human participants were recruited and no new human samples were collected, so no additional ethics approval or consent was obtained for the work reported here. The ethics approval and consent procedures that cover each human dataset are those reported by its original study. The two in-house rodent cohorts (the mouse P1/P2 focal ischaemia series and the rat 28-day cortical ischaemia cohort) were generated under institutionally approved protocols. Committee name and approval reference numbers: Biomedical Research Ethics Committee of Gannan Medical University /Approval No.: 2022201).

### Consent for publication

Not applicable.

## Availability of data and materials

The datasets supporting the conclusions are available in the Gene Expression Omnibus (GEO) repository: GSE105132, GSE137482, GSE137595, GSE141972, GSE154396, GSE162955, GSE163654, GSE16561, GSE171169, GSE174574, GSE202518, GSE22255, GSE225948, GSE227651, GSE23160, GSE233815, GSE23651, GSE250245, GSE267023, GSE318800, GSE319238, GSE331114, GSE332910, GSE337578, GSE345807, GSE37587, GSE58294, GSE86941. The full list with platform, species, tissue and disposition is in Supplementary Table S2. Proteome data are in the PRIDE Archive: PXD012151, PXD016538, PXD022850, PXD025077, PXD055588. Eight series were scored or inspected and then withdrawn or downgraded (GSE110141, GSE142361, GSE204702, GSE248793, GSE285659, GSE294912, GSE56267, GSE64811); the reason for each is in Table 1, and two of them (GSE64811, GSE110141) still contribute one number each. The two in-house rodent series have no public accession at the time of submission, are available from the authors on reasonable request during peer review, and will be deposited in GEO on acceptance. Analysis code, the frozen framework gene sets, per-dataset coverage tables and the dataset manifest are archived in the Zenodo record under doi:10.5281/zenodo.24415702 (project name: efferocytosis framework; project home page: https://zenodo.org/records/24415702; archived version: 1.0.1; operating system: platform independent; programming language: Python and R; other requirements: Python 3.12.13 and R 4.3.3 with scanpy, limma and edgeR, with all further package versions listed in docs/ENVIRONMENT.md inside the archive; license: MIT for the analysis code and CC BY 4.0 for the framework gene sets, with six KEGG-derived membership files under Kanehisa Laboratories’ terms; non-academic use of those six files requires a commercial licence from Kanehisa Laboratories, and no other restriction applies to non-academic use). No individual participant data appear in any file.

## Authors’ contributions

S. D., Z.H., and Z.Y. conceived and designed the framework; S. D. and Y. Z. assembled and curated the datasets; S. D., Y. L., and Y. Z. implemented the scoring, null-distribution and composition machinery; S. D., and Z.Y. ran the analyses and generated every figure and table from the tabulated results; Y. L., Z.H., and Z.Y. discussed and drafted the manuscript; Z.Y. and Z.Y. supervised the study, and critically revised the manuscript. All authors reviewed and approved the final version of the manuscript and agree to be accountable for all aspects of the work.

## Acknowledgements

We really appreciate the researchers who submitted and shared their research data to the GEO database and PRIDE Archive.

## Additional files

**Additional file 1.** Format: .docx. Title: Supplementary Information. Description: Tables S1–S19 — framework gene-set membership with source map (S1), file checksums in MANIFEST.md5 in the Zenodo code archive, dataset inventory with disposition (S2), program-level effect sizes (S3), per-dataset coverage (S4), operational phase classification (S5), composition adjustment (S6), cell-type-specific deltas (S7), claims-to-evidence grade (S8), cross-cohort concordance and PBMC equivalence (S9), expression-matched controls (S10), detection limits (S11), design limit of every layer by contrast in GSE86941 (S12), contrast with prior reports (S13), probe-aggregation sensitivity (S14), GO-only reconstruction of the vote-built output program (S15), detection-matched null (S16), scope of the energy axis (S17), the family over which each correction step was applied with the three null constructions side by side (S18), and dataset-by-dataset coverage of the seven evidence layers (S19). This version: 2026-11-22, nineteen tables.

