## Additional file 1 for "Bounded by compartment, composition and time: efferocytic transcriptional readouts after ischaemic stroke"

Supplementary Information

**Efferocytic Transcriptional Programs After Stroke Are Compartment- and Composition-Dependent: A Three-Layer Framework and Its Limits**

This file contains the supplementary note and Tables S1–S19. The tables cited in the main text are S1, S2, S13, S14, S16, S17 and S19. All program scores use the 2026-10-21 framework build (L2_CAPACITY = 85 genes in all three species; CD300LF reassigned to EXT_CONTEXT_DEPENDENT) and the 2026-10-22 analysis freeze. Numeric cells were generated from the result files named in each table; dataset metadata were cross-checked against Methods §4.2 and dataset_catalog.md.

### Supplementary Note S1. Context datasets: design and per-cell outputs

Eight context datasets were scored under the same framework build and are reported in §2.10. None is carried into a primary conclusion. Per-cell program scores are archived in results/newdata/ (gse154396_paper_celltype_scores.csv, gse332910_celltype_scores.csv, gse345807_ratday21_percell_programs.csv, gse267023_pbmc_percell_programs.csv, gse337578_day3spatial_scores.csv, gse318800_percell_programs.csv); bulk context scores are in results/gse23651/ and results/gse37587/. Effect sizes in §2.10 are Cliff's δ; no p-values are reported for datasets without biological replication.

#### S1.1 Genetic brake read: GSE154396

Mouse Cd300a^fl/fl versus LysM-Cre CD45+ brain cells at 0, 1 and 3 h after MCAO-reperfusion (51,005 cells; 10/10 programs passed the ≥75% coverage gate). The GEO description specifies 0 h after reperfusion, so this is a reperfusion reference rather than a true naive group. Lyz2-Cre recombines efficiently in blood monocytes and macrophages but only partially in resident microglia, so the microglial contrast mixes deleted and undeleted cells. CD300A expression fell in infiltrating myeloid cells (pooled Cliff's δ = −0.26; 1 h −0.35), less so in microglia (δ = −0.10), and the untargeted paralogue CD300LF was unchanged (δ −0.01 to +0.08). CAPACITY did not rise at any timepoint (Δ +0.044/−0.053/−0.004 across all cells; |δ| < 0.15), and the lysosomal-degradation module was lower in cKO macrophages (δ = −0.27/−0.41/−0.29). With one pooled sample per condition, this dataset confirms compartment-specific deletion but cannot establish a capacity-program response; whether CD300a removal enhances engulfment requires measured phagocytosis with verified recombination.

#### S1.2 Animal-level time course: GSE332910

Mouse striatal MCAO/R snRNA at sham/0 h/24 h/7 d/14 d, n = 3 animals per group (235,711 cells; 10/10 coverage). Per-cell scores were averaged within each library, and the three sham animals were compared with the three animals at each post-injury timepoint by exact two-sided permutation of animal labels. With 3 versus 3 animals the smallest attainable two-sided p is 0.10, so the design can detect only complete 3/3 separation. At 0 h, pseudobulk DEMAND rose by +0.133 and CAPACITY by +0.144, leaving θ at −0.011; OUTCOME rose by +0.070 and STING by +0.027, and each of these four comparisons separated 3/3 (exact p = 0.10), whereas θ did not (p = 0.30). The pattern repeated at 24 h (+0.076/+0.082; θ −0.006; STING +0.031; p = 0.10 except θ p = 0.70) and at 14 d (+0.071/+0.090; θ −0.019; STING +0.026; p = 0.10). At 7 d one sham animal sat above one injured animal and no program separated the groups (p = 0.40–0.70). Microglial θ values (−0.051 at 7 d, −0.015 at 14 d) are cell-level descriptive summaries without animal-level intervals. This is the only additional series with animal-level replication; its pseudobulk read shows balanced co-upregulation rather than demand outrunning capacity, so it bounds rather than confirms the θ-opening claim.

#### S1.3 Rat day-21 chronic window: GSE345807

Sham n = 2 versus MCAO n = 4 (72,018 cells). All nine framework sets compared at the cell level moved in the same direction as the in-house 28-day series; CAPACITY δ = +0.362 and STING δ = +0.223. The pseudobulk comparison was concordant for all eight programs compared (CAPACITY δ = +1.000). With two sham animals the analysis is descriptive.

#### S1.4 Spatial context: GSE337578

One SeekSpace sample at day 3 (30,560 spots). CAPACITY and DEMAND were spatially co-localized (ρ = +0.894; torus-shift p = 0.001 within this section), while CAPACITY showed no global autocorrelation (Moran's I ≈ 0.0001). The shipped spot score subtracts one global scalar rather than each spot's own mean, so it retains a within-spot expression term: the shipped CAPACITY and DEMAND scores each correlate with the log framework-gene count of the spot at ρ = +0.947 and +0.956, larger than their mutual correlation. Because the denominator of that count contains the tested sets themselves (243 of 841 framework symbols = CAPACITY 85 ∪ DEMAND 159), those two correlations are an upper bound on the depth contribution rather than a clean depth coefficient. Re-scoring the same section under two centring variants and after depth residualisation gives ρ = +0.817 (all-gene centring, denominator independent of the tested sets), +0.702 (depth-residualised shipped score) and +0.162 (framework-gene centring) — results/newdata/gse337578_day3spatial_score_calibre.csv, script 381. The framework-centred number conditions on a variable that is itself 29% composed of the sets under test, so it is over-correction and is not a fair comparator. Under the two principled adjustments most of the co-localization survives (0.82 and 0.70 against the shipped +0.89), so we read this section as depth-confounded but with the co-localization largely preserved rather than as convention-dependent. Radial band means are flat for CAPACITY (0.079 → 0.090 across inner/mid/outer) but rise outward for L1 subprograms (apoptotic burden 0.009 → 0.014; PS_EXPOSURE 0.057 → 0.132), consistent with a capture-depth gradient rather than regional amplification. This is a single section, so it cannot support slice-level inference in any case.

#### S1.5 Human PBMC single-cell series: GSE267023

Eight patients, 77,844 cells, 10x GEX only. The deposit contains no cell-type annotation, so the prespecified myeloid-versus-lymphoid per-cell contrast is not computable. This is a design limitation, not a null result.

#### S1.6 Htr2b myeloid cKO: GSE318800

n = 2 per group, 63,181 cells. The driver is Ms4a3-Cre, which targets granulocyte–monocyte progenitors and labels blood-derived macrophages and monocyte-derived cells; it does not target yolk-sac-derived resident microglia. The deposited annotation labels microglia and monocyte-derived cells, but a "microglia" label in a Ms4a3-Cre pedigree cannot be read as cell-autonomous microglial deletion. cKO cells had a lower median library size (1,716 versus 2,343 UMIs), and all nine programs shifted downward, so absolute deltas are confounded by a global technical shift. Within-cell θ was higher in the cKO "microglial" bin (Δ = +0.085; animal-level pseudobulk +0.088), but with two animals per group, a non-microglial driver and a global shift this is a correlate, not a cell-autonomous effect.

#### S1.7 Acute rat cortex: GSE23651

Sham/vehicle/ZM447439 at 8 and 24 h, n = 4 per design cell (24 samples, GPL6101). At 24 h, vehicle minus sham gave CAPACITY +0.198 (δ = +0.625) and the two-gene INACTIVATION co-expression composite +1.641. BRAKE was −0.563 but covered only 4/6 genes (66.7%), below the 75% scoring gate; BRAKE and constructed θ₂ comparisons are therefore exploratory and do not enter a primary conclusion. ZM447439 minus vehicle at 24 h gave θ₂ −0.487 (δ = −0.750), exploratory BRAKE +0.601 and CAPACITY −0.010 (δ = −0.125). CAPACITY coverage was 77/85 (91%); the eight missing genes are absent from the GPL6101 manifest, so 91% is the platform ceiling rather than an annotation gap. With four animals per cell and no exact permutation p above the 0.029 floor, only effect sizes are reported.

#### S1.8 Within-patient whole-blood time course: GSE37587

Thirty-four patients sampled at 0–24 h and 24–48 h (68 samples, GPL6883). Paired changes were DEMAND +0.044 (p = 0.027), CAPACITY +0.011 (p = 0.698), θ +0.033 (p = 0.343), θ₂ −0.329 (p = 0.168), OUTCOME −0.096 (p = 0.031) and STING −0.091 (p = 0.029); BRAKE and INACTIVATION did not change (p = 0.309 and 0.153). Coverage was 84/85 for CAPACITY, 167/178 for DEMAND, 101/108 for OUTCOME and 36/38 for STING. Benjamini–Hochberg correction across these eight paired metrics, computed from the same file (results/gse37587/gse37587_paired_deltas_bh.csv, column q_bh_8metric_family), gives q = 0.084 for DEMAND, OUTCOME and STING and q ≥ 0.27 for the rest, so no metric reaches q < 0.05 and the three nominally significant shifts are reported as directional only. The interval is within-patient, so it is not a case–control contrast and does not enter a primary claim.

#### S1.9 Sensitivity of the CARGO phase rule

L4_STING is almost entirely contained within L3_OUTCOME (36/38 human, 39/41 mouse, 21/23 rat genes; only IFNB1 and TICAM1 are STING-specific), so co-movement is partly definitional. The operational phase margin (+0.05 for DEMAND, STING and OUTCOME) is half the smallest between-group difference interpreted in the main text and is not a validated cut-off. It classifies GSE171169 at 14 d and the 28-day rat Model/ICT/ICTF groups as phase II; baseline samples are phase I and most post-injury samples are intermediate (Table S5). θ does not track the call, so the phase labels are descriptive.

#### S1.10 Earliest window, four-arm design: GSE137595

Mouse cortex 2 h after 90 min MCAO with reperfusion, ipsilateral and contralateral hemisphere of both ischaemic and sham animals (Clariom S, GPL24242; 24 samples, 8/8/4/4). This is the only dataset in the project that crosses injury status with hemisphere, so it can separate an ischaemia effect from an animal effect and can carry its own negative control (the sham ipsilateral-versus-contralateral contrast). All fourteen framework layers passed the 75% coverage gate (CAPACITY 82/85, DEMAND 158/159, OUTCOME 107/110, BRAKE 6/6, STING 39/41; results/newdata/gse137595_coverage.csv).

Two animals (one ischaemic, one sham) were removed before scoring because their four samples fell more than three robust standard deviations below the cohort median on two quality indicators that are independent of the principal components of variation — the median intensity of twelve housekeeping genes and the sample mean expression. Removal was at the animal level, so the ipsilateral–contralateral pairing was preserved; 20 samples remained (7/7/3/3). Because a program score is the mean of member-gene z-scores computed over the whole sample pool, four systematically low-intensity samples distort the z scale of every program (the within-animal ipsilateral–contralateral correlation of CAPACITY reaches r = 0.991 in the ischaemic arm on the full pool), so quantile normalisation, per-gene z-scoring and scoring were re-run on the retained samples rather than deleting columns from an already-scored matrix (results/newdata/gse137595_diagnostics.json).

Results. On the ischaemic-versus-sham axis, θ rose by +0.196 (Welch p = 0.0024; BH q = 0.009 across the four contrasts), the only contrast whose θ exceeds the 95th percentile of 2,000 expression-matched random gene sets (null q95 = 0.174; one-sided p = 0.031). DEMAND rose by +0.239 and also exceeded its null (one-sided p = 0.002; null q95 = 0.129); CAPACITY did not move (+0.043; p = 0.66) and did not exceed its null (one-sided p = 0.254). Within ischaemic animals the ipsilateral hemisphere did not differ from the contralateral one (CAPACITY +0.045, p = 0.53; DEMAND +0.090, p = 0.170), and the sham negative control showed no hemisphere difference either (CAPACITY +0.065, p = 0.65; DEMAND −0.052, p = 0.64), both inside the random-set null. A difference-in-differences test returned an indeterminate verdict for all eight scored programs (CAPACITY DiD −0.021, 95% CI −0.452 to +0.411, p = 0.89; DEMAND DiD +0.142, 95% CI −0.178 to +0.462, p = 0.27), so the 2 h difference is a hemisphere-level difference between ischaemic and sham animals and cannot be assigned to the ischaemic hemisphere itself. Depth confounding was the largest at |ρ| = 0.747 (DEMAND) and residualisation did not preserve the effects. All 16 CD300-family transcripts were detected; Cd300a did not move in any of the four contrasts (ischaemic versus sham ipsilateral −0.543, p = 0.108).

Limits. There is no drug arm, so this dataset says nothing about NBP or icaritin. The 2 h window is earlier than the 1 d and 28 d datasets and is deliberately not placed on the same time axis. The companion proteome PXD016538 reaches 21/85 CAPACITY genes (24.7%), far below the 75% gate, so the protein layer is not estimable here.

#### S1.11 Five-point protein time course: PXD025077

Mouse cortex after distal hypoxic MCAO at 1, 3, 7, 14 and 28 days (TMT 10-plex, seven plexes; 6 versus 6 animals at days 1, 3 and 7, 6 versus 6 at day 14, 4 versus 2 at day 28). The TMT channel assignment was randomised within each plex and not deposited, so the reporter channels cannot be mapped to groups and the contrasts cannot be recomputed; layer differences are read from the authors' supplementary per-protein table, which lists the 3,625 proteins that changed against both controls and is therefore a pre-filtered subset of the 8,068 quantified protein groups. Coverage is 55 of 85 CAPACITY genes (64.7%), 86 of 159 DEMAND genes and 16 of 33 phago-maturation genes, so no layer clears the 75% gate and nothing here is called a score. The layer means are computed on the members present in that table: 32 of the 55 CAPACITY genes and 50 of the 86 DEMAND genes. Because those subsets differ in size and composition, their difference is not the θ of §4.3 and no θ is reported for this series.

Four checks accompany the estimate (results/proteomics/pxd025077_readiness_verdict.json). First, the whole-proteome background is reported alongside, so a layer shift can be separated from a global one: the background mean over all proteins with a value at that timepoint is +0.040 at day 1 and +0.051 at day 7, so the day-7 layer peak is not a proteome-wide shift. Second, layer means are tested against a size-matched null drawn from the same selection domain (random layers of the same size from the proteins quantified and present in the table at that timepoint; 5,000 draws, seed 20261024). The CAPACITY layer exceeds the 95th percentile at day 7 (+0.789 against +0.225; one-sided p = 0.0002) and does so at days 3, 14 and 28, but not at day 1 (+0.028 against +0.150). Third, the estimate is recomputed with the phagocyte markers present in the layer removed (C1QA, C1QB, C1QC, STAB1, TREM2): the day-7 mean falls from +0.789 to +0.602 and the peak does not move, while the same exclusion leaves DEMAND unchanged at every time point, so the exclusion is not simply deleting a signal. Fourth, peak timing is assessed against the peak-timepoint distribution of the whole table: 17 of 32 CAPACITY members reach their own maximum at day 7, against a background distribution whose largest group is day 1 (hypergeometric p = 7.5×10⁻⁵). The authors' own four-cluster assignment, made without reference to this framework, independently places 10 of the 32 CAPACITY members in their later-up cluster against 1.7 expected (p = 3.0×10⁻⁶).

The composition question is bounded rather than closed. Per-cell CAPACITY in border-associated macrophages in our own single-cell cohorts is −0.131 at day 7 and −0.177 at day 14, the opposite sign to the whole-tissue protein layer, which measures total tissue mass rather than a within-cell quantity. Using the phagocyte (microglia plus border-associated macrophage) share as a ceiling on what infiltration alone could contribute, the day-7 rise is larger than the ceiling allows: the phagocyte share changes 1.15-fold between sham and day 7 in the day-7 cohort (2.46% to 2.83%), bounding the composition contribution at +0.202 log2 units against an observed +0.789, so at most a quarter of the day-7 signal can be attributed to more phagocytes in the tissue. At day 14 the same ceiling is +0.517 against an observed +0.495, so the day-14 value is fully explicable by infiltration and carries no independent information. This ceiling is a cross-dataset comparison — the single-cell cohorts are transient occlusion, the protein series is distal hypoxic occlusion — and it is a ceiling rather than an estimate of the contribution.

Limits. No drug arm, so nothing about NBP or icaritin. Day 28 has two sham animals. The contrasts are read from a pre-filtered table rather than re-derived, so the magnitudes are conditional on that selection. The occlusion model differs from the tMCAO-with-reperfusion series used elsewhere in this paper, so the five-point trajectory is not placed on the same time axis as the 2 h or 28 d transcript datasets. The series is not among the 34 scored datasets.

### Table S1. Framework gene-set membership

Set sizes under the frozen build. Full long-format membership (species, program, gene) is provided in supplement/Table_S1_gene_set_membership.tsv and in the frozen GMT files results/io_framework/io_framework_{human,mouse,rat}.gmt.

| **Program** | **Human** | **Mouse** | **Rat** |
| --- | --- | --- | --- |
| L1_APOPTOTIC_BURDEN | 178 | 159 | 155 |
| L1_PS_EXPOSURE | 6 | 6 | 6 |
| L1_FIND_ME | 7 | 7 | 7 |
| L2_CAPACITY | 85 | 85 | 85 |
| L2_ENGULF | 11 | 11 | 11 |
| L2_LAP | 9 | 9 | 9 |
| L2_PHAGO_MATUR | 33 | 33 | 33 |
| L2_DEGRADATION | 7 | 7 | 7 |
| L2_METABOLIC | 5 | 5 | 5 |
| L2_DONT_EAT_ME | 6 | 6 | 6 |
| L2_BRIDGE | 6 | 6 | 6 |
| L2_PS_RECEPTOR | 11 | 11 | 11 |
| L2_TAM_RTK | 3 | 3 | 3 |
| L2b_INACTIVATION | 2 | 2 | 2 |
| L2_MACHINERY_FULL | 85 | 85 | 85 |
| L3_RESOLUTION_OUTPUT | 5 | 5 | 5 |
| L3_IMMUNOGENIC_CORE | 137 | 138 | 114 |
| L3_IMMUNOGENIC_DECOUPLED | 108 | 110 | 89 |
| L4_PYROPTOSIS | 57 | 69 | 43 |
| L4_STING_CARGO_OVERLOAD | 38 | 41 | 23 |
| L0_NECROPTOSIS_EXECUTOR | 24 | 24 | 24 |

Membership of the small sets (all members listed; larger sets are in the TSV):

| **Program** | **Human** | **Mouse** | **Rat** |
| --- | --- | --- | --- |
| L1_PS_EXPOSURE | ANO6, XKR4, XKR6, XKR7, XKR8, XKR9 | ANO6, XKR4, XKR6, XKR7, XKR8, XKR9 | ANO6, XKR4, XKR6, XKR7, XKR8, XKR9 |
| L1_FIND_ME | CX3CL1, CX3CR1, P2RY2, PANX1, S1PR1, SPHK1, SPHK2 | CX3CL1, CX3CR1, P2RY2, PANX1, S1PR1, SPHK1, SPHK2 | CX3CL1, CX3CR1, P2RY2, PANX1, S1PR1, SPHK1, SPHK2 |
| L2_ENGULF | ARPC5, CDC42, CRK, DNM2, DOCK1, ELMO1, GULP1, ITGAV, ITGB3, ITGB5, RAC1 | ARPC5, CDC42, CRK, DNM2, DOCK1, ELMO1, GULP1, ITGAV, ITGB3, ITGB5, RAC1 | ARPC5, CDC42, CRK, DNM2, DOCK1, ELMO1, GULP1, ITGAV, ITGB3, ITGB5, RAC1 |
| L2_LAP | ATG16L1, ATG3, ATG5, ATG7, BECN1, MAP1LC3B, PIK3C3, RAB5C, RUBCN | ATG16L1, ATG3, ATG5, ATG7, BECN1, MAP1LC3B, PIK3C3, RAB5C, RUBCN | ATG16L1, ATG3, ATG5, ATG7, BECN1, MAP1LC3B, PIK3C3, RAB5C, RUBCN |
| L2_DEGRADATION | CTSB, CTSD, CTSL, DNASE2, LIPA, SMPD1, TCIRG1 | CTSB, CTSD, CTSL, DNASE2, LIPA, SMPD1, TCIRG1 | CTSB, CTSD, CTSL, DNASE2, LIPA, SMPD1, TCIRG1 |
| L2_METABOLIC | ABCA1, NR1H2, NR1H3, PPARD, PPARG | ABCA1, NR1H2, NR1H3, PPARD, PPARG | ABCA1, NR1H2, NR1H3, PPARD, PPARG |
| L2_DONT_EAT_ME | CD24, CD300A, CD47, PECAM1, SIGLEC10, SIRPA | CD24A, CD300A, CD47, PECAM1, SIGLECG, SIRPA | CD24, CD300A, CD47, PECAM1, SIGLEC10, SIRPA |
| L2_BRIDGE | C1QA, C1QB, C1QC, GAS6, MFGE8, PROS1 | C1QA, C1QB, C1QC, GAS6, MFGE8, PROS1 | C1QA, C1QB, C1QC, GAS6, MFGE8, PROS1 |
| L2_PS_RECEPTOR | ADGRB1, CD36, HAVCR1, HAVCR2, LRP1, MEGF10, STAB1, STAB2, TIMD4, TREM2, TYROBP | ADGRB1, CD36, HAVCR1, HAVCR2, LRP1, MEGF10, STAB1, STAB2, TIMD4, TREM2, TYROBP | ADGRB1, CD36, HAVCR1, HAVCR2, LRP1, MEGF10, STAB1, STAB2, TIMD4, TREM2, TYROBP |
| L2_TAM_RTK | AXL, MERTK, TYRO3 | AXL, MERTK, TYRO3 | AXL, MERTK, TYRO3 |
| L2b_INACTIVATION | ADAM17, TIMP3 | ADAM17, TIMP3 | ADAM17, TIMP3 |
| L3_RESOLUTION_OUTPUT | IL10, PTGER2, PTGER4, PTGS2, TGFB1 | IL10, PTGER2, PTGER4, PTGS2, TGFB1 | IL10, PTGER2, PTGER4, PTGS2, TGFB1 |

### Table S2. Dataset inventory

Thirty-six public accessions and two in-house cohorts (38 dataset rows; GSE174574 and GSE319238 form one integrated analysis). Thirty contribute a number to the results and eight were scored or examined and then withdrawn or downgraded, each with its reason in Table 1 and in the Disposition column below. “Discovery” denotes datasets used for a primary claim; “context” denotes boundary conditions.

| **#** | **Dataset** | **Species** | **Tissue** | **Samples / cells** | **Assay** | **Role** | **Disposition** |
| --- | --- | --- | --- | --- | --- | --- | --- |
| 1 | P1/P2 (in-house) | Mouse | Brain, ischemic tissue | 19 samples after exclusion (ICT/NBP/tMCAO/Sham, 1 d) | Bulk RNA-seq | Discovery/context; direction only (n=2–3/group) | contributes |
| 2 | GSE250245 | Rat | Cortex, ipsi/contra | 6 libraries; 73,816 nuclei | snRNA-seq | Cell-type CAPACITY response | contributes |
| 3 | GSE16561 | Human | Whole blood | 39 stroke / 24 control donors | Microarray (GPL6883) | Discovery cohort | contributes |
| 4 | GSE58294 | Human | Whole blood | 92 samples / 46 subjects (23 patients × 3/5/24 h; 23 controls) | Microarray (GPL570) | Replication cohort; subject-level inference | contributes |
| 5 | GSE22255 | Human | PBMC | 20 stroke / 20 control, age- and sex-matched | Expression array | Matched PBMC control | contributes |
| 6 | GSE163654 | Mouse + rat | Cortex | n=3/group/species; pMCAO 2 and 6 h | Microarray | Cross-species rank order plus gene-level sign concordance (script 103) | contributes |
| 7 | GSE162955 | Human | Brain, infarct core vs contralateral cortex | 6 paired patients (12 samples) | Microarray (GPL17586) | Boundary-condition comparison | contributes |
| 8 | GSE23160 | Mouse | Cortex + striatum | n=4/group; MCAO 2 h + reperfusion 2/8/24 h | Microarray | Temporal θ–OUTCOME association; incremental validity | contributes |
| 9 | GSE233815 | Mouse | Brain, Visium | 5 slices (control/D1/D3/D7×2); ~2,500 spots/slice | Spatial transcriptomics | Spot-level spatial context | contributes |
| 10 | GSE171169 | Mouse | CD45-high immune cells | 2 mice/timepoint; 6,029 (5 d) and 4,896 (14 d) cells | scRNA-seq | Late infiltrating-leukocyte compartment | contributes |
| 11 | GSE227651 | Mouse | Whole brain | Sham/1/3/7 d; 11,744/17,780/11,983/17,922 cells | scRNA-seq | Early whole-brain context | contributes |
| 12 | GSE154396 | Mouse | CD45+ brain cells; Cd300a^fl/fl × LysM-Cre | 0 h/1 h/3 h after reperfusion; 51,005 cells | scRNA-seq | Genetic brake read | contributes |
| 13 | GSE332910 | Mouse | Striatum | Sham/0/24 h/7/14 d; n=3/group; 235,711 cells | snRNA-seq | Animal-level time course | contributes |
| 14 | GSE345807 | Rat | Whole brain | Sham n=2 / MCAO n=4; 72,018 cells | scRNA-seq | Chronic-window context | contributes |
| 15 | GSE267023 | Human | PBMC | 8 patients; 77,844 cells | scRNA-seq (GEX only) | Design-limited context | contributes |
| 16 | GSE337578 | Mouse | Brain, SeekSpace | 1 sample at day 3; 30,560 spots | Spatial transcriptomics | Spatial co-localization context | contributes |
| 17 | GSE318800 | Mouse | Brain myeloid; Ms4a3-Cre Htr2b cKO vs flox | n=2/group; 63,181 cells | scRNA-seq | Non-microglial genetic context | contributes |
| 18 | GSE137482 | Mouse | Parietal cortex | 3/18 months × sham/pMCAO d3 × n=6; 24 libraries | QuantSeq 3′ mRNA-seq | Age × injury factorial | contributes |
| 19 | GSE56267 | Human | Neocortex | 13 libraries; group of 3 undocumented | Bulk RNA-seq | Examined, not used (group assignment undocumented) | withdrawn or downgraded |
| 20 | GSE23651 | Rat | Cortex | Sham/vehicle/ZM447439 × 8/24 h; n=4/design cell; 24 samples | Microarray (GPL6101) | Acute rat context; exploratory | contributes |
| 21 | GSE37587 | Human | Whole blood | 34 paired patients (0–24 h vs 24–48 h; 68 samples) | Microarray (GPL6883) | Within-patient temporal context | contributes |
| 22 | GSE174574 | Mouse | Whole brain | Integrated with GSE319238/GSE227651; 110,664 cells | scRNA-seq | Cell-type ranking; d1 animal-level component | contributes |
| 23 | GSE319238 | Mouse | Whole brain | Integrated with GSE174574/GSE227651 | scRNA-seq | Cell-type ranking (integration constituent) | contributes |
| 24 | Rat 28 d (in-house) | Rat | Cortex, ischemic | Sham 7 / Model 3 / ICT 5 / ICTF 5 (n=20) | Bulk RNA-seq | Chronic brake/inactivation result | contributes |
| 25 | GSE137595 | Mouse | Cortex, ipsi/contra, ischaemic + sham | 24 samples (8/8/4/4); 2 h after 90 min MCAO/R | Microarray (GPL24242, Clariom S) | Earliest window; 2 × 2 ischaemia × hemisphere design, 24 arrays (§2.4) | contributes |
| 26 | GSE202518 | Human | Whole blood | 6 severe (NIHSS ≥ 8) / 6 moderate (NIHSS < 8) / 4 elderly controls; 16 libraries | Bulk RNA-seq (GPL24676, NovaSeq 6000) | Third whole-blood cohort; severity-stratified admission (§2.2); controls reanalysed from GSE192707 | contributes |
| 27 | GSE285659 | Human | PBMC, endovascular aspirate beyond the occluding MCA thrombus vs systemic artery | 3 paired patients; 78,809 cells (14,246 monocytes) | CITE-seq (GEX + 135 ADT) | Withdrawn: three paired donors cap the sign-flip probability at 0.25 | withdrawn or downgraded |
| 28 | GSE142361 | Mouse | FACS-sorted microglia | young/aged × sham/dMCAO, n = 2 per group; 8 libraries | Bulk RNA-seq (raw counts) | Withdrawn: the day-14 contrast was removed in v5 | withdrawn or downgraded |
| 29 | GSE225948 | Mouse | Brain CD45-intermediate/high leukocytes and CD45-negative endothelium | 13 brain samples (day 2: young sham 4, young stroke 4, aged sham 2, aged stroke 3) | scRNA-seq (10x) | Boundary: compartment inferable, interaction not estimable | contributes |
| 30 | GSE294912 | Mouse | Thromboembolic MCAo at 1 d | 8 animals, ipsi/contra paired (vehicle 4, BLZ945 4) | Bulk RNA-seq (CAM and whole tissue) | Not pooled: answers a different question | withdrawn or downgraded |
| 31 | GSE105132 | Mouse | Sorted microglia, macrophages and neutrophils at 48 h | MG 6 / MΦ 3 / NΦ 4, same ischaemic hemisphere, one batch; 13 arrays | Clariom array (oligo RMA) | Division of labour between cell types | contributes |
| 32 | GSE141972 | Mouse | Sorted microglia and macrophages at day 3 | WT MG 3 + WT MΦ 2; mKO 6; all post-MCAO, no sham | Bulk RNA-seq (featureCounts) | Context: usable but design-limited | contributes |
| 33 | GSE110141 | Mouse | MACS-sorted microglia at 24 h | WT/TG/KD × before/after MCAO, 2 vs 2 per arm; 12 arrays | Agilent array (two probe-set versions) | Knockdown contrast kept for one comparison only | withdrawn or downgraded |
| 34 | GSE248793 | Mouse | Sorted microglia | Sham/MCAO/±exosomes, n = 4 each; 16 libraries | RSEM FPKM | Downgraded to direction: sampling time undisclosed | withdrawn or downgraded |
| 35 | GSE64811 | Mouse | Sorted resident microglia (GFP+) and infiltrating macrophages (RFP+) at 7 d | 4 vs 4; both post-stroke, no sham; 8 arrays | Agilent GPL10787 | Withdrawn; BRIDGE layer retained | withdrawn or downgraded |
| 36 | GSE86941 | Rat | Microglia, monocytes and sham | day-3 family 9 + day-7 family 12; 21 samples, none excluded | RSEM transcripts | Constrained entry: journal-level FDR readout | contributes |
| 37 | GSE331114 | Human | Brain parenchyma, CD45-positive sorted | 4 stroke / 4 control donors; 8 libraries; 70,773 cells after QC | scRNA-seq (10x 3′ v3) | Primary: composition, against a 4-versus-4 resolution floor | contributes |
| 38 | GSE204702 | Human | Brain, 16 region combinations | 40 donors / 67 samples / 214,331 cells (9 disease classes, 1 control) | scRNA-seq (two CellRanger references) | Examined, not used: non-stroke, single control | withdrawn or downgraded |

GSE162955 is the microarray arm of a published multi-omics study of six paired patients (Ramiro et al., *Mol Neurobiol* 2021;58(8):4107-4121, doi:10.1007/s12035-021-02401-1). The companion label-free LC-MS/MS proteomics acquired on the same patients, on an LTQ Orbitrap Velos with gas-phase fractionation into four ranges, is deposited in the ProteomeXchange/PRIDE repository under PXD022850 (CC0). PXD022850 is not one of the 30 scored datasets above; it is recorded here as provenance for GSE162955.

Three further PRIDE series are used as a protein-level layer check and are likewise not among the thirty scored datasets: PXD012151 (rat cortex, 35 gel slices per sample, 1/7/14 days after 2 h tMCAO with reperfusion; 8,175 protein groups), PXD055588 (mouse microglia sorted 24 h after 1 h tMCAO, ipsilateral versus contralateral; 9,785 protein groups) and PXD025077 (mouse cortex after distal hypoxic MCAO, TMT 10-plex, 1/3/7/14/28 days; 8,068 quantified protein groups, of which 3,625 appear in the authors' changed-protein table). None is scored: layer coverage must reach 75% of a program's genes before it is called a score, and only PXD055588 clears that gate for CAPACITY (73/85 = 86%); PXD012151 reaches 42/85 (49%) and PXD025077 55/85 (64.7%), so both supply directional readouts only (Fig. 7, Table 15). PXD055588 deposits no run-to-group map, so its layer means are taken from the authors' own limma contrast rather than re-derived, and its differences are on the variance-stabilising scale rather than log2 fold change. PXD025077's TMT channel assignment was randomised within each plex and not deposited, so its contrasts are likewise read from the authors' per-protein table rather than re-derived; its five-point trajectory and the three checks that accompany it are set out in Note S1.11 and §2.10. The four remaining PRIDE series held in the project (PXD067754, PXD067636, PXD003218, PXD017280) are project-level only: none clears the gate, and each has a documented blocking issue (see repro/proteomics_manifest_20261024.csv). A sixth, PXD016538 (mouse neurovascular unit, 2 h after 90 min MCAO/R; 2,485 proteins, 2,466 gene symbols), is the label-free proteome paired with the GSE137595 transcriptomes; its CAPACITY coverage is 21/85 (24.7%), so the protein layer is not estimable and only the transcript arm is scored (see §2.11 and Note S1.10).

### Table S3. Program-level effect sizes

#### S3A. Human blood cohorts (case–control)

Scores are set means after within-sample z-scoring; p-values are two-sided Welch tests and q-values are BH-adjusted within cohort. GSE58294 statistics in this table are sample-level and are **not** used for inference in the main text, where the subject-level contrast is primary (23 vs 23; δ = +0.970, p = 5.5×10⁻¹¹, subject-label permutation p = 2.0×10⁻⁵).

The two-gene INACTIVATION composite is listed at the bottom of each blood cohort block because it is scored by a separate pipeline (human_efferocytosis_set_level.csv) and carries no Welch t statistic. Its q-value comes from a nine-test family, whereas the eight rows above use an eight-test family; re-running the families side by side (results/human_validation/gse_blood_set_level_9prog_bh.csv, columns q_bh_8prog_family, q_bh_9prog_family, q_9prog_minus_8prog) changes any q-value by at most 6.7×10⁻³ in absolute terms and leaves the set of programs significant in both cohorts unchanged (CAPACITY, OUTCOME, EXECUTOR). INACTIVATION is not scored for GSE22255 in this pipeline (status = not_scored_script95_cohort_list). It is reported here and not used for ranking for the same reason as in §2.7: with two genes the composite's dispersion is unstable, so its mean difference is not on a comparable scale with an 85-gene program.

| **Cohort** | **Program** | **Genes used** | **Δ score** | **Cliff's δ** | **p** | **q (BH)** | **n case/control** |
| --- | --- | --- | --- | --- | --- | --- | --- |
| GSE16561 | DEMAND | 167 | 0.216 | 0.327 | 3.34e-02 | 5.34e-02 | 39/24 |
| GSE16561 | PS_EXPOSURE | 6 | 0.325 | 0.259 | 9.74e-02 | 1.30e-01 | 39/24 |
| GSE16561 | FIND_ME | 7 | -0.170 | -0.233 | 1.65e-01 | 1.88e-01 | 39/24 |
| GSE16561 | CAPACITY | 84 | 0.374 | 0.483 | 1.58e-03 | 4.20e-03 | 39/24 |
| GSE16561 | BRIDGE | 6 | 0.199 | 0.216 | 3.02e-01 | 3.02e-01 | 39/24 |
| GSE16561 | BRAKE | 6 | 0.545 | 0.626 | 5.77e-06 | 2.72e-05 | 39/24 |
| GSE16561 | OUTCOME | 101 | 0.324 | 0.453 | 2.35e-03 | 4.71e-03 | 39/24 |
| GSE16561 | EXECUTOR | 23 | 0.272 | 0.592 | 6.80e-06 | 2.72e-05 | 39/24 |
| GSE16561 | INACTIVATION | 2 | 0.378 | 0.466 | 5.14e-02 | 7.71e-02 | 39/24 |
| GSE58294 | DEMAND | 178 | 0.334 | 0.637 | 2.86e-07 | 5.72e-07 | 69/23 |
| GSE58294 | PS_EXPOSURE | 5 | 0.287 | 0.338 | 2.67e-03 | 3.56e-03 | 69/23 |
| GSE58294 | FIND_ME | 7 | 0.150 | 0.275 | 8.20e-02 | 9.37e-02 | 69/23 |
| GSE58294 | CAPACITY | 85 | 0.613 | 0.928 | 4.61e-13 | 3.69e-12 | 69/23 |
| GSE58294 | BRIDGE | 6 | 0.400 | 0.468 | 2.42e-03 | 3.56e-03 | 69/23 |
| GSE58294 | BRAKE | 6 | 0.085 | 0.095 | 4.65e-01 | 4.65e-01 | 69/23 |
| GSE58294 | OUTCOME | 106 | 0.498 | 0.749 | 1.17e-07 | 3.12e-07 | 69/23 |
| GSE58294 | EXECUTOR | 24 | 0.508 | 0.710 | 5.13e-08 | 2.05e-07 | 69/23 |
| GSE58294 | INACTIVATION | 2 | 0.824 | 0.632 | 8.73e-07 | 1.57e-06 | 69/23 |
| GSE22255 | DEMAND | 178 | -0.022 | 0.105 | 7.19e-01 | 9.14e-01 | 20/20 |
| GSE22255 | PS_EXPOSURE | 5 | -0.005 | 0.025 | 9.69e-01 | 9.69e-01 | 20/20 |
| GSE22255 | FIND_ME | 7 | 0.032 | 0.000 | 8.00e-01 | 9.14e-01 | 20/20 |
| GSE22255 | CAPACITY | 85 | -0.054 | -0.085 | 5.82e-01 | 9.14e-01 | 20/20 |
| GSE22255 | BRIDGE | 6 | -0.128 | -0.090 | 4.71e-01 | 9.14e-01 | 20/20 |
| GSE22255 | BRAKE | 6 | -0.289 | -0.345 | 1.14e-01 | 4.58e-01 | 20/20 |
| GSE22255 | OUTCOME | 106 | -0.138 | -0.320 | 9.47e-02 | 4.58e-01 | 20/20 |
| GSE22255 | EXECUTOR | 24 | -0.060 | -0.045 | 5.26e-01 | 9.14e-01 | 20/20 |
| GSE22255 | INACTIVATION | — | — | — | — | — | 20/20 |

#### S3B. Rat 28-day cohort (treatment vs Model)

Group means are VST-based framework z-scores, one library per animal. δ is the Cliff's δ for treatment versus Model. The rescue index uses the signed definition given in Methods: (treatment mean − Model mean) / (Sham mean − Model mean), so +1 places the treated group on the sham mean and values outside 0–1 are possible when a group mean crosses either anchor (L1_PS_EXPOSURE under ICT is −2.053). Every column of this table is reproduced from results/rat_framework_28d/scale_model_vs_sham.csv (sham_mean, model_mean, ict_mean, ictf_mean, delta_raw_z, rescue_index_ict, rescue_index_ictf); Δ vs Model in the printed table is treatment mean minus Model mean, which is the numerator of the index with its sign intact. These are descriptive: Model n = 3, treatment n = 5, Sham n = 7, one bulk library per animal, and the cohort has no chronic vehicle arm.

| **Program** | **Treatment** | **Sham mean** | **Model mean** | **Treatment mean** | **Δ vs Model** | **Cliff's δ** | **Rescue index** |
| --- | --- | --- | --- | --- | --- | --- | --- |
| L1_APOPTOTIC_BURDEN | ICT | -0.3392 | 0.4639 | 0.0656 | -0.398 | -0.867 | 0.496 |
| L1_APOPTOTIC_BURDEN | ICTF | -0.3392 | 0.4639 | 0.1309 | -0.333 | -0.867 | 0.415 |
| L2_CAPACITY | ICT | -0.4231 | 0.5692 | 0.0725 | -0.497 | -1.000 | 0.501 |
| L2_CAPACITY | ICTF | -0.4231 | 0.5692 | 0.1783 | -0.391 | -0.733 | 0.394 |
| L2_LAP | ICT | 0.1239 | -0.1204 | -0.0843 | 0.036 | 0.200 | 0.148 |
| L2_LAP | ICTF | 0.1239 | -0.1204 | -0.0169 | 0.103 | 0.333 | 0.424 |
| L2_DEGRADATION | ICT | -0.9508 | 1.3449 | 0.2397 | -1.105 | -1.000 | 0.481 |
| L2_DEGRADATION | ICTF | -0.9508 | 1.3449 | 0.2845 | -1.060 | -0.867 | 0.462 |
| L2_DONT_EAT_ME | ICT | -0.7279 | 1.057 | 0.2119 | -0.845 | -1.000 | 0.473 |
| L2_DONT_EAT_ME | ICTF | -0.7279 | 1.057 | 0.173 | -0.884 | -1.000 | 0.495 |
| L2b_INACTIVATION | ICT | -0.7273 | 1.3307 | -0.1161 | -1.447 | -1.000 | 0.703 |
| L2b_INACTIVATION | ICTF | -0.7273 | 1.3307 | 0.3359 | -0.995 | -0.733 | 0.483 |
| L3_IMMUNOGENIC_DECOUPLED | ICT | -0.587 | 0.8086 | 0.1168 | -0.692 | -1.000 | 0.496 |
| L3_IMMUNOGENIC_DECOUPLED | ICTF | -0.587 | 0.8086 | 0.2198 | -0.589 | -0.867 | 0.422 |
| L4_STING_CARGO_OVERLOAD | ICT | -0.9021 | 1.2277 | 0.1865 | -1.041 | -1.000 | 0.489 |
| L4_STING_CARGO_OVERLOAD | ICTF | -0.9021 | 1.2277 | 0.3398 | -0.888 | -0.867 | 0.417 |
| L0_NECROPTOSIS_EXECUTOR | ICT | -0.4116 | 0.5565 | 0.0441 | -0.512 | -0.867 | 0.529 |
| L0_NECROPTOSIS_EXECUTOR | ICTF | -0.4116 | 0.5565 | 0.1982 | -0.358 | -0.733 | 0.370 |

### Table S4. Coverage of scored programs by dataset

Cells give detected/membership genes over the number the program contains. The scoring gate was 75% of a program's genes; a count marked ‡ falls below it, and that program was not scored in that dataset. “—” denotes a program not scored in that dataset or a coverage count not emitted to CSV. GSE22255 passed the gate in the scoring run, but its per-gene coverage counts were not re-emitted to a coverage CSV; the row is therefore left blank rather than reconstructed. GSE163654 was not scored with the frozen framework; it carries the cross-species rank-order comparison (Spearman ρ = +0.312 at 2 h and +0.371 at 6 h over 12,474 1:1 orthologs) and the gene-level sign-concordance test quoted in §2.2 (300 framework-core orthologs, script 103); GSE171169 and GSE227651 were scored for DEMAND, CAPACITY, OUTCOME and STING only, all four passing the gate, but their per-gene hit counts were not emitted to CSV.

| **Dataset** | **DEMAND** | **CAPACITY** | **BRAKE** | **INACT** | **OUTCOME** | **EXEC** | **STING** | **PS_EXP** |
| --- | --- | --- | --- | --- | --- | --- | --- | --- |
| GSE16561 | 167/178 | 84/85 | 6/6 | 2/2 | 101/108 | 23/24 | — | 6/6 |
| GSE58294 | 178/178 | 85/85 | 6/6 | 2/2 | 106/108 | 24/24 | — | 5/6 |
| GSE22255 | — | — | — | — | — | — | — | — |
| GSE23160 | 149/159 | 80/85 | 5/6 | 2/2 | 102/110 | 24/24 | — | 6/6 |
| P1/P2 | 159/159 | 81/85 | 6/6 | 2/2 | 110/110 | 24/24 | — | 6/6 |
| GSE250245 | 148/155 | 82/85 | 6/6 | 2/2 | 82/89 | 24/24 | — | 6/6 |
| Rat 28 d | 151/155 | 85/85 | 5/6 | 2/2 | 86/89 | 24/24 | 23/23 | 6/6 |
| GSE332910 | 158/159 | 82/85 | 6/6 | 2/2 | 107/110 | 24/24 | 39/41 | 6/6 |
| GSE345807 | 148/155 | 82/85 | 5/6 | 2/2 | 86/89 | 24/24 | 21/23 | 6/6 |
| GSE337578 | 159/159 | 82/85 | 6/6 | 2/2 | 107/110 | 24/24 | 39/41 | 6/6 |
| GSE267023 | 175/178 | 85/85 | 6/6 | 2/2 | 105/108 | 24/24 | 37/38 | 6/6 |
| GSE154396 | 158/159 | 82/85 | 6/6 | — | 106/110 | 24/24 | 38/41 | 6/6 |
| GSE318800 | 159/159 | 82/85 | 6/6 | 2/2 | 108/110 | 24/24 | 40/41 | 6/6 |
| GSE137482 | 159/159 | 81/85 | 6/6 | — | 110/110 | 24/24 | — | 6/6 |
| GSE56267 | 178/178 | 85/85 | 6/6 | — | 107/108 | 24/24 | — | 6/6 |
| GSE162955 | 172/178 | 81/85 | — | 2/2 | 101/108 | 23/24 | — | — |
| GSE23651 | 140/155 | 77/85 | 4/6‡ | 2/2 | 73/89 | 22/24 | 19/23 | 5/6 |
| GSE37587 | 167/178 | 84/85 | 6/6 | 2/2 | 101/108 | — | 36/38 | 6/6 |
| Integrated mouse | 157/159 | 82/85 | 6/6 | 2/2 | 93/110 | 24/24 | 26/41‡ | 6/6 |
| GSE137595 | 158/159 | 82/85 | 6/6 | 2/2 | 107/110 | 24/24 | 39/41 | 6/6 |

### Table S5. Operational phase classification

Phase II requires DEMAND > +0.05, STING > +0.05 and OUTCOME > +0.05; Phase I requires all three ≤ 0; all other combinations are Intermediate. The +0.05 margin is half the smallest between-group difference interpreted elsewhere and is not a validated cut-off. Only datasets with a complete DEMAND/STING/OUTCOME triad and a group baseline are classifiable: GSE23160, GSE137482, GSE250245 and the human blood cohorts lack STING; GSE154396, GSE318800, GSE267023 and GSE337578 lack a comparable baseline; GSE162955 is a paired core–contralateral contrast, not a timepoint series. These are omitted.

| **Dataset** | **Timepoint/group** | **DEMAND** | **STING** | **OUTCOME** | **Phase** |
| --- | --- | --- | --- | --- | --- |
| GSE171169 | 5 d | 0.1025 | 0.0213 | 0.0546 | Intermediate |
| GSE171169 | 14 d | 0.0933 | 0.1135 | 0.0766 | Phase II |
| GSE227651 | Sham | 0.1276 | 0.1116 | 0.0350 | Intermediate |
| GSE227651 | 1 d | 0.1020 | −0.0125 | 0.0022 | Intermediate |
| GSE227651 | 3 d | 0.1148 | 0.0601 | 0.0247 | Intermediate |
| GSE227651 | 7 d | 0.1104 | 0.0328 | 0.0143 | Intermediate |
| GSE332910 | Sham | −0.0620 | −0.0241 | −0.0647 | Phase I |
| GSE332910 | 0 h | 0.0741 | 0.0012 | 0.0284 | Intermediate |
| GSE332910 | 24 h | 0.0107 | 0.0059 | 0.0065 | Intermediate |
| GSE332910 | 7 d | −0.0076 | 0.0179 | 0.0127 | Intermediate |
| GSE332910 | 14 d | 0.0034 | 0.0006 | 0.0045 | Intermediate |
| GSE345807 | Sham | −0.0410 | −0.0731 | −0.0584 | Phase I |
| GSE345807 | MCAO | 0.0276 | 0.0491 | 0.0392 | Intermediate |
| Rat 28 d | Sham | −0.3392 | −0.9021 | −0.587 | Phase I |
| Rat 28 d | Model | 0.4639 | 1.2277 | 0.8086 | Phase II |
| Rat 28 d | ICT | 0.0656 | 0.1865 | 0.1168 | Phase II |
| Rat 28 d | ICTF | 0.1309 | 0.3398 | 0.2198 | Phase II |

The classification separates silent baseline from a cargo-loaded state in two datasets (GSE171169 at 14 d; the 28-day rat Model group, with ICT and ICTF also above the absolute margin). It does not produce a dated two-phase sequence in every dataset, and θ does not track the call: θ moves from +0.084 in the 28-day sham to −0.105 in the Phase II Model group, and is positive in the Phase I GSE332910 sham (+0.0118).

### Table S6. Composition adjustment of the CAPACITY effect

Each row is an OLS model of the CAPACITY score on stroke status plus the listed covariates. GSE16561 and GSE22255 models include age (z-scored) and sex; GSE58294 models do not, and the subject-mean rows use one value per donor (23 patients, 23 controls). Proxies are z-scored marker scores built after removing framework genes. Attenuation is the percentage reduction in the stroke coefficient relative to the model without proxies. Source: results/human_validation/composition_adjustment_audit.csv.

| **Cohort** | **Model** | **CAPACITY β** | **p** | **n** | **Attenuation %** |
| --- | --- | --- | --- | --- | --- |
| GSE58294 (subject means) | stroke | 0.613 | 5.49e-11 | 46 | — |
| GSE58294 (subject means) | stroke + NEUTROPHIL | 0.566 | 8.23e-09 | 46 | 7.7 |
| GSE58294 (subject means) | stroke + MONOCYTE | 0.404 | 2.62e-08 | 46 | 34.0 |
| GSE58294 (subject means) | stroke + LYMPHOID | 0.597 | 5.17e-11 | 46 | 2.6 |
| GSE58294 (subject means) | stroke + ERYTHROID | 0.601 | 4.21e-11 | 46 | 2.0 |
| GSE58294 (subject means) | stroke + PLATELET | 0.646 | 6.03e-14 | 46 | -5.4 |
| GSE58294 (subject means) | stroke + ALL5 | 0.473 | 2.59e-10 | 46 | 22.8 |
| GSE58294 (sample level) | stroke | 0.613 | 3.35e-13 | 92 | — |
| GSE58294 (sample level) | stroke + NEUTROPHIL | 0.539 | 3.74e-10 | 92 | 12.0 |
| GSE58294 (sample level) | stroke + MONOCYTE | 0.383 | 6.99e-10 | 92 | 37.5 |
| GSE58294 (sample level) | stroke + LYMPHOID | 0.600 | 2.96e-13 | 92 | 2.2 |
| GSE58294 (sample level) | stroke + ERYTHROID | 0.598 | 7.78e-14 | 92 | 2.4 |
| GSE58294 (sample level) | stroke + PLATELET | 0.645 | 4.56e-17 | 92 | -5.2 |
| GSE58294 (sample level) | stroke + ALL5 | 0.449 | 7.35e-13 | 92 | 26.7 |
| GSE16561 (whole blood) | stroke + age_z + male | 0.488 | 1.94e-04 | 63 | — |
| GSE16561 (whole blood) | stroke + age_z + male + NEUTROPHIL | 0.293 | 1.08e-02 | 63 | 39.9 |
| GSE16561 (whole blood) | stroke + age_z + male + MONOCYTE | 0.180 | 1.73e-01 | 63 | 63.2 |
| GSE16561 (whole blood) | stroke + age_z + male + LYMPHOID | 0.033 | 7.72e-01 | 63 | 93.2 |
| GSE16561 (whole blood) | stroke + age_z + male + ERYTHROID | 0.223 | 2.61e-03 | 63 | 54.2 |
| GSE16561 (whole blood) | stroke + age_z + male + PLATELET | 0.232 | 5.98e-03 | 63 | 52.5 |
| GSE16561 (whole blood) | stroke + age_z + male + ALL5 | 0.014 | 8.39e-01 | 63 | 97.1 |
| GSE22255 (PBMC) | stroke + age_z + male | -0.049 | 6.15e-01 | 40 | — |
| GSE22255 (PBMC) | stroke + age_z + male + NEUTROPHIL | -0.100 | 2.58e-01 | 40 | -106.1 |
| GSE22255 (PBMC) | stroke + age_z + male + MONOCYTE | -0.062 | 3.63e-01 | 40 | -27.3 |
| GSE22255 (PBMC) | stroke + age_z + male + LYMPHOID | -0.046 | 5.62e-01 | 40 | 4.8 |
| GSE22255 (PBMC) | stroke + age_z + male + ERYTHROID | -0.062 | 5.34e-01 | 40 | -27.9 |
| GSE22255 (PBMC) | stroke + age_z + male + PLATELET | -0.062 | 5.34e-01 | 40 | -26.5 |
| GSE22255 (PBMC) | stroke + age_z + male + ALL5 | -0.053 | 4.58e-01 | 40 | -9.9 |

### Table S7. Cell-type-specific program deltas in rat sMCAO (GSE250245)

Deltas are ipsilateral minus contralateral within the severe-MCAO libraries; intervals are cell-level stratified bootstrap intervals and split-half sign consistency is the proportion of 200 half-splits with the same sign. Six libraries, one animal per severity and hemisphere. The intervals describe within-library sampling variation and are not animal-level confidence intervals. Source: results/rat_gse250245/rat_bootstrap_delta.csv.

| **Cell type** | **Program** | **Nuclei (ipsi/contra)** | **Δ** | **95% CI** | **Split-half** |
| --- | --- | --- | --- | --- | --- |
| Microglia | DEMAND | 1608/1687 | 0.872 | 0.782 to 0.957 | 1.00 |
| Microglia | CAPACITY | 1608/1687 | 1.853 | 1.757 to 1.943 | 1.00 |
| Microglia | BRIDGE | 1608/1687 | 1.281 | 1.135 to 1.420 | 1.00 |
| Microglia | BRAKE | 1608/1687 | 0.787 | 0.694 to 0.885 | 1.00 |
| Microglia | OUTCOME | 1608/1687 | 1.321 | 1.224 to 1.416 | 1.00 |
| Microglia | EXECUTOR | 1608/1687 | 0.551 | 0.472 to 0.632 | 1.00 |
| Endothelial | DEMAND | 1813/2279 | 0.330 | 0.259 to 0.410 | 1.00 |
| Endothelial | CAPACITY | 1813/2279 | 0.760 | 0.681 to 0.837 | 1.00 |
| Endothelial | BRIDGE | 1813/2279 | 0.550 | 0.454 to 0.638 | 1.00 |
| Endothelial | BRAKE | 1813/2279 | 0.520 | 0.428 to 0.616 | 1.00 |
| Endothelial | OUTCOME | 1813/2279 | 0.490 | 0.418 to 0.562 | 1.00 |
| Endothelial | EXECUTOR | 1813/2279 | 0.134 | 0.066 to 0.204 | 1.00 |
| Pericyte | DEMAND | 1908/576 | 0.404 | 0.293 to 0.504 | 1.00 |
| Pericyte | CAPACITY | 1908/576 | 0.729 | 0.629 to 0.826 | 1.00 |
| Pericyte | BRIDGE | 1908/576 | 0.380 | 0.269 to 0.490 | 1.00 |
| Pericyte | BRAKE | 1908/576 | 0.177 | 0.088 to 0.268 | 1.00 |
| Pericyte | OUTCOME | 1908/576 | 0.588 | 0.489 to 0.688 | 1.00 |
| Pericyte | EXECUTOR | 1908/576 | 0.277 | 0.180 to 0.378 | 1.00 |
| Astrocyte | DEMAND | 2427/2539 | 0.423 | 0.362 to 0.486 | 1.00 |
| Astrocyte | CAPACITY | 2427/2539 | 0.592 | 0.531 to 0.651 | 1.00 |
| Astrocyte | BRIDGE | 2427/2539 | 0.315 | 0.254 to 0.377 | 1.00 |
| Astrocyte | BRAKE | 2427/2539 | 0.053 | -0.005 to 0.111 | 0.94 |
| Astrocyte | OUTCOME | 2427/2539 | 0.319 | 0.260 to 0.375 | 1.00 |
| Astrocyte | EXECUTOR | 2427/2539 | 0.129 | 0.072 to 0.188 | 1.00 |
| Oligodendrocyte | DEMAND | 3846/4091 | 0.457 | 0.409 to 0.505 | 1.00 |
| Oligodendrocyte | CAPACITY | 3846/4091 | 0.406 | 0.353 to 0.455 | 1.00 |
| Oligodendrocyte | BRIDGE | 3846/4091 | 0.234 | 0.185 to 0.279 | 1.00 |
| Oligodendrocyte | BRAKE | 3846/4091 | 0.178 | 0.131 to 0.220 | 1.00 |
| Oligodendrocyte | OUTCOME | 3846/4091 | 0.449 | 0.404 to 0.494 | 1.00 |
| Oligodendrocyte | EXECUTOR | 3846/4091 | 0.247 | 0.197 to 0.294 | 1.00 |
| Neuron | DEMAND | 10140/14133 | 0.150 | 0.128 to 0.171 | 1.00 |
| Neuron | CAPACITY | 10140/14133 | 0.087 | 0.067 to 0.107 | 1.00 |
| Neuron | BRIDGE | 10140/14133 | 0.019 | -0.002 to 0.039 | 0.99 |
| Neuron | BRAKE | 10140/14133 | 0.023 | 0.002 to 0.046 | 0.98 |
| Neuron | OUTCOME | 10140/14133 | 0.106 | 0.086 to 0.129 | 1.00 |
| Neuron | EXECUTOR | 10140/14133 | 0.091 | 0.068 to 0.114 | 1.00 |

### Table S8. Claims–evidence grade

| **Claim** | **Evidence in this paper** | **Grade** | **Main limit** |  |  |
| --- | --- | --- | --- | --- | --- |
| CAPACITY is the most consistent program in rat brain and human whole blood | Rat sMCAO +0.892 versus DEMAND +0.575; GSE58294 +0.613 (δ 0.970); GSE16561 +0.374 (δ 0.483) | Moderate–strong, within the eight programs | GSE16561 BRAKE effect is larger; "most consistent" is program-set-specific |  |  |
| The response is compartment- and composition-dependent | PBMC null (−0.054, q = 0.914); infarct core 59/81 genes down (p = 4.8×10⁻⁵); whole-blood attenuation after composition adjustment | Strong descriptive | Infarct-core necrosis and RNA degradation cannot be excluded; no causal test |  |  |
| θ adds little beyond DEMAND | θ-only R² 0.591 versus DEMAND 0.762 (striatum); ΔR² 0.057, p = 0.064 (striatum), 0.063, p = 0.108 (cortex); split-half CAPACITY r = 0.799 cortex / 0.445 striatum, striatum SB = 0.616 | Strong for the negative claim | n = 16; θ is descriptive, not a surrogate |  |  |
| Composition versus per-cell attribution | Oaxaca–Blinder: composition −5.0% to +21.0% across contrasts and bases; within-cell 79.0%–105.0%; severe-versus-moderate composition +13.7% to +21.0% | Moderate | Six libraries, one animal per group; decomposition assumptions |  |  |
| Microglia are the largest per-cell responder, but the weighted contribution is distributed | CAPACITY +1.853 (CI 1.757–1.943); +0.905 (0.751–1.070) from moderate to severe; 24.0% of the within-cell term versus neuron 24.6% | Moderate | snRNA one library per group; bootstrap precision is not animal-level |  |  |
| Chronic clearance, brake and inactivation programs co-induce | CAPACITY +0.992, BRAKE +1.785, INACTIVATION +2.058; θ remains −0.189; θ₂ +1.869 is definitional and exploratory | Descriptive | Model n = 3; BRAKE detected in 5/6 genes; θ₂ is post hoc and INACTIVATION is not net activity |  |  |
| A late immunogenic-output state co-moves with STING | GSE171169 CAPACITY 0.274→0.152; STING and OUTCOME rise | Weak–moderate | L4_STING overlaps OUTCOME (36/38, 39/41, 21/23); report as subscore |  |  |
| CD300a is a brake or coupling checkpoint | Literature plus expression/co-variation in this paper; GSE154396 shows compartment-specific deletion without an early capacity-program rise | Not tested here | No loss-of-function or occupancy experiment in this paper |  |  |
| ICT partially reverses the 28-day program | CAPACITY −0.497 (rescue index 0.501), BRAKE −0.845, INACTIVATION −1.447, θ₂ −1.348 | Weak descriptive | Model n = 3, ICT n = 5; one timepoint; no dose–response in this paper |  |  |
| At 2 h the clearance program is present but not engaged, while apoptotic demand has risen | GSE137595: CAPACITY +0.043 (p = 0.66, inside random-set null) versus DEMAND +0.239 (exceeds null, one-sided p = 0.002); θ +0.196 (q = 0.009, exceeds null); sham-arm negative control null; DiD indeterminate for all eight programs | Moderate–strong for the negative capacity claim; weak for the positive demand claim | 20 samples after animal-level QC exclusion; sham n = 3 per arm; the difference is between animals, not assignable to the ischaemic hemisphere; depth confound | ρ | = 0.747; no drug arm; 2 h not on the 1 d / 28 d axis |
| In whole tissue, the clearance program is not yet engaged at 24 h but is engaged in the following week | PXD025077 CAPACITY protein layer: day 1 +0.028 (inside size-matched null), day 3 +0.215, day 7 +0.789 (exceeds null, one-sided p = 0.0002), day 14 +0.495, day 28 +0.299; 17 of 32 members peak at day 7 (hypergeometric p = 7.5×10⁻⁵); authors' later-up cluster 10 versus 1.7 expected (p = 3.0×10⁻⁶) | Weak–moderate (directional readout only) | Coverage 55/85 (64.7%), below the 75% gate; contrasts read from a pre-filtered changed-protein table because the TMT channel map was not deposited; day-14 value fully explicable by infiltration; day-28 sham n = 2; no drug arm; not on the 2 h / 28 d axis |  |  |

### Table S9. Cross-cohort gene-level concordance and PBMC equivalence

| **Analysis** | **n** | **Statistic** | **Estimate** | **p** |
| --- | --- | --- | --- | --- |
| Shared genes, GSE16561 vs GSE58294 | 371 | Same-sign changes | 281 (75.7%; null 64.0%) | 1.0×10⁻⁴ (10,000 permutations; floor) |
| Shared genes, GSE16561 vs GSE58294 | 371 | Spearman ρ of log-fold-changes | +0.49 | 1.0×10⁻⁴ (10,000 permutations; floor) |
| CAPACITY subset | 83 | Same-sign changes | 68 (81.9%) | — |
| CAPACITY subset | 83 | Spearman ρ of log-fold-changes | +0.295 | 0.0068 |
| PBMC CAPACITY, GSE22255 | 20/20 | Mean difference (stroke − control) | −0.054 | 0.582 |
| PBMC CAPACITY, GSE22255 | 20/20 | 90% CI for difference | −0.218 to +0.110 | — |
| PBMC equivalence, margin 0.3 SD | 20/20 | Raw margin; TOST p; power | ±0.092; p = 0.348; power = 0.00 | not equivalent |

The 371-gene set is the intersection of genes detected in both whole-blood cohorts; the CAPACITY subset is the intersection with the 85-gene CAPACITY program. A 0.3-SD equivalence margin was not met: with a pooled SD of 0.307, the 90% CI spans zero and exceeds the margin, so the PBMC result is a null contrast, not a demonstrated equivalence.

### Table S10. CAPACITY expression-matched random-gene-set control

| **Cohort** | **Detected CAPACITY genes** | **Observed** | **Null mean** | **Null 95th percentile** | **One-sided p** | **Standardised distance** |
| --- | --- | --- | --- | --- | --- | --- |
| GSE16561 | 84 | +0.374 | +0.094 | +0.174 | 5.0e-04 | +5.60 |
| GSE58294 | 85 | +0.613 | +0.226 | +0.338 | 5.0e-04 | +5.48 |
| GSE22255 | 85 | -0.054 | -0.038 | +0.027 | 6.6e-01 | -0.43 |

For each cohort, 2,000 random gene sets of the same detected size were drawn from non-framework genes after expression-decile matching to the CAPACITY members. GSE58294 was averaged within subject before group comparison. The 2,000-draw resolution floor is p = 5.0×10⁻⁴. This control excludes gene number and expression-level artefacts but does not compare CAPACITY with a generic myeloid-activation signature.

### Table S11. Detection limits of the null and boundary contrasts

| **Dataset** | **Contrast** | **Program** | **n** | **Observed effect** | **Effect on SD scale** | **80%-power floor** | **Floor in score units** | **Reading** |
| --- | --- | --- | --- | --- | --- | --- | --- | --- |
| GSE56267 | stroke vs control | CAPACITY | 7/6 | 0.3445 | 1.2159 | 1.7127 | 0.4853 | not interpretable (group assignment undocumented) |
| GSE22255 | stroke vs control | CAPACITY | 20/20 | -0.0540 | -0.1757 | 0.9091 | 0.2793 | underpowered, and equivalence not established |
| GSE162955 | paired (core - contralateral) | theta | 6/6 | 0.1285 | 0.4110 | 1.4345 | paired dz scale; smallest attainable two-sided Wilcoxon p at n = 6 is 0.03125 | underpowered (observed paired d below MDE) and resolution-floored |
| GSE162955 | paired (core - contralateral) | theta2 | 6/6 | 0.3471 | 0.3608 | 1.4345 | paired dz scale; smallest attainable two-sided Wilcoxon p at n = 6 is 0.03125 | underpowered (observed paired d below MDE) and resolution-floored |
| GSE233815 | within-section rho(theta, OUTCOME) | theta | — | -0.441 to +0.083 | not_applicable | not_applicable | rho >= 0.9627 for 80% power; rho >= 0.8783 reaches alpha = 0.05 | cannot detect anything short of near-perfect correlation |

Source: results/negative_power/mde_negative_contrasts.csv (script 444). Floors are analytic at two-sided α = 0.05 and 80% power — noncentral t for the two-group and paired contrasts, Fisher z for the section-level correlation — computed from the already-shipped score files, not from resampling. The GSE162955 rows also carry a resolution limit: with six pairs the smallest attainable two-sided Wilcoxon p is 0.031. A cohort whose observed effect sits below its own floor cannot be read as evidence of absence, which is why §3.5 separates underpowered contrasts from the PBMC equivalence test, where a 0.3-SD margin was specified in advance and was not excluded either.

An earlier time-lag analysis (H2), computed by archived script 98b on the pre-freeze efferocytosis set and an ad-hoc injury panel in GSE23160, gave a striatum result of D = 1.217 (95% CI 0.411–6.434; bootstrap p = 0.0089; results/mouse_array/gse23160_h2_lag.json). Its gene sets and estimand are not the frozen L1/L2 hypotheses, so it is disclosed here for transparency and is not pooled with the framework contrasts.

### Table S12. Design floor of every layer × contrast in GSE86941

| **Contrast** | **Layer** | **k/det.** | **Δ** | **d** | **Floor** | **d/floor** | **q** | **Reading** |
| --- | --- | --- | --- | --- | --- | --- | --- | --- |
| 3d_macro_vs_sham | L2_CAPACITY | 85/76 | +0.173 | +2.16 | 3.071 | 0.70 | 0.2944 | uninf. null |
| 3d_macro_vs_sham | L1_APOPTOTIC_BURDEN | 155/133 | +0.037 | +0.66 | 3.071 | 0.22 | 0.5043 | uninf. null |
| 3d_macro_vs_sham | L3_IMMUNOGENIC_DECOUPLED | 89/77 | -0.118 | -1.60 | 3.071 | 0.52 | 0.9461 | uninf. null |
| 3d_macro_vs_sham | L2_DEGRADATION | 7/7 | +0.907 | +8.05 | 3.071 | 2.62 | 0.2550 | inf. null |
| 3d_macro_vs_sham | L3_IMMUNOGENIC_FULL | 560/480 | +0.302 | +4.52 | 3.071 | 1.47 | 0.0023 | sig |
| 3d_macro_vs_sham | L3_IMMUNOGENIC_EXT | 497/427 | +0.375 | +5.43 | 3.071 | 1.77 | 0.0023 | sig |
| 3d_macro_vs_sham | L3_IMMUNOGENIC_CORE | 114/100 | -0.295 | -3.97 | 3.071 | 1.29 | 0.4886 | inf. null |
| 3d_macro_vs_sham | L2_MACHINERY_FULL | 85/76 | +0.173 | +2.16 | 3.071 | 0.70 | 0.2944 | uninf. null |
| 3d_macro_vs_sham | L3_IMMUNOGENIC_CORE_STRICT | 49/44 | -0.165 | -1.76 | 3.071 | 0.57 | 0.9201 | uninf. null |
| 3d_macro_vs_sham | L4_PYROPTOSIS | 43/39 | +0.345 | +6.75 | 3.071 | 2.20 | 0.2426 | inf. null |
| 3d_macro_vs_sham | L2_PHAGO_MATUR | 33/28 | -0.209 | -2.16 | 3.071 | 0.70 | 0.9201 | uninf. null |
| 3d_macro_vs_sham | L0_NECROPTOSIS_EXECUTOR | 24/23 | +0.176 | +1.59 | 3.071 | 0.52 | 0.6010 | uninf. null |
| 3d_macro_vs_sham | L4_STING_CARGO_OVERLOAD | 23/20 | -0.252 | -1.60 | 3.071 | 0.52 | 0.9201 | uninf. null |
| 3d_macro_vs_sham | L2_ENGULF | 11/9 | -0.201 | -1.30 | 3.071 | 0.42 | 0.9201 | uninf. null |
| 3d_macro_vs_sham | L2_PS_RECEPTOR | 11/11 | +0.416 | +5.52 | 3.071 | 1.80 | 0.4886 | inf. null |
| 3d_macro_vs_sham | L2_LAP | 9/8 | +1.399 | +6.56 | 3.071 | 2.14 | 0.0613 | inf. null |
| 3d_macro_vs_sham | L1_FIND_ME | 7/7 | +0.006 | +0.03 | 3.071 | 0.01 | 0.9201 | uninf. null |
| 3d_macro_vs_sham | L1_PS_EXPOSURE | 6/4 | -0.374 | -1.35 | 3.071 | 0.44 | 0.9201 | uninf. null |
| 3d_macro_vs_sham | L2_BRIDGE | 6/6 | -1.454 | -3.40 | 3.071 | 1.11 | 0.1602 | inf. null |
| 3d_macro_vs_sham | L2_DONT_EAT_ME | 6/4 | -1.505 | -4.25 | 3.071 | 1.38 | — | not tested |
| 3d_macro_vs_sham | L2_METABOLIC | 5/5 | +1.583 | +6.73 | 3.071 | 2.19 | 0.1150 | inf. null |
| 3d_macro_vs_sham | L3_RESOLUTION_OUTPUT | 5/5 | +1.295 | +8.04 | 3.071 | 2.62 | 0.1853 | inf. null |
| 3d_macro_vs_sham | L2_TAM_RTK | 3/2 | -0.239 | -1.73 | 3.071 | 0.56 | — | not tested |
| 3d_macro_vs_sham | L2b_INACTIVATION | 2/1 | -1.369 | -3.36 | 3.071 | 1.09 | — | not tested |
| 3d_macro_vs_sham | EXT_CONTEXT_DEPENDENT | 1/1 | -2.259 | -13.69 | 3.071 | 4.46 | 0.1150 | inf. null |
| 3d_macro_vs_sham | L2_EAT_ME | 1/1 | +1.013 | +2.54 | 3.071 | 0.83 | 0.9201 | uninf. null |
| 3d_macro_vs_sham | HALLMARK_OXIDATIVE_PHOSPHORYLATION | 188/153 | +1.512 | +14.67 | 3.071 | 4.78 | 0.0006 | sig |
| 3d_macro_vs_sham | HALLMARK_GLYCOLYSIS | 197/160 | +0.413 | +5.41 | 3.071 | 1.76 | 0.0009 | sig |
| 3d_macro_vs_sham | HALLMARK_HYPOXIA | 197/162 | +0.200 | +6.25 | 3.071 | 2.03 | 0.0172 | sig |
| 3d_macro_vs_sham | OXPHOS_respiratory_only | 62/48 | +1.896 | +20.18 | 3.071 | 6.57 | 0.0002 | sig |
| 3d_macro_vs_sham | OXPHOS_minus_apop | 181/148 | +1.528 | +14.92 | 3.071 | 4.86 | 0.0002 | sig |
| 3d_macro_vs_sham | OXPHOS_minus_VATPase | 180/148 | +1.496 | +14.12 | 3.071 | 4.60 | 0.0002 | sig |
| 3d_microglia_vs_sham | L2_CAPACITY | 85/76 | +0.050 | +0.82 | 3.071 | 0.27 | 0.2734 | uninf. null |
| 3d_microglia_vs_sham | L1_APOPTOTIC_BURDEN | 155/133 | -0.114 | -0.99 | 3.071 | 0.32 | 0.7737 | uninf. null |
| 3d_microglia_vs_sham | L3_IMMUNOGENIC_DECOUPLED | 89/77 | -0.097 | -0.99 | 3.071 | 0.32 | 0.7737 | uninf. null |
| 3d_microglia_vs_sham | L2_DEGRADATION | 7/7 | +0.836 | +6.49 | 3.071 | 2.11 | 0.1368 | inf. null |
| 3d_microglia_vs_sham | L3_IMMUNOGENIC_FULL | 560/480 | +0.253 | +2.59 | 3.071 | 0.84 | 0.0023 | sig |
| 3d_microglia_vs_sham | L3_IMMUNOGENIC_EXT | 497/427 | +0.351 | +3.31 | 3.071 | 1.08 | 0.0023 | sig |
| 3d_microglia_vs_sham | L3_IMMUNOGENIC_CORE | 114/100 | -0.253 | -3.74 | 3.071 | 1.22 | 0.7737 | inf. null |
| 3d_microglia_vs_sham | L2_MACHINERY_FULL | 85/76 | +0.050 | +0.82 | 3.071 | 0.27 | 0.2734 | uninf. null |
| 3d_microglia_vs_sham | L3_IMMUNOGENIC_CORE_STRICT | 49/44 | -0.159 | -2.77 | 3.071 | 0.90 | 0.9971 | uninf. null |
| 3d_microglia_vs_sham | L4_PYROPTOSIS | 43/39 | +0.372 | +3.50 | 3.071 | 1.14 | 0.0368 | sig |
| 3d_microglia_vs_sham | L2_PHAGO_MATUR | 33/28 | -0.102 | -1.28 | 3.071 | 0.42 | 0.8503 | uninf. null |
| 3d_microglia_vs_sham | L0_NECROPTOSIS_EXECUTOR | 24/23 | +0.151 | +0.88 | 3.071 | 0.29 | 0.4429 | uninf. null |
| 3d_microglia_vs_sham | L4_STING_CARGO_OVERLOAD | 23/20 | -0.318 | -2.35 | 3.071 | 0.77 | 0.7737 | uninf. null |
| 3d_microglia_vs_sham | L2_ENGULF | 11/9 | -0.282 | -1.45 | 3.071 | 0.47 | 0.8503 | uninf. null |
| 3d_microglia_vs_sham | L2_PS_RECEPTOR | 11/11 | -0.169 | -0.91 | 3.071 | 0.30 | 0.9971 | uninf. null |
| 3d_microglia_vs_sham | L2_LAP | 9/8 | +0.260 | +1.12 | 3.071 | 0.36 | 0.6406 | uninf. null |
| 3d_microglia_vs_sham | L1_FIND_ME | 7/7 | +0.688 | +5.22 | 3.071 | 1.70 | 0.1978 | inf. null |
| 3d_microglia_vs_sham | L1_PS_EXPOSURE | 6/4 | +0.164 | +0.34 | 3.071 | 0.11 | 0.7737 | uninf. null |
| 3d_microglia_vs_sham | L2_BRIDGE | 6/6 | +0.145 | +3.18 | 3.071 | 1.04 | 0.7737 | inf. null |
| 3d_microglia_vs_sham | L2_DONT_EAT_ME | 6/4 | -0.991 | -2.80 | 3.071 | 0.91 | — | not tested |
| 3d_microglia_vs_sham | L2_METABOLIC | 5/5 | +0.199 | +1.44 | 3.071 | 0.47 | 0.7737 | uninf. null |
| 3d_microglia_vs_sham | L3_RESOLUTION_OUTPUT | 5/5 | +0.895 | +6.95 | 3.071 | 2.26 | 0.1858 | inf. null |
| 3d_microglia_vs_sham | L2_TAM_RTK | 3/2 | +0.616 | +9.81 | 3.071 | 3.19 | — | not tested |
| 3d_microglia_vs_sham | L2b_INACTIVATION | 2/1 | +0.032 | +0.03 | 3.071 | 0.01 | — | not tested |
| 3d_microglia_vs_sham | EXT_CONTEXT_DEPENDENT | 1/1 | -0.897 | -4.96 | 3.071 | 1.61 | 0.8054 | inf. null |
| 3d_microglia_vs_sham | L2_EAT_ME | 1/1 | +1.798 | +2.13 | 3.071 | 0.69 | 0.2734 | uninf. null |
| 3d_microglia_vs_sham | HALLMARK_OXIDATIVE_PHOSPHORYLATION | 188/153 | +0.801 | +4.15 | 3.071 | 1.35 | 0.0003 | sig |
| 3d_microglia_vs_sham | HALLMARK_GLYCOLYSIS | 197/160 | +0.213 | +1.82 | 3.071 | 0.59 | 0.0003 | sig |
| 3d_microglia_vs_sham | HALLMARK_HYPOXIA | 197/162 | +0.094 | +0.94 | 3.071 | 0.31 | 0.0036 | sig |
| 3d_microglia_vs_sham | OXPHOS_respiratory_only | 62/48 | +1.162 | +5.32 | 3.071 | 1.73 | 0.0002 | sig |
| 3d_microglia_vs_sham | OXPHOS_minus_apop | 181/148 | +0.835 | +4.41 | 3.071 | 1.44 | 0.0002 | sig |
| 3d_microglia_vs_sham | OXPHOS_minus_VATPase | 180/148 | +0.795 | +4.14 | 3.071 | 1.35 | 0.0002 | sig |
| 3d_macro_vs_microglia | L2_CAPACITY | 85/76 | +0.123 | +1.68 | 3.071 | 0.55 | 0.8696 | uninf. null |
| 3d_macro_vs_microglia | L1_APOPTOTIC_BURDEN | 155/133 | +0.150 | +1.46 | 3.071 | 0.48 | 0.8696 | uninf. null |
| 3d_macro_vs_microglia | L3_IMMUNOGENIC_DECOUPLED | 89/77 | -0.022 | -0.18 | 3.071 | 0.06 | 0.8696 | uninf. null |
| 3d_macro_vs_microglia | L2_DEGRADATION | 7/7 | +0.071 | +0.52 | 3.071 | 0.17 | 0.9817 | uninf. null |
| 3d_macro_vs_microglia | L3_IMMUNOGENIC_FULL | 560/480 | +0.049 | +0.42 | 3.071 | 0.14 | 0.9817 | uninf. null |
| 3d_macro_vs_microglia | L3_IMMUNOGENIC_EXT | 497/427 | +0.023 | +0.19 | 3.071 | 0.06 | 0.8696 | uninf. null |
| 3d_macro_vs_microglia | L3_IMMUNOGENIC_CORE | 114/100 | -0.042 | -0.46 | 3.071 | 0.15 | 0.8696 | uninf. null |
| 3d_macro_vs_microglia | L2_MACHINERY_FULL | 85/76 | +0.123 | +1.68 | 3.071 | 0.55 | 0.8696 | uninf. null |
| 3d_macro_vs_microglia | L3_IMMUNOGENIC_CORE_STRICT | 49/44 | -0.006 | -0.06 | 3.071 | 0.02 | 0.9542 | uninf. null |
| 3d_macro_vs_microglia | L4_PYROPTOSIS | 43/39 | -0.026 | -0.25 | 3.071 | 0.08 | 0.9107 | uninf. null |
| 3d_macro_vs_microglia | L2_PHAGO_MATUR | 33/28 | -0.107 | -0.93 | 3.071 | 0.30 | 0.8696 | uninf. null |
| 3d_macro_vs_microglia | L0_NECROPTOSIS_EXECUTOR | 24/23 | +0.026 | +0.14 | 3.071 | 0.05 | 0.9817 | uninf. null |
| 3d_macro_vs_microglia | L4_STING_CARGO_OVERLOAD | 23/20 | +0.066 | +0.34 | 3.071 | 0.11 | 0.9817 | uninf. null |
| 3d_macro_vs_microglia | L2_ENGULF | 11/9 | +0.082 | +0.50 | 3.071 | 0.16 | 0.9817 | uninf. null |
| 3d_macro_vs_microglia | L2_PS_RECEPTOR | 11/11 | +0.585 | +3.10 | 3.071 | 1.01 | 0.5694 | inf. null |
| 3d_macro_vs_microglia | L2_LAP | 9/8 | +1.138 | +5.39 | 3.071 | 1.76 | 0.0736 | inf. null |
| 3d_macro_vs_microglia | L1_FIND_ME | 7/7 | -0.682 | -2.64 | 3.071 | 0.86 | 0.5439 | uninf. null |
| 3d_macro_vs_microglia | L1_PS_EXPOSURE | 6/4 | -0.538 | -1.39 | 3.071 | 0.45 | 0.8696 | uninf. null |
| 3d_macro_vs_microglia | L2_BRIDGE | 6/6 | -1.600 | -3.73 | 3.071 | 1.22 | 0.0138 | sig |
| 3d_macro_vs_microglia | L2_DONT_EAT_ME | 6/4 | -0.514 | -2.38 | 3.071 | 0.77 | — | not tested |
| 3d_macro_vs_microglia | L2_METABOLIC | 5/5 | +1.385 | +5.64 | 3.071 | 1.84 | 0.0736 | inf. null |
| 3d_macro_vs_microglia | L3_RESOLUTION_OUTPUT | 5/5 | +0.400 | +3.18 | 3.071 | 1.04 | 0.8696 | inf. null |
| 3d_macro_vs_microglia | L2_TAM_RTK | 3/2 | -0.855 | -6.85 | 3.071 | 2.23 | — | not tested |
| 3d_macro_vs_microglia | L2b_INACTIVATION | 2/1 | -1.401 | -1.46 | 3.071 | 0.48 | — | not tested |
| 3d_macro_vs_microglia | EXT_CONTEXT_DEPENDENT | 1/1 | -1.362 | -5.66 | 3.071 | 1.84 | 0.8696 | inf. null |
| 3d_macro_vs_microglia | L2_EAT_ME | 1/1 | -0.785 | -0.94 | 3.071 | 0.31 | 0.8696 | uninf. null |
| 3d_macro_vs_microglia | HALLMARK_OXIDATIVE_PHOSPHORYLATION | 188/153 | +0.710 | +3.40 | 3.071 | 1.11 | 0.0006 | sig |
| 3d_macro_vs_microglia | HALLMARK_GLYCOLYSIS | 197/160 | +0.200 | +1.84 | 3.071 | 0.60 | 0.1650 | uninf. null |
| 3d_macro_vs_microglia | HALLMARK_HYPOXIA | 197/162 | +0.106 | +1.08 | 3.071 | 0.35 | 0.5685 | uninf. null |
| 3d_macro_vs_microglia | OXPHOS_respiratory_only | 62/48 | +0.734 | +3.30 | 3.071 | 1.07 | 0.0002 | sig |
| 3d_macro_vs_microglia | OXPHOS_minus_apop | 181/148 | +0.694 | +3.39 | 3.071 | 1.10 | 0.0002 | sig |
| 3d_macro_vs_microglia | OXPHOS_minus_VATPase | 180/148 | +0.701 | +3.34 | 3.071 | 1.09 | 0.0002 | sig |
| 7d_macro_vs_sham | L2_CAPACITY | 85/76 | +0.340 | +2.37 | 2.381 | 0.99 | 0.0015 | sig |
| 7d_macro_vs_sham | L1_APOPTOTIC_BURDEN | 155/135 | -0.219 | -1.70 | 2.381 | 0.71 | 0.7758 | uninf. null |
| 7d_macro_vs_sham | L3_IMMUNOGENIC_DECOUPLED | 89/77 | -0.192 | -0.98 | 2.381 | 0.41 | 0.7720 | uninf. null |
| 7d_macro_vs_sham | L2_DEGRADATION | 7/7 | +1.180 | +6.48 | 2.381 | 2.72 | 0.0314 | sig |
| 7d_macro_vs_sham | L3_IMMUNOGENIC_FULL | 560/483 | +0.183 | +1.03 | 2.381 | 0.43 | 0.0015 | sig |
| 7d_macro_vs_sham | L3_IMMUNOGENIC_EXT | 497/430 | +0.267 | +1.43 | 2.381 | 0.60 | 0.0015 | sig |
| 7d_macro_vs_sham | L3_IMMUNOGENIC_CORE | 114/100 | -0.413 | -2.32 | 2.381 | 0.98 | 0.5594 | uninf. null |
| 7d_macro_vs_sham | L2_MACHINERY_FULL | 85/76 | +0.340 | +2.37 | 2.381 | 0.99 | 0.0034 | sig |
| 7d_macro_vs_sham | L3_IMMUNOGENIC_CORE_STRICT | 49/44 | -0.442 | -2.18 | 2.381 | 0.92 | 0.6648 | uninf. null |
| 7d_macro_vs_sham | L4_PYROPTOSIS | 43/38 | +0.357 | +3.01 | 2.381 | 1.26 | 0.0314 | sig |
| 7d_macro_vs_sham | L2_PHAGO_MATUR | 33/28 | -0.156 | -0.69 | 2.381 | 0.29 | 0.7758 | uninf. null |
| 7d_macro_vs_sham | L0_NECROPTOSIS_EXECUTOR | 24/23 | -0.074 | -0.32 | 2.381 | 0.14 | 0.7016 | uninf. null |
| 7d_macro_vs_sham | L4_STING_CARGO_OVERLOAD | 23/20 | -0.362 | -1.03 | 2.381 | 0.43 | 0.8273 | uninf. null |
| 7d_macro_vs_sham | L2_ENGULF | 11/9 | +0.056 | +0.14 | 2.381 | 0.06 | 0.7016 | uninf. null |
| 7d_macro_vs_sham | L2_PS_RECEPTOR | 11/11 | +0.539 | +2.55 | 2.381 | 1.07 | 0.1355 | inf. null |
| 7d_macro_vs_sham | L2_LAP | 9/8 | +0.697 | +3.42 | 2.381 | 1.44 | 0.1355 | inf. null |
| 7d_macro_vs_sham | L1_FIND_ME | 7/7 | -0.062 | -0.26 | 2.381 | 0.11 | 0.7758 | uninf. null |
| 7d_macro_vs_sham | L1_PS_EXPOSURE | 6/5 | -0.408 | -1.64 | 2.381 | 0.69 | 0.8611 | uninf. null |
| 7d_macro_vs_sham | L2_BRIDGE | 6/6 | +0.761 | +3.50 | 2.381 | 1.47 | 0.1564 | inf. null |
| 7d_macro_vs_sham | L2_DONT_EAT_ME | 6/4 | -1.358 | -3.32 | 2.381 | 1.40 | — | not tested |
| 7d_macro_vs_sham | L2_METABOLIC | 5/5 | +1.108 | +7.23 | 2.381 | 3.04 | 0.0949 | inf. null |
| 7d_macro_vs_sham | L3_RESOLUTION_OUTPUT | 5/5 | +1.411 | +4.95 | 2.381 | 2.08 | 0.0368 | sig |
| 7d_macro_vs_sham | L2_TAM_RTK | 3/2 | -0.071 | -0.35 | 2.381 | 0.15 | — | not tested |
| 7d_macro_vs_sham | L2b_INACTIVATION | 2/1 | -2.050 | -3.52 | 2.381 | 1.48 | — | not tested |
| 7d_macro_vs_sham | EXT_CONTEXT_DEPENDENT | 1/1 | -2.186 | -6.17 | 2.381 | 2.59 | 0.1150 | inf. null |
| 7d_macro_vs_sham | L2_EAT_ME | 1/1 | +1.302 | +1.37 | 2.381 | 0.57 | 0.6648 | uninf. null |
| 7d_macro_vs_sham | HALLMARK_OXIDATIVE_PHOSPHORYLATION | 188/152 | +1.320 | +9.39 | 2.381 | 3.94 | 0.0006 | sig |
| 7d_macro_vs_sham | HALLMARK_GLYCOLYSIS | 197/164 | +0.097 | +1.27 | 2.381 | 0.54 | 0.0015 | sig |
| 7d_macro_vs_sham | HALLMARK_HYPOXIA | 197/163 | -0.100 | -0.69 | 2.381 | 0.29 | 0.1308 | uninf. null |
| 7d_macro_vs_sham | OXPHOS_respiratory_only | 62/47 | +1.811 | +11.26 | 2.381 | 4.73 | 0.0002 | sig |
| 7d_macro_vs_sham | OXPHOS_minus_apop | 181/147 | +1.343 | +9.12 | 2.381 | 3.83 | 0.0002 | sig |
| 7d_macro_vs_sham | OXPHOS_minus_VATPase | 180/147 | +1.304 | +9.18 | 2.381 | 3.85 | 0.0002 | sig |
| 7d_microglia_vs_sham | L2_CAPACITY | 85/76 | +0.147 | +1.02 | 2.381 | 0.43 | 0.0583 | uninf. null |
| 7d_microglia_vs_sham | L1_APOPTOTIC_BURDEN | 155/135 | -0.092 | -0.61 | 2.381 | 0.25 | 0.6970 | uninf. null |
| 7d_microglia_vs_sham | L3_IMMUNOGENIC_DECOUPLED | 89/77 | -0.127 | -0.63 | 2.381 | 0.26 | 0.8380 | uninf. null |
| 7d_microglia_vs_sham | L2_DEGRADATION | 7/7 | +1.207 | +5.82 | 2.381 | 2.44 | 0.0184 | sig |
| 7d_microglia_vs_sham | L3_IMMUNOGENIC_FULL | 560/483 | +0.026 | +0.16 | 2.381 | 0.07 | 0.0023 | sig |
| 7d_microglia_vs_sham | L3_IMMUNOGENIC_EXT | 497/430 | +0.058 | +0.36 | 2.381 | 0.15 | 0.0023 | sig |
| 7d_microglia_vs_sham | L3_IMMUNOGENIC_CORE | 114/100 | -0.244 | -1.30 | 2.381 | 0.55 | 0.8380 | uninf. null |
| 7d_microglia_vs_sham | L2_MACHINERY_FULL | 85/76 | +0.147 | +1.02 | 2.381 | 0.43 | 0.0583 | uninf. null |
| 7d_microglia_vs_sham | L3_IMMUNOGENIC_CORE_STRICT | 49/44 | -0.294 | -1.34 | 2.381 | 0.56 | 0.8380 | uninf. null |
| 7d_microglia_vs_sham | L4_PYROPTOSIS | 43/38 | +0.314 | +1.94 | 2.381 | 0.81 | 0.0583 | uninf. null |
| 7d_microglia_vs_sham | L2_PHAGO_MATUR | 33/28 | -0.169 | -0.79 | 2.381 | 0.33 | 0.9101 | uninf. null |
| 7d_microglia_vs_sham | L0_NECROPTOSIS_EXECUTOR | 24/23 | +0.081 | +0.33 | 2.381 | 0.14 | 0.6311 | uninf. null |
| 7d_microglia_vs_sham | L4_STING_CARGO_OVERLOAD | 23/20 | -0.355 | -1.12 | 2.381 | 0.47 | 0.8380 | uninf. null |
| 7d_microglia_vs_sham | L2_ENGULF | 11/9 | -0.037 | -0.09 | 2.381 | 0.04 | 0.8380 | uninf. null |
| 7d_microglia_vs_sham | L2_PS_RECEPTOR | 11/11 | -0.066 | -0.38 | 2.381 | 0.16 | 0.8380 | uninf. null |
| 7d_microglia_vs_sham | L2_LAP | 9/8 | +0.306 | +1.40 | 2.381 | 0.59 | 0.6311 | uninf. null |
| 7d_microglia_vs_sham | L1_FIND_ME | 7/7 | -0.102 | -0.39 | 2.381 | 0.16 | 0.8582 | uninf. null |
| 7d_microglia_vs_sham | L1_PS_EXPOSURE | 6/5 | -0.361 | -1.76 | 2.381 | 0.74 | 0.8542 | uninf. null |
| 7d_microglia_vs_sham | L2_BRIDGE | 6/6 | +0.714 | +4.89 | 2.381 | 2.06 | 0.2162 | inf. null |
| 7d_microglia_vs_sham | L2_DONT_EAT_ME | 6/4 | -1.268 | -3.70 | 2.381 | 1.55 | — | not tested |
| 7d_microglia_vs_sham | L2_METABOLIC | 5/5 | +0.271 | +1.89 | 2.381 | 0.79 | 0.7381 | uninf. null |
| 7d_microglia_vs_sham | L3_RESOLUTION_OUTPUT | 5/5 | +0.866 | +2.81 | 2.381 | 1.18 | 0.1840 | inf. null |
| 7d_microglia_vs_sham | L2_TAM_RTK | 3/2 | +0.205 | +1.00 | 2.381 | 0.42 | — | not tested |
| 7d_microglia_vs_sham | L2b_INACTIVATION | 2/1 | -0.992 | -1.57 | 2.381 | 0.66 | — | not tested |
| 7d_microglia_vs_sham | EXT_CONTEXT_DEPENDENT | 1/1 | -1.243 | -3.73 | 2.381 | 1.56 | 0.8380 | inf. null |
| 7d_microglia_vs_sham | L2_EAT_ME | 1/1 | +1.091 | +1.96 | 2.381 | 0.82 | 0.7381 | uninf. null |
| 7d_microglia_vs_sham | HALLMARK_OXIDATIVE_PHOSPHORYLATION | 188/152 | +1.198 | +7.28 | 2.381 | 3.06 | 0.0006 | sig |
| 7d_microglia_vs_sham | HALLMARK_GLYCOLYSIS | 197/164 | -0.031 | -0.63 | 2.381 | 0.26 | 0.1359 | uninf. null |
| 7d_microglia_vs_sham | HALLMARK_HYPOXIA | 197/163 | -0.196 | -1.55 | 2.381 | 0.65 | 0.9887 | uninf. null |
| 7d_microglia_vs_sham | OXPHOS_respiratory_only | 62/47 | +1.545 | +8.29 | 2.381 | 3.48 | 0.0002 | sig |
| 7d_microglia_vs_sham | OXPHOS_minus_apop | 181/147 | +1.224 | +7.24 | 2.381 | 3.04 | 0.0002 | sig |
| 7d_microglia_vs_sham | OXPHOS_minus_VATPase | 180/147 | +1.175 | +7.18 | 2.381 | 3.02 | 0.0002 | sig |
| 7d_macro_vs_microglia | L2_CAPACITY | 85/76 | +0.193 | +2.30 | 2.381 | 0.97 | 0.0276 | sig |
| 7d_macro_vs_microglia | L1_APOPTOTIC_BURDEN | 155/135 | -0.126 | -1.23 | 2.381 | 0.52 | 0.8008 | uninf. null |
| 7d_macro_vs_microglia | L3_IMMUNOGENIC_DECOUPLED | 89/77 | -0.065 | -0.40 | 2.381 | 0.17 | 0.9153 | uninf. null |
| 7d_macro_vs_microglia | L2_DEGRADATION | 7/7 | -0.027 | -0.14 | 2.381 | 0.06 | 0.9153 | uninf. null |
| 7d_macro_vs_microglia | L3_IMMUNOGENIC_FULL | 560/483 | +0.157 | +0.83 | 2.381 | 0.35 | 0.0023 | sig |
| 7d_macro_vs_microglia | L3_IMMUNOGENIC_EXT | 497/430 | +0.209 | +1.01 | 2.381 | 0.42 | 0.0023 | sig |
| 7d_macro_vs_microglia | L3_IMMUNOGENIC_CORE | 114/100 | -0.170 | -1.25 | 2.381 | 0.52 | 0.6292 | uninf. null |
| 7d_macro_vs_microglia | L2_MACHINERY_FULL | 85/76 | +0.193 | +2.30 | 2.381 | 0.97 | 0.0276 | sig |
| 7d_macro_vs_microglia | L3_IMMUNOGENIC_CORE_STRICT | 49/44 | -0.148 | -0.78 | 2.381 | 0.33 | 0.8008 | uninf. null |
| 7d_macro_vs_microglia | L4_PYROPTOSIS | 43/38 | +0.042 | +0.27 | 2.381 | 0.12 | 0.7861 | uninf. null |
| 7d_macro_vs_microglia | L2_PHAGO_MATUR | 33/28 | +0.013 | +0.12 | 2.381 | 0.05 | 0.8008 | uninf. null |
| 7d_macro_vs_microglia | L0_NECROPTOSIS_EXECUTOR | 24/23 | -0.155 | -1.75 | 2.381 | 0.73 | 0.8008 | uninf. null |
| 7d_macro_vs_microglia | L4_STING_CARGO_OVERLOAD | 23/20 | -0.007 | -0.02 | 2.381 | 0.01 | 0.8008 | uninf. null |
| 7d_macro_vs_microglia | L2_ENGULF | 11/9 | +0.093 | +0.46 | 2.381 | 0.19 | 0.8008 | uninf. null |
| 7d_macro_vs_microglia | L2_PS_RECEPTOR | 11/11 | +0.605 | +4.10 | 2.381 | 1.72 | 0.0276 | sig |
| 7d_macro_vs_microglia | L2_LAP | 9/8 | +0.390 | +1.79 | 2.381 | 0.75 | 0.3461 | uninf. null |
| 7d_macro_vs_microglia | L1_FIND_ME | 7/7 | +0.039 | +0.26 | 2.381 | 0.11 | 0.8008 | uninf. null |
| 7d_macro_vs_microglia | L1_PS_EXPOSURE | 6/5 | -0.046 | -0.16 | 2.381 | 0.07 | 0.9153 | uninf. null |
| 7d_macro_vs_microglia | L2_BRIDGE | 6/6 | +0.047 | +0.23 | 2.381 | 0.10 | 0.8008 | uninf. null |
| 7d_macro_vs_microglia | L2_DONT_EAT_ME | 6/4 | -0.090 | -0.32 | 2.381 | 0.13 | — | not tested |
| 7d_macro_vs_microglia | L2_METABOLIC | 5/5 | +0.837 | +4.68 | 2.381 | 1.97 | 0.0759 | inf. null |
| 7d_macro_vs_microglia | L3_RESOLUTION_OUTPUT | 5/5 | +0.545 | +3.11 | 2.381 | 1.31 | 0.3272 | inf. null |
| 7d_macro_vs_microglia | L2_TAM_RTK | 3/2 | -0.276 | -2.46 | 2.381 | 1.03 | — | not tested |
| 7d_macro_vs_microglia | L2b_INACTIVATION | 2/1 | -1.058 | -3.00 | 2.381 | 1.26 | — | not tested |
| 7d_macro_vs_microglia | EXT_CONTEXT_DEPENDENT | 1/1 | -0.943 | -1.99 | 2.381 | 0.83 | 0.6292 | uninf. null |
| 7d_macro_vs_microglia | L2_EAT_ME | 1/1 | +0.211 | +0.20 | 2.381 | 0.08 | 0.8008 | uninf. null |
| 7d_macro_vs_microglia | HALLMARK_OXIDATIVE_PHOSPHORYLATION | 188/152 | +0.122 | +0.94 | 2.381 | 0.40 | 0.0042 | sig |
| 7d_macro_vs_microglia | HALLMARK_GLYCOLYSIS | 197/164 | +0.128 | +2.09 | 2.381 | 0.88 | 0.0042 | sig |
| 7d_macro_vs_microglia | HALLMARK_HYPOXIA | 197/163 | +0.096 | +0.77 | 2.381 | 0.32 | 0.0078 | sig |
| 7d_macro_vs_microglia | OXPHOS_respiratory_only | 62/47 | +0.266 | +2.57 | 2.381 | 1.08 | 0.0036 | sig |
| 7d_macro_vs_microglia | OXPHOS_minus_apop | 181/147 | +0.118 | +0.90 | 2.381 | 0.38 | 0.0034 | sig |
| 7d_macro_vs_microglia | OXPHOS_minus_VATPase | 180/147 | +0.129 | +0.97 | 2.381 | 0.41 | 0.0024 | sig |

All rows are GSE86941. Each row is one layer × contrast. Framework layers, the three Hallmark anchors (names beginning HALLMARK_) and the three size-matched oxidative variants (names beginning OXPHOS_) each form their own Benjamini–Hochberg family. k/det. is the number of set members after one-to-one mouse→rat mapping and the number entering the score in that family. Floor is the minimum detectable standardised difference at α = 0.05 two-sided and 80% power, solved exactly from the noncentral t distribution at the design's n (3 versus 3 at day 3; 4 versus 4 at day 7); d/floor ≥ 1 means the observed difference lies at or above that floor. q is Benjamini–Hochberg within the layer's own family and tests specificity against size-matched random sets on the same samples; it is not sample-level significance. Reading: sig (q < 0.05); inf. null (q ≥ 0.05 and |d| at or above the floor); uninf. null (q ≥ 0.05 and |d| below the floor); not tested (the layer fails the 75% coverage gate in this dataset — L2_DONT_EAT_ME with 4 of 6 members, L2_TAM_RTK 2 of 3, L2b_INACTIVATION 1 of 2 — so it never entered the specificity family, carries no q, and supports no negative claim; its floor is listed for completeness only). Rows are not independent: framework layers overlap by construction, and the three size-matched variants are nested within the oxidative anchor. Three entries with one or two genes — EXT_CONTEXT_DEPENDENT, L2_EAT_ME and L2b_INACTIVATION — are markers, not programs; their d values are not program-level effect sizes and are listed only for completeness. Source: results/newdata/gse86941_measurement_envelope.csv; scripts 593 and 597.

### Table S13. Point-by-point contrast with prior reports

| **Prior finding (citation)** | **What it claims** | **What this study finds** | **Relation** |
| --- | --- | --- | --- |
| A bulk or single-cell-derived efferocytosis score characterises ischaemic stroke and its hub genes are proposed as markers [9] | Efferocytosis patterns computed from bulk RNA with single-cell support, with Abca1, C1qc, Ptx3, Irf5 and Pros1 nominated as hub genes | Reproduced in direction, bounded in interpretation. In whole blood the same kind of program score does rise after stroke, but most of that rise is attributable to lineage composition (Table 6; 52–105% of the gap on the pooled basis in GSE16561), so a blood score cannot be read as a per-cell program. | Bounded |
| CD300a blockade enhances efferocytosis by infiltrating myeloid cells and improves neurological deficit after stroke [32] | A causal, protein-level claim: blocking an inhibitory receptor increases uptake and improves outcome | Not contradicted and not tested. CD300A sits in the brake layer and is scored separately from CAPACITY; its transcript did not move in any of the four protein-layer contrasts, and knockout of Cd300a was detectable but small (δ = −0.26 pooled). A transcript readout cannot test receptor blockade; at 28 days the brake raw score rises more than CAPACITY, which is consistent with a brake-dominant chronic phase rather than with blockade. | Not tested at the level of the claim |
| Reviews nominate STAT6, PPARG, CD300A and SIGMAR1 as candidate efferocytosis modulators in stroke [7,8] | Candidate list assembled from the literature, without a shared measurement scale | Only CD300A is scored here, and only as part of the six-gene brake. The other three are not in any layer, so this paper neither supports nor excludes them; they are outside its measurement frame rather than tested and rejected. | Out of scope |
| Soluble TREM2 and recombinant MFG-E8 improve outcome in stroke models [41,43]; plasma sTREM2 tracks death and cardiovascular events in patients [42] | Protein-level and shed-form claims | Not measurable here. TREM2 and MFG8 are in the gene sets, but neither the shed form nor the ligand–integrin engagement can be read from mRNA, and no plasma protein was measured in this study. Listed as an explicit gap, not as a negative result. | Out of scope |
| Clearance capacity is saturable and depends on phagocyte metabolic state, including Ucp2 [62,63] | Capacity as a saturable, rate-limited quantity | A transcriptional score cannot address saturation: it measures how much machinery is transcribed, not how much substrate a cell can process. This is the reason the paper reports layer deltas and explicitly declines to call them clearance rates. | Bound |

Relation vocabulary: *reproduced* — the same direction is observed on a comparable readout; *bounded* — the direction holds but the inference the prior work draws is not supported at this level of measurement; *not tested* — the claim is causal or protein-level and no measurement here can adjudicate it; *out of scope* — the entity was never measured. No row is a refutation: the papers above and this one measure different things, and the table exists so that the difference is on the record rather than left to the reader.

### Table S14. Probe-aggregation sensitivity of the CAPACITY effect

| **Cohort** | **Aggregation** | **CAPACITY genes** | **Effect** | **Δ vs published** | **Baseline β** | **Adjusted β** | **Attenuation** |
| --- | --- | --- | --- | --- | --- | --- | --- |
| GSE58294 | mean of probes (published) | 85 | +0.613 | +0.000 | +0.613 | +0.473 | 22.8% |
| GSE58294 | sum of probes | 85 | +0.613 | +0.000 | +0.613 | +0.473 | 22.8% |
| GSE58294 | median of probes | 85 | +0.507 | −0.105 | +0.507 | +0.369 | 27.3% |
| GSE58294 | single brightest probe | 85 | +0.605 | −0.008 | +0.605 | +0.449 | 25.8% |
| GSE16561 | mean of probes (published) | 84 | +0.374 | +0.000 | +0.488 | +0.014 | 97.1% |
| GSE16561 | sum of probes | 84 | +0.376 | +0.001 | +0.488 | +0.016 | 96.7% |
| GSE16561 | median of probes | 84 | +0.361 | −0.013 | +0.475 | +0.005 | 99.0% |
| GSE16561 | single brightest probe | 84 | +0.374 | −0.001 | +0.467 | +0.016 | 96.6% |

Baseline model: stroke only in GSE58294, whose 92 GEO records carry no age field; stroke plus age and sex in GSE16561. The adjusted model adds the five lineage proxies. Sum and mean are algebraically identical here — summing multiplies each gene by its probe count and the score standardises every gene before averaging — and the identity is asserted in script 696 rather than assumed; in GSE16561 the two differ by 0.0015 because 64 CAPACITY genes have a partially missing probe vector, and a plain sum propagates the missing value where a mean does not. Across the three admissible aggregations the CAPACITY effect stays positive and the attenuation stays in 22.8–27.3% for GSE58294 and 96.6–99.0% for GSE16561. The choice of aggregation moves the 22.8% point estimate by at most 4.5 percentage points and never changes a conclusion.

### Table S15. GO-only reconstruction of the vote-built output program

| **Cohort** | **Set** | **Genes in set** | **Detected** | **Effect** | **Welch p** | **Cliff's δ** |
| --- | --- | --- | --- | --- | --- | --- |
| GSE58294 | GO_ONLY_DECOUPLED | 17 | 17 | +0.753 | 6.708e−08 | +0.875 |
| GSE58294 | PUBLISHED_OUTCOME | 106 | 106 | +0.498 | 2.035e−07 | +0.815 |
| GSE16561 | GO_ONLY_DECOUPLED | 17 | 17 | +0.451 | 0.0002515 | +0.536 |
| GSE16561 | PUBLISHED_OUTCOME | 101 | 101 | +0.324 | 0.002353 | +0.453 |

GO-only reconstruction of a **vote-built** set. L2_CAPACITY is curated by hand from named sublayers and two hard-coded receptor lists (Table 2; results/io_framework/framework_layer_source_map.csv). It is not the product of a vote, so a GO-only CAPACITY does not exist and the reviewer's request cannot be executed as literally phrased. The vote-built program in this framework is the immunogenic core (L3_IMMUNOGENIC_CORE, majority vote over KEGG, MSigDB and GO sources). Rebuilding it from the three GO biological-process sources alone (GO_IL1B_PROD, GO_TLR_SIGNALING, GO_TYPE_I_IFN_PROD) and applying the published decoupling rule leaves a much smaller set, so this is a direction check, not a like-for-like comparison of magnitudes: with 17 genes the GO-only set moves in the same direction and at a comparable or larger standardised effect than the published set in both cohorts. Library-based construction therefore does not manufacture the immunogenic-output signal. Source: repro/go_only_reconstruction_20261030.csv; script 696.

### Table S16. Detection-matched null for the CAPACITY excess

| **Cohort** | **Matching scheme** | **Observed** | **Null mean** | **Null SD** | **SD relative to the published null** | **Null 95th** | **One-sided p** | **Standardised distance** | **Calibrated** |
| --- | --- | --- | --- | --- | --- | --- | --- | --- | --- |
| GSE16561 | decile_only | +0.374 | +0.093 | 0.052 | 1.00 | +0.175 | 5.0×10⁻4 | +5.45 | Y |
| GSE16561 | decile_depth25 | +0.374 | +0.082 | 0.050 | 0.97 | +0.165 | 5.0×10⁻4 | +5.80 | N |
| GSE16561 | decile_depth50 | +0.374 | +0.105 | 0.048 | 0.93 | +0.182 | 5.0×10⁻4 | +5.57 | N |
| GSE16561 | decile_depth25_sd | +0.374 | +0.117 | 0.051 | 0.98 | +0.201 | 5.0×10⁻4 | +5.07 | Y |
| GSE16561 | decile_setdepth25 | +0.374 | +0.074 | 0.055 | 1.06 | +0.163 | 5.0×10⁻4 | +5.48 | Y |
| GSE16561 | decile_setdepth50 | +0.374 | +0.087 | 0.051 | 0.99 | +0.172 | 5.0×10⁻4 | +5.60 | Y |
| GSE58294 | decile_only | +0.613 | +0.225 | 0.069 | 1.00 | +0.340 | 5.0×10⁻4 | +5.62 | Y |
| GSE58294 | decile_depth25 | +0.613 | +0.134 | 0.028 | 0.40 | +0.177 | 5.0×10⁻4 | +17.39 | N |
| GSE58294 | decile_depth50 | +0.613 | +0.113 | 0.042 | 0.61 | +0.182 | 5.0×10⁻4 | +11.89 | N |
| GSE58294 | decile_depth25_sd | +0.613 | +0.185 | 0.068 | 0.98 | +0.299 | 5.0×10⁻4 | +6.33 | Y |
| GSE58294 | decile_setdepth25 | +0.613 | +0.228 | 0.070 | 1.02 | +0.343 | 5.0×10⁻4 | +5.47 | Y |
| GSE58294 | decile_setdepth50 | +0.613 | +0.225 | 0.069 | 1.00 | +0.340 | 5.0×10⁻4 | +5.63 | Y |
| GSE22255 | decile_only | −0.054 | −0.036 | 0.039 | 1.00 | +0.028 | 0.67 | −0.46 | Y |
| GSE22255 | decile_depth25 | −0.054 | −0.095 | 0.013 | 0.33 | −0.075 | 5.0×10⁻4 | +3.21 | N |
| GSE22255 | decile_depth50 | −0.054 | −0.114 | 0.024 | 0.62 | −0.075 | 5.0×10⁻3 | +2.51 | N |
| GSE22255 | decile_depth25_sd | −0.054 | −0.048 | 0.036 | 0.92 | +0.011 | 0.57 | −0.17 | Y |
| GSE22255 | decile_setdepth25 | −0.054 | −0.036 | 0.038 | 0.96 | +0.026 | 0.67 | −0.47 | Y |
| GSE22255 | decile_setdepth50 | −0.054 | −0.034 | 0.039 | 1.00 | +0.032 | 0.70 | −0.51 | Y |

All schemes exclude framework genes and match the detected CAPACITY members on expression decile; the depth-matched schemes additionally match detection depth, defined as the fraction of samples in which a gene rises above that sample's own 25th or 50th percentile of gene-level expression. Matching is either gene by gene (each CAPACITY member replaced by a background gene from the same decile drawn from its ten nearest neighbours in depth) or set by set (whole sets drawn as in the published control and kept only if their mean depth is within a half-SD caliper of the observed mean). B = 2,000 draws per cohort and scheme; p is one-sided at the 5.0×10⁻⁴ resolution floor. **Reading.** The last column is the point of the table: it is read off GSE22255, the cohort that does not replicate the CAPACITY increase. A correctly calibrated null leaves that cohort non-significant. The gene-by-gene depth schemes fail that check — they shrink the null SD to 0.33–0.62 of the published null and return p at the floor in a cohort whose point estimate is negative — so their p values are an artefact of an over-constrained null and are reported here only to document why they are not used. Both set-level schemes pass the check (p = 0.67–0.70 in GSE22255) and still return p at the floor in the two replication cohorts. Detection depth therefore does not explain the CAPACITY excess, and CAPACITY's own depth profile is not a fixed property: it sits below the background in GSE16561 (0.711 versus 0.742) and above it in GSE58294 (0.896 versus 0.742). Source: results/human_validation/specificity_controls/detection_matched_null_summary.csv; script 695.

### Table S17. Energy-axis scope: every dataset, its status and the reason

This is the screening registry for the energy axis, and it is wider than the 38 assembled datasets of section 4.2: it also keeps the accessions that were examined and set aside before assembly, and it folds GSE319238 into the GSE174574 row because the two form one integrated analysis. Every row is listed one by one with the reason it did not enter, so that a reader can apply a different rule rather than inherit this one; nine of the accessions below never reached the assembled inventory and appear here only so that the screening is on the record. This registry lists contrasts, not datasets: thirty datasets were assessed for eligibility and eighteen contrasts in six of them met both conditions. Two conditions were necessary — an explicit between-group contrast, and at least four libraries in each group — and a third was scope: the axis was computed only where it tests a claim the paper already makes. The eight rows marked † meet both necessary conditions and were still not run; that is a judgement about what the paper needed, not a limit of the data, and each of them could be scored on the same engine without any further curation.

| **#** | **Dataset** | **Status** | **Basis** | **Reason** |
| --- | --- | --- | --- | --- |
| 1 | GSE86941 | Scored | — | Scored on the unified engine; its arms are in Table 11 |
| 2 | GSE202518 | Scored | — | Scored on the unified engine; its arms are in Table 11 |
| 3 | GSE331114 | Scored | — | Scored on the unified engine; its arms are in Table 11 |
| 4 | GSE16561 | Scored | — | Scored on the unified engine; its arms are in Table 11 |
| 5 | GSE58294 | Scored | — | Scored on the unified engine; its arms are in Table 11 |
| 6 | GSE22255 | Scored | — | Scored on the unified engine; its arms are in Table 11 |
| 7 | GSE142361 | Withdrawn | — | Withdrawn for library depth (section 2.8); its aged and young arms share one row z-standardisation, so no separable subset exists |
| 8 | GSE23160† | Not run | scope | Four arrays per group and a 2/8/24 h design; it would support an energy axis, but the paper makes no energy claim about this series |
| 9 | GSE163654† | Not run | scope | Twelve arrays across mouse and rat cortex; conditions are met, no energy claim |
| 10 | GSE23651† | Not run | scope | 24 arrays; conditions are met, no energy claim |
| 11 | GSE137482† | Not run | scope | 24 libraries, six per cell, ageing design; conditions are met, and the ageing claim in section 2.13 does not involve the energy axis |
| 12 | GSE294912† | Not run | scope | 16 + 16 libraries; conditions are met, no energy claim |
| 13 | GSE105132† | Not run | scope | 13 libraries across sorted microglia, macrophages and neutrophils; conditions are met, no energy claim |
| 14 | GSE37587† | Not run | scope | 68 whole-blood arrays with a time dimension; conditions are met, but the series is used as a time-replication check and not for energy |
| 15 | GSE148350† | Not run | scope | 30 sorted-microglia arrays; conditions are met, and the series is archived as a reversed boundary result |
| 16 | GSE250245 | Not run | structure | Single-nucleus series with one animal per severity and hemisphere; no animal-level replicate, so a per-group permutation p cannot be formed |
| 17 | GSE233815 | Not run | structure | Visium sections without a between-group contrast; spots mix cell types and there is no group factor to test |
| 18 | GSE337578 | Not run | structure | One SeekSpace section; no between-group contrast |
| 19 | GSE232936 | Not run | structure | Four aggregated PSD matrices; the archived condition contrast cannot be rebuilt |
| 20 | GSE171169 | Not run | structure | ScRNA-seq, CD45-high cells at two time points; one animal per time point, so cell-level bootstrap cannot stand in for an animal-level test |
| 21 | GSE227651 | Not run | structure | ScRNA-seq; one animal per time point |
| 22 | GSE332910 | Not run | structure | SnRNA-seq with three animals per condition but 235,711 cells dominated by neuronal mitochondrial transcripts; energy anchors are not separable from cell composition at nucleus level |
| 23 | GSE318800 | Not run | structure | Four libraries; no group structure usable for a two-group contrast |
| 24 | GSE154396 | Not run | structure | Six libraries across a genotype series; no untreated-versus-treated contrast with four or more per arm |
| 25 | GSE345807 | Not run | structure | Six libraries; no between-group contrast with four per arm |
| 26 | GSE267023 | Not run | structure | Eight patients, within-patient enrichment only; no case-control contrast |
| 27 | GSE304010 | Not run | structure | Four FACS-enriched libraries; no replicate structure |
| 28 | GSE174574 | Not run | structure | Integration reference; no contrast of its own |
| 29 | GSE285659 | Not run | structure | Three paired donors; the sign-flip bound at n = 3 is 0.25, so no exact test can reach significance |
| 30 | GSE225948 | Not run | structure | Aged arm has two sham and three injured animals; below the four-per-group floor |
| 31 | GSE137595 | Not run | structure | The 2 h mouse series has seven paired hemispheres in the ischaemic arm but three sham animals, below the four-per-group floor |
| 32 | GSE64811 | Not run | structure | Four libraries per arm and a paired design that no non-parametric test can resolve at n = 4 (section 2.8) |
| 33 | GSE248793 | Not run | structure | Four groups of four; the dataset is already archived as directional only |
| 34 | GSE141972 | Not run | structure | Wild-type 5 versus knockout 6 libraries; the contrast is genotype, not injury |
| 35 | GSE110141 | Not run | structure | Two platforms mixed within one series and six per arm; the platform switch is confounded with the genotype arm |
| 36 | P1-P2 | Not run | structure | In-house mouse series with 19 samples across four injury models; no single two-group contrast of interest for the energy axis |
| 37 | Rat28d | Not run | structure | The 28-day cohort retains three model animals after exclusions |
| 38 | GSE162955 | Not run | — | 29 layer scores, none reproducible; it is reported as a boundary in section 2.3 and carries no capacity claim, so an energy axis there would inherit the same failure |
| 39 | GSE56267 | Not run | — | The group of three of its thirteen libraries is undocumented and the case–control estimate moves between +0.135 and +0.453 depending on where they are placed, so no contrast from it is used (section 2.3) |
| 40 | GSE131193 | Not run | not scored | Excluded before scoring for tissue comparability |
| 41 | GSE212008 | Not run | not scored | CRISPR screen, not a transcriptome |
| 42 | GSE175504 | Not run | not scored | Nhe1 cKO series in an old build; no capacity readout |
| 43 | GSE171171 | Not run | not scored | Wrong accession number; withdrawn from the inventory |
| 44 | GSE110993 | Not run | not scored | MiRNA only; no mRNA endpoint |
| 45 | GSE326055 | Not run | not scored | Two pooled libraries; archived for compartment description only |
| 46 | GSE204702 | Not run | not scored | Not a stroke cohort; no age, sex or post-mortem interval for controls |

**Reading.** Of the forty-six rows, six were scored on the unified engine, one is a series withdrawn elsewhere in this paper, and thirty-nine were not run. Within those thirty-nine: twenty-two have no between-group contrast, or no contrast with four libraries per arm, so the expression-matched floor for a set of this size sits above any difference the data can produce; two are withdrawn for reasons already given in sections 2.3 and 2.8; seven were never scored for reasons unrelated to the energy axis; and eight meet both necessary conditions but carry no energy claim in this paper. The last group is the one a reader should press on, and it is the reason the rule is written out rather than left implicit: the axis asks a question this paper only asks in six places, and extending it elsewhere would add tabulated numbers without adding a test of anything stated here. Source: repro/bioenergetic_unified_exclusions.csv; scripts 732 (ledger) and 733 (this table).

### Table S18. Correction families: what each Benjamini–Hochberg step was applied over

Every q value in the paper is a within-family quantity. The family is the set of programs scored on the same samples in the same design, and no family is pooled across designs, compartments or species — which is why a q is never compared with a q from another family. Table A lists each testing family that carries a q in the main text, read from the analysis ledger rather than typed. Tables B and C split the two axes whose families are defined by block rather than by dataset, because there a single dataset contributes more than one correction. “Scored” is the number of program scores that entered the family; “with q” is the number that produced a q, which is smaller wherever a program was dropped for coverage or returned a degenerate null.

**A. Family per dataset and contrast.**

| **Dataset** | **Contrast** | **Scored** | **With q** | **Smallest q in family** | **Any q < 0.05** |
| --- | --- | --- | --- | --- | --- |
| GSE110141_KD | KD_after_minus_before | 3 | 3 | 0.0007 | yes |
| GSE110141_genotype | TG_minus_WT_after_P10787 | 2 | 2 | 0.0005 | yes |
| GSE110141_genotype | TG_minus_WT_before_P21163 | 1 | 1 | 0.8362 | no |
| GSE137482 | aged_vs_young_within_mcao | 7 | 7 | 0.0006 | yes |
| GSE141972_mKO | TAK1mKO_vs_WT_macrophage_3d | 3 | 3 | 0.0202 | yes |
| GSE141972_mKO | TAK1mKO_vs_WT_microglia_3d | 2 | 2 | 0.0221 | yes |
| GSE148350 | 12mon_14d_L | 1 | 1 | 0.5192 | no |
| GSE148350 | 12mon_3d_L | 1 | 1 | 0.3536 | no |
| GSE148350 | 3mon_14d_L | 2 | 2 | 0.0004 | yes |
| GSE148350 | 3mon_3d_L | 2 | 2 | 0.0004 | yes |
| GSE16561 | Stroke_vs_Ctrl | 5 | 3 | 0.0003 | yes |
| GSE202518 | MS_vs_Ctrl | 3 | 3 | 0.0003 | yes |
| GSE202518 | SS_vs_MS | 4 | 4 | 0.0006 | yes |
| GSE202518 | Stroke_vs_Ctrl | 5 | 3 | 0.0004 | yes |
| GSE22255 | IS_vs_Ctrl | 2 | 2 | 0.0006 | yes |
| GSE225948 | aged_vs_young_within_mcao[EC] | 5 | 5 | 0.0033 | yes |
| GSE225948 | aged_vs_young_within_mcao[MdC] | 5 | 5 | 0.0033 | yes |
| GSE225948 | aged_vs_young_within_mcao[Mg] | 5 | 5 | 0.0033 | yes |
| GSE248793 | MCAO_vs_Sham | 5 | 5 | 0.0442 | yes |
| GSE331114 | AIS_vs_Control | 20 | 16 | 0.0001 | yes |
| GSE331114 | GRANULOCYTE；AIS_vs_Control | 1 | 0 | — | no |
| GSE331114 | MYELOID_INFILTRATE；AIS_vs_Control | 1 | 0 | — | no |
| GSE331114 | MYELOID_RESIDENT；AIS_vs_Control | 2 | 0 | — | no |
| GSE58294 | Stroke_vs_Ctrl | 7 | 3 | 0.0003 | yes |
| GSE64811 | paired_MP_minus_MG_7d | 3 | 3 | 0.0156 | yes |
| GSE86941 | 3d_macro_vs_microglia | 2 | 2 | 0.0138 | yes |
| GSE86941 | 3d_macro_vs_sham | 6 | 6 | 0.0006 | yes |
| GSE86941 | 7d_macro_vs_microglia | 2 | 2 | 0.0276 | yes |
| GSE86941 | 7d_macro_vs_sham | 3 | 3 | 0.0006 | yes |
| GSE86941 | 7d_microglia_vs_sham | 3 | 3 | 0.0006 | yes |

Source: ledger_claims_20261101.csv — 113 scores in 30 testing families. The two in-house series carry no q of their own and are not listed. A family of eight is the eight counted programs; the nine-test families of the blood cohorts are the eight plus INACTIVATION, which is reported with its own family size where it is used because it is scored by a separate pipeline.

**B. Energy axis, family by block and arm class.**

| **Dataset** | **Block** | **Arm class** | **Family size** | **Arms scored** | **Arms over floor** |
| --- | --- | --- | --- | --- | --- |
| GSE16561 | GSE16561 | demand | 2 | 2 | 1 |
| GSE16561 | GSE16561 | demand_projection | 3 | 3 | 3 |
| GSE16561 | GSE16561 | reference | 3 | 3 | 3 |
| GSE16561 | GSE16561 | supply | 3 | 3 | 1 |
| GSE202518 | GSE202518 | demand | 2 | 8 | 2 |
| GSE202518 | GSE202518 | demand_projection | 3 | 12 | 0 |
| GSE202518 | GSE202518 | reference | 3 | 12 | 3 |
| GSE202518 | GSE202518 | supply | 3 | 12 | 2 |
| GSE22255 | GSE22255 | demand | 2 | 2 | 1 |
| GSE22255 | GSE22255 | demand_projection | 3 | 3 | 0 |
| GSE22255 | GSE22255 | reference | 3 | 3 | 2 |
| GSE22255 | GSE22255 | supply | 3 | 3 | 2 |
| GSE331114_GRANULOCYTE | GSE331114_GRANULOCYTE | demand_projection | 2 | 2 | 0 |
| GSE331114_GRANULOCYTE | GSE331114_GRANULOCYTE | reference | 1 | 1 | 0 |
| GSE331114_GRANULOCYTE | GSE331114_GRANULOCYTE | supply | 1 | 1 | 0 |
| GSE331114_MYELOID_INFILTRATE | GSE331114_MYELOID_INFILTRATE | demand | 2 | 2 | 0 |
| GSE331114_MYELOID_INFILTRATE | GSE331114_MYELOID_INFILTRATE | demand_projection | 3 | 3 | 0 |
| GSE331114_MYELOID_INFILTRATE | GSE331114_MYELOID_INFILTRATE | reference | 3 | 3 | 1 |
| GSE331114_MYELOID_INFILTRATE | GSE331114_MYELOID_INFILTRATE | supply | 3 | 3 | 1 |
| GSE331114_MYELOID_RESIDENT | GSE331114_MYELOID_RESIDENT | demand | 1 | 1 | 0 |
| GSE331114_MYELOID_RESIDENT | GSE331114_MYELOID_RESIDENT | demand_projection | 3 | 3 | 0 |
| GSE331114_MYELOID_RESIDENT | GSE331114_MYELOID_RESIDENT | reference | 3 | 3 | 0 |
| GSE331114_MYELOID_RESIDENT | GSE331114_MYELOID_RESIDENT | supply | 3 | 3 | 0 |
| GSE331114_NK | GSE331114_NK | demand | 1 | 1 | 0 |
| GSE331114_NK | GSE331114_NK | demand_projection | 2 | 2 | 0 |
| GSE331114_NK | GSE331114_NK | reference | 3 | 3 | 0 |
| GSE331114_NK | GSE331114_NK | supply | 1 | 1 | 0 |
| GSE331114_T_CELL | GSE331114_T_CELL | demand | 1 | 1 | 0 |
| GSE331114_T_CELL | GSE331114_T_CELL | demand_projection | 3 | 3 | 0 |
| GSE331114_T_CELL | GSE331114_T_CELL | reference | 3 | 3 | 1 |
| GSE331114_T_CELL | GSE331114_T_CELL | supply | 1 | 1 | 1 |
| GSE58294 | GSE58294 | demand | 2 | 2 | 0 |
| GSE58294 | GSE58294 | demand_projection | 3 | 3 | 1 |
| GSE58294 | GSE58294 | reference | 3 | 3 | 1 |
| GSE58294 | GSE58294 | supply | 3 | 3 | 1 |
| GSE86941 | GSE86941_3d | demand | 3 | 9 | 4 |
| GSE86941 | GSE86941_3d | demand_projection | 2 | 6 | 2 |
| GSE86941 | GSE86941_3d | reference | 3 | 9 | 0 |
| GSE86941 | GSE86941_3d | supply | 3 | 9 | 4 |
| GSE86941 | GSE86941_7d | demand | 3 | 9 | 2 |
| GSE86941 | GSE86941_7d | demand_projection | 2 | 6 | 0 |
| GSE86941 | GSE86941_7d | reference | 3 | 9 | 1 |
| GSE86941 | GSE86941_7d | supply | 3 | 9 | 2 |

Source: results/newdata/bioenergetic_unified.csv. Supply and demand form separate three-arm families and are never pooled. The reference and projection arms repeat their block's family size for completeness but are not hypotheses of their own, which is why Table 11 does not mark them.

**C. GSE331114, family by compartment and arm class.**

| **Compartment** | **Arm class** | **Family size** | **Layers scored** | **Layers with q < 0.05** |
| --- | --- | --- | --- | --- |
| GRANULOCYTE | bioenergetic_anchor | 1 | 1 | 0 |
| GRANULOCYTE | framework | 4 | 4 | 0 |
| MYELOID_INFILTRATE | bioenergetic_anchor | 3 | 3 | 1 |
| MYELOID_INFILTRATE | framework | 19 | 19 | 1 |
| MYELOID_RESIDENT | bioenergetic_anchor | 3 | 3 | 0 |
| MYELOID_RESIDENT | framework | 18 | 18 | 0 |
| NK | bioenergetic_anchor | 1 | 1 | 0 |
| NK | framework | 13 | 13 | 0 |
| T_CELL | bioenergetic_anchor | 1 | 1 | 1 |
| T_CELL | framework | 16 | 16 | 0 |

Source: results/newdata/gse331114_two_gate.csv. Framework programs and energy anchors are kept apart, so a compartment contributes its total number of scored layers to the paper but never a single correction over all of them. In the T-cell compartment the energy family has one member, so its q equals its gene-set p: the correction there is empty, not strong.

**D. Sensitivity: what one correction over all energy arms would leave.**

| **Scope** | **Arms tested** | **Arms with q < 0.05** | **Both gates** | **Smallest q through both gates** |
| --- | --- | --- | --- | --- |
| Within family, as reported | 183 | 88 | 38 | 0.0002 |
| One correction over all arms | 183 | 82 | 37 | 0.0009149 |

Source: bioenergetic_unified.csv, restricted to the 183 arms that produced a p value. 18 families were scored; family sizes run from 4 to 11, with 14 of the 18 families at the maximum and a median of 11 — small enough that a reader may ask whether within-family correction is too permissive. The second row answers that directly: a single Benjamini–Hochberg step over every arm on the unified engine leaves 37 arms through both gates against 38 under the reported within-family correction, a difference of one arm. The only arm that does not survive the stricter scope is GSE22255 IS_vs_Ctrl HALLMARK_HYPOXIA (+0.101, 1.84x floor, q 0.0400 to 0.0703). No arm passes the global scope and fails the within-family one, as a monotone correction over the same p values requires. The last column is not a ranking of strength. Both values sit at or near the resolution limit of a 10,000-draw null, where the smallest attainable two-sided p is 2/(10,001) and a q can go no lower; several arms reach it, so equal values do not mean equally strong effects. It is reported only to show that the within-family and global scopes do not coincide. The global scope is not used anywhere in the paper: it pools arms across tissues, species and designs, so it corrects for hypotheses that were never tested together. It is reported here as a lower bound on how much of the energy result survives the strictest available reading.

**E. Two nulls, two questions: gene-set permutation against sample-label permutation.**

| **Dataset** | **Samples** | **Smallest attainable label p** | **Arms scored** | **Gene-set p < 0.05** | **Label p ≤ 0.05** | **Label p at its floor** | **Conflict** |
| --- | --- | --- | --- | --- | --- | --- | --- |
| GSE16561 | 39 vs 24 | 9.999e-05 (10,000 draws) | 11 | 9 | 9 | 4 | 1 |
| GSE202518 | 6 vs 4 / 6 vs 6 / 12 vs 4 | 0.001099 (exact) | 44 | 27 | 0 | 0 | 27 |
| GSE22255 | 20 vs 20 | 9.999e-05 (10,000 draws) | 11 | 7 | 2 | 0 | 5 |
| GSE331114 | 4 vs 4 | 0.02857 (exact) | 40 | 14 | 0 | 0 | 14 |
| GSE58294 | 23 vs 23 | 9.999e-05 (10,000 draws) | 11 | 8 | 11 | 9 | 0 |
| GSE86941 | 3 vs 3 / 4 vs 4 | 0.02857 (exact) | 66 | 29 | 12 | 35 | 6 |
| All datasets | — | — | 183 | 94 | 34 | 48 | 53 |

Source: three_null_comparison_20261104.csv, built by 815 from the same per-sample scores as the main tables. Column 5 counts arms whose p comes from the expression-matched gene-set null reported in the paper; column 6 counts arms whose sample-label permutation also reaches 0.05, which in the small designs means reaching the smallest value the design can return. The two are not a before-and-after: a label permutation asks whether the group difference survives relabelling, a gene-set permutation asks whether the module moves more than random modules of the same size and expression level. Neither corrects the other. Column 8 counts the 53 arms where the gene-set p is below 0.05 and the label permutation is neither at its floor nor below 0.05 — significance with no sample-level counterpart. Of the 38 arms that clear both gates in the paper, 12 are in that column. They are not spread at random. 19 arms combine a double-gate pass with an absolute difference of at least 0.5, and 18 of those sit at the floor of their own design, meaning every relabelling gives a smaller difference than the one observed. The conflict column is therefore made of small effects and of the two designs that carry no surviving claim: GSE202518, whose control arm has four samples and whose CAPACITY difference returns −0.047 once composition is adjusted, and GSE331114, at four donors per compartment, where no arm clears three gates. It does not contain the large energy readings. The resolution column is the smallest value each dataset can return; within a dataset it differs by contrast, because a 12 vs 4 split has a finer grid than a 6 vs 4 one.What the table cannot do is adjudicate: in a 3 vs 3 design the label permutation cannot return a value below 0.10, so a real difference of moderate size and no difference at all are indistinguishable there. The gene-set null is the instrument that gives the paper its resolution and it is not confirmed at the sample level wherever groups are small; section 4.5 states what that constrains.

### Table S19. Coverage of the seven evidence layers, dataset by dataset

The 38 assembled datasets do not form one study. Each layer of the argument was tested in the datasets that could carry it, and no dataset carries all seven. The maximum is 5 layers, reached by 5 datasets (GSE58294, GSE171169, GSE332910, GSE225948, GSE86941); the median is 3. This table exists so that a reader can see which layers rest on the same samples and which are joined across series, rather than reconstructing that from ten sections of the text.

Layers. L1: the program-level 75% coverage gate was applied and at least one program was scored. L2: the primary contrast was tested against an expression-matched or size-matched null. L3: a leukocyte-lineage proxy adjustment or an Oaxaca-Blinder decomposition was performed. L4: the assay resolves cell types, so a score can be attributed to a compartment. L5: the series contains two or more post-injury time points. L6: the series was scored on the unified energy engine (Table 11). L7: the series extends to 14 days or beyond, or age enters as an explicit factor. Symbols: ● performed, with a product in the analysis ledger; ○ structurally possible and not performed; – structurally impossible, because bulk tissue cannot resolve cell types or the deposited matrices cannot be rebuilt; × scored and then withdrawn or downgraded (Table 1).

Counts by layer: L1 38, L2 29, L3 5, L4 15, L5 14, L6 6, L7 4. The three constraints the argument needs together — composition (L3), compartment (L4) and time (L5) — are never jointly testable in any dataset. This is not an omission: a series without cell resolution is the only place composition is a live question, and a sorted series has no composition to adjust for. The one dataset that decomposes composition inside a cell-resolved series is GSE250245, and it is single-time-point. Every layer rests on at least two independent datasets. No dataset carries more than 5 of the seven layers, so no single sample set demonstrates the composite argument end to end; section 3.5 states what follows from that, and section 2.4 states it for the carrier sequence in particular.

| **#** | **Dataset** | **Species** | **Assay** | **L1 Scored** | **L2 Null** | **L3 Composition** | **L4 Cell-resolved** | **L5 Time** | **L6 Energy** | **L7 Chronic/aged** | **Basis** |
| --- | --- | --- | --- | --- | --- | --- | --- | --- | --- | --- | --- |
| 1 | P1/P2 (in-house) | mouse | bulk RNA-seq | ● | ● | ○ | – | ○ | ○ | ○ | bulk tissue, no cell separation |
| 2 | Rat 28 d (in-house) | rat | bulk RNA-seq | ● | ● | ○ | – | ○ | ○ | ○ | bulk tissue, no cell separation |
| 3 | GSE250245 | rat | snRNA-seq | ● | ● | ● | ● | – | ○ | ○ | Oaxaca-Blinder on sorted cell types (Table 9); single time point by design |
| 4 | GSE16561 | human | microarray GPL6883 | ● | ● | ● | – | ○ | ● | ○ | five lineage proxies (Table 3); OB decomposition (Table 6); whole blood, no cell separation; on the unified energy engine (Table 11) |
| 5 | GSE58294 | human | microarray GPL570 | ● | ● | ● | – | ● | ● | ○ | five lineage proxies (Table 3); OB decomposition (Table 6); whole blood, no cell separation; on the unified energy engine (Table 11) |
| 6 | GSE22255 | human | Expression profile | ● | ● | ● | – | ○ | ● | ○ | five lineage proxies, negative control (Table 3); PBMC bulk, no cell separation; on the unified energy engine (Table 11) |
| 7 | GSE163654 | mouse+rat | microarray | ● | ● | ○ | – | ● | ○ | ○ | bulk tissue, no cell separation |
| 8 | GSE162955 | human | bulk GPL17586 | ● | ● | ○ | – | ○ | ○ | ○ | bulk tissue, no cell separation |
| 9 | GSE23160 | mouse | microarray | ● | ● | ○ | – | ● | ○ | ○ | bulk tissue, no cell separation |
| 10 | GSE233815 | mouse | Visium spatial | ● | ● | ○ | – | ● | ○ | ○ | Visium spots mix cell types by design |
| 11 | GSE171169 | mouse | scRNA-seq | ● | ● | – | ● | ● | ○ | ● |  |
| 12 | GSE227651 | mouse | scRNA-seq | ● | ● | – | ● | ● | ○ | ○ |  |
| 13 | GSE332910 | mouse | snRNA-seq | ● | ● | – | ● | ● | ○ | ● |  |
| 14 | GSE318800 | mouse | scRNA-seq | ● | ● | – | ● | ○ | ○ | ○ |  |
| 15 | GSE154396 | mouse | scRNA-seq | ● | ● | – | ● | ● | ○ | ○ |  |
| 16 | GSE345807 | rat | scRNA-seq | ● | ● | – | ● | ○ | ○ | ○ |  |
| 17 | GSE267023 | human | scRNA-seq (GEX only) | ● | ● | – | ● | ○ | ○ | ○ |  |
| 18 | GSE285659 | human | CITE-seq（GEX 36,601 + ADT 135） | ● | × | ○ | × | ○ | ○ | ○ | scored then withdrawn or downgraded (Table 1) |
| 19 | GSE337578 | mouse | SeekSpace spatial | ● | ○ | ○ | – | ○ | ○ | ○ | single section, no between-group contrast; single SeekSpace section, no cell-level annotation |
| 20 | GSE174574 | mouse | scRNA-seq | ● | ● | – | ● | ● | ○ | ○ |  |
| 21 | GSE37587 | human | microarray GPL6883 | ● | ● | ○ | – | ● | ○ | ○ | whole blood, no cell separation |
| 22 | GSE23651 | rat | microarray GPL6101 | ● | ● | ○ | – | ● | ○ | ○ | bulk tissue, no cell separation |
| 23 | GSE137482 | mouse | QuantSeq 3′ | ● | ● | ○ | – | ○ | ○ | ● | bulk tissue, no cell separation; 3 and 18 months, age as an explicit factor |
| 24 | GSE137595 | mouse | Multi-omics (transcriptome + proteome arms) | ● | ● | ○ | – | ○ | ○ | ○ | bulk tissue, no cell separation |
| 25 | GSE56267 | human | Deep bulk sequencing (GRCh38) | ● | × | ○ | – | ○ | ○ | ○ | bulk tissue, no cell separation; scored then withdrawn or downgraded (Table 1) |
| 26 | GSE142361 | mouse | bulk RNA-seq (raw count) | ● | × | ○ | – | ○ | ○ | × | bulk tissue, no cell separation; young versus aged, 14 d; scored then withdrawn or downgraded (Table 1) |
| 27 | GSE225948 | mouse | scRNA-seq (10x; brain CD45 mid/high + endothelium) | ● | ● | – | ● | ● | ○ | ● | young versus aged, d2 and d14 |
| 28 | GSE294912 | mouse | bulk RNA-seq (CAM + whole tissue) | ● | × | – | × | ○ | ○ | ○ | sorted CNS-associated macrophages alongside whole tissue; scored then withdrawn or downgraded (Table 1) |
| 29 | GSE105132 | mouse | Clariom array（oligo RMA） | ● | ● | – | ● | ○ | ○ | ○ | sorted microglia, macrophages and neutrophils |
| 30 | GSE141972 | mouse | bulk RNA-seq（featureCounts） | ● | ● | – | ● | ○ | ○ | ○ | sorted microglia and macrophages |
| 31 | GSE110141 | mouse | Agilent array (two platforms mixed) | ● | × | – | × | ○ | ○ | ○ | sorted microglia; scored then withdrawn or downgraded (Table 1) |
| 32 | GSE248793 | mouse | RSEM FPKM | ● | × | ○ | – | – | ○ | ○ | donor time after MCAO not disclosed (section 2.8); scored then withdrawn or downgraded (Table 1) |
| 33 | GSE64811 | mouse | Agilent GPL10787 (single platform) | ● | × | – | × | – | ○ | ○ | sorted resident microglia and invading macrophages (GFP+/RFP+); one time point (day 7); scored then withdrawn or downgraded (Table 1) |
| 34 | GSE86941 | rat | RSEM transcripts (21 samples) | ● | ● | – | ● | ● | ● | ○ | sorted microglia and macrophages (design field); on the unified energy engine (Table 11) |
| 35 | GSE331114 | human | scRNA-seq 10x Chromium 3′ v3 (CD45⁺ sorted) | ● | ● | – | ● | ○ | ● | ○ | on the unified energy engine (Table 11) |
| 36 | GSE204702 | human | scRNA-seq (67 h5; two CellRanger references) | ● | × | – | × | ○ | ○ | ○ | scored then withdrawn or downgraded (Table 1) |
| 37 | GSE202518 | human | bulk RNA-seq (whole blood, GPL24676 / NovaSeq 6000; NCBI | ● | ● | ● | – | ○ | ● | ○ | five lineage proxies, unidentified (R2 = 0.875; Table 3); whole blood, no cell separation; on the unified energy engine (Table 11) |
| 38 | GSE319238 | mouse | scRNA-seq | ● | ● | – | ● | ● | ○ | ○ |  |
